# Evidence for a bacterial Lands cycle phospholipase A: Structural and mechanistic insights into membrane phospholipid remodeling

**DOI:** 10.1101/2021.06.22.448587

**Authors:** Florian Bleffert, Joachim Granzin, Muttalip Caliskan, Stephan N. Schott-Verdugo, Meike Siebers, Björn Thiele, Laurence Rahme, Sebastian Felgner, Peter Dörmann, Holger Gohlke, Renu Batra-Safferling, Karl-Erich Jaeger, Filip Kovačić

**Author notes:** Corresponding authors:* Filip Kovacic, Renu Batra-Safferling, Holger Gohlke.

## Abstract

Cells steadily adapt their membrane glycerophospholipid (GPL) composition to changing environmental and developmental conditions. While the regulation of membrane homeostasis *via* GPL synthesis in bacteria has been studied in detail, the mechanisms underlying the controlled degradation of endogenous GPLs remain unknown. Thus far, the function of intracellular phospholipases A (PLAs) in GPL remodeling (Lands cycle) in bacteria is not clearly established. Here, we identified the first cytoplasmic membrane-bound phospholipase A_1_ (PlaF) from *Pseudomonas aeruginosa* involved in the Lands cycle. PlaF is an important virulence factor, as the *P. aeruginosa* Δ*plaF* mutant showed strongly attenuated virulence in *Galleria mellonella* and macrophages. We present a 2.0-Å-resolution crystal structure of PlaF, the first structure that reveals homodimerization of a single-pass transmembrane (TM) full-length protein. PlaF dimerization, mediated solely through the intermolecular interactions of TM and juxtamembrane regions, inhibits its activity. A dimerization site and the catalytic sites are linked by an intricate ligand-mediated interaction network which likely explains the product (fatty acid) feedback inhibition observed with the purified PlaF protein. We used molecular dynamics simulations and configurational free energy computations to suggest a model of PlaF activation through a coupled monomerization and tilting of the monomer in the membrane, which constrains the active site cavity into contact with the GPL substrates. Thus, these data show the importance of the GPL remodeling pathway for virulence and pave the way for the development of a novel therapeutic class of antibiotics targeting PlaF-mediated membrane GPL remodeling.

**Synopsis:** Membrane homeostasis can be regulated by phospholipase-controlled deacylation of endogenous glycerophospholipids (GPLs) followed by reacylation of products, known as the Lands cycle in eukaryotes. Here we show that the human pathogen *Pseudomonas aeruginosa* uses intracellular phospholipase A_1_ (PlaF) to modulate membrane GPL composition, which is the first example in bacteria. This newly identified PLA_1_ indirectly regulates the bacterial virulence properties by hydrolyzing a specific set of membrane GPLs. The crystal structure of full-length PlaF dimers bound to natural ligands, MD simulations, and biochemical approaches provide insights into the molecular mechanism of dimerization-mediated inactivation of this single-pass transmembrane PLA_1_. Our findings shed light on a mechanism by which bacterial intracellular PLAs might regulate membrane homeostasis what showcases these enzymes as a promising target for a new class of antibiotics.

## Introduction

Biological membranes are steadily changing and adapting to environmental and developmental conditions [1, 2]. These changes affect processes indispensable for cell life, such as nutrient uptake [3], chemical signaling [4], protein secretion [5], folding [6], interaction with hosts [7], and antibiotic resistance [8]. An important mechanism to maintain membrane functionality in bacteria is the alteration of lipid composition [9–11]. The adjustment of the fatty acid (FA) composition of glycerophospholipids (GPL) upon thermal adaptation represents one of the most important mechanisms of membrane lipid homeostasis [12, 13]. Adaptive changes in membrane GPL composition were observed under numerous other conditions, including environmental stresses [9], the transition from planktonic to sessile lifestyle [14], and heterologous protein production [15].

*De novo* synthesis of GPLs is the main pathway used to tune the proportions of different lipid classes in bacteria [11, 16]. Furthermore, bacteria rapidly alter their membrane GPL composition by chemical modifications (cis-trans isomerization and cyclopropanation) of acyl chains in GPLs as a response to environmental changes [11]. However, the bacterial pathway for remodeling of GPLs involving a rapid turnover of the acyl chains of GPLs is unknown. Interestingly, such a pathway was discovered in eukaryotes by W. E. Lands more than sixty years ago [17]. This Lands cycle involves PLA-catalyzed deacylation of membrane GPLs to mono-acyl GPLs (lysoGPLs) followed by lysophospholipid acyltransferase (LPLAT)-mediated acylation of lysoGPL to yield a new GPL molecule with acyl chain composition different from the starting GPL [17]. Despite the importance of this metabolic process in different animal and plant tissues, it took nearly fifty years before the enzymes involved in phospholipid remodeling have been discovered [18]. Three mammalian LPLAT with different substrate specificities were reported to be involved in the Lands cycle [19]. At least six mammalian PLAs (α, β, γ, δ, ε, ζ) that act on the membrane GPLs with different substrate profiles and tissue expression patterns are known [20–23]. PLAγ is the only one with a suggested role in the remodeling of membrane GPLs [24], while other PLAs are involved in the production of lipid mediators and bioenergetics [25].

Whereas extensive studies have been carried out for secreted bacterial PLAs acting as host-cell effectors [26], only limited information is available for the enzymes from the intracellular PLA family [27]. Previously, we reported that periplasmic TesA from *P. aeruginosa* is a multifunctional enzyme with lysoPLA activity [28]. However, this enzyme has no PLA activity, and therefore it is most likely not related to membrane GPL remodeling [29]. Comprehensive lipidomic profiling of 113 *E. coli* strains with deleted or overexpressed lipid metabolism genes did not reveal the identity of an intracellular PLA suitable for the Lands cycle [16]. Here, we describe PlaF from *P. aeruginosa* [30, 31] as the first cytoplasmic membrane-bound PLA with a role in virulence and GPL remodeling pathway in bacteria. We determined the X-ray crystal structure of PlaF [30, 31] as a basis to provide mechanistic insights into PLA-mediated membrane phospholipid degradation related to bacterial virulence.

## Results

### PlaF is an integral cytoplasmic membrane-bound enzyme

We previously purified PlaF from the Triton X-100 solubilized membranes of a *P. aeruginosa* strain carrying the *p-plaF* expression plasmid [30, 31]. Here, we show that catalytically active PlaF is an intrinsic integral membrane protein as it was absent in the soluble fraction of the *P. aeruginosa* p-*plaF* (Fig. 1a) and it remained membrane-associated after treatment of PlaF-containing membranes with denaturation agents (Na_2_CO_3_ or urea), which destabilize weak interactions between peripheral proteins and the membrane (Fig. 1b). To identify if PlaF is associated with the inner or outer membrane, *P. aeruginosa p-plaF* membranes were fractioned by ultracentrifugation in a sucrose density gradient. Western blot analysis of the cytoplasmic membrane protein SecG [32], and the outer membrane-associated Lipid A [33] combined with PlaF activity measurement revealed that the majority of PlaF was in the cytoplasmic membrane fractions (# 9-13) (Fig. 1c). As expected, the Lipid-A-containing fractions (# 1-3) showed negligible PlaF activity (Fig. 1c), overall demonstrating that PlaF is a cytoplasmic integral membrane protein. Proteolysis experiments in which *P. aeruginosa p-plaF* cells with a chemically permeabilized outer membrane were treated with trypsin revealed a time-dependent degradation of PlaF (Fig. 1d). These results suggest that PlaF is likely anchored to the cytoplasmic membrane *via* a TM domain at the N-terminus which was predicted from the sequence analysis [30], and its catalytic C-terminal domain protrudes into the periplasm.

**Fig. 1.**
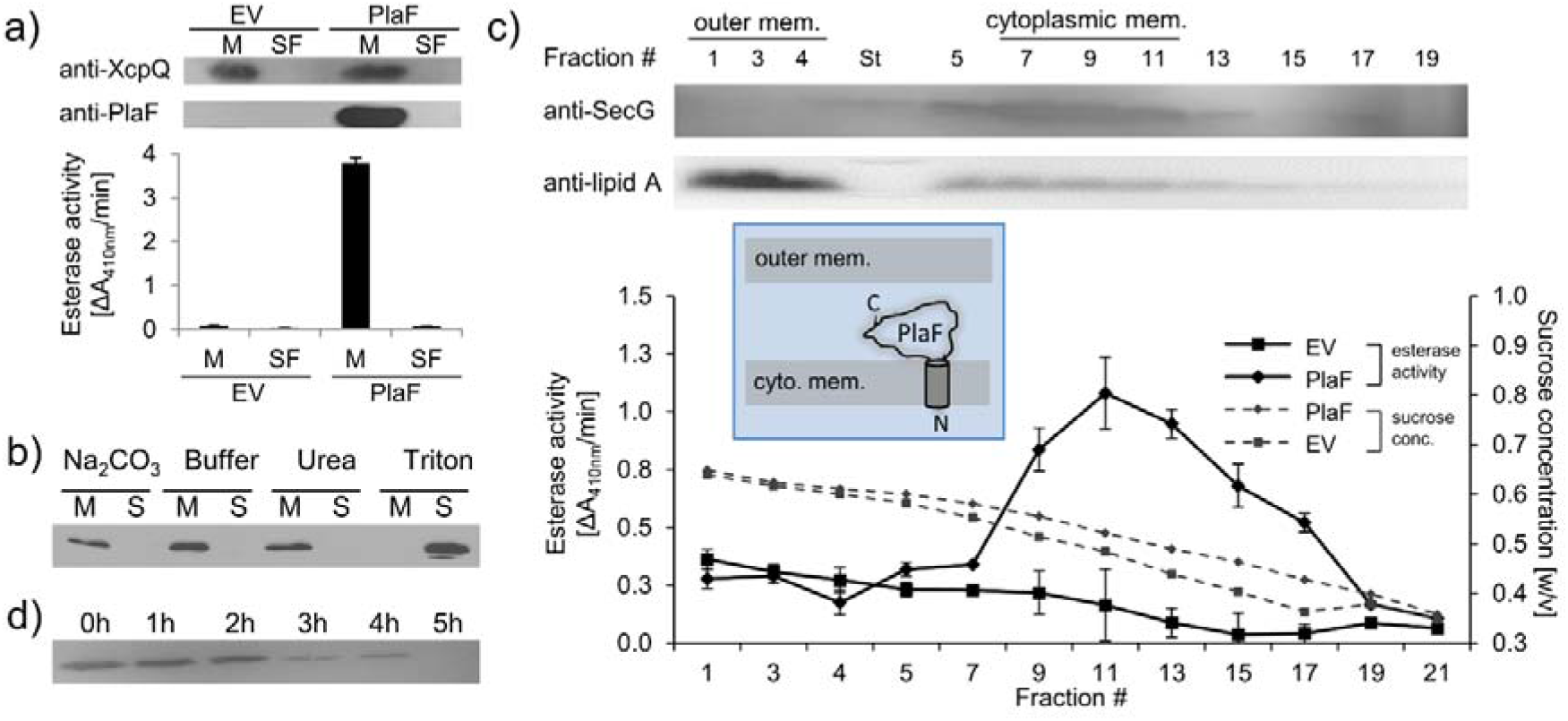
Subcellular localization of PlaF. **a) PlaF is a membrane protein of *P. aeruginosa***. The membrane (M), and soluble fractions (SF) of cell extracts from *P. aeruginosa p-plaF*, and the empty vector control strain (EV) were separated, analyzed by immunodetection with anti-His_6_-tag antibodies, and by esterase activity assay. The membrane protein marker *P. aeruginosa* XcpQ was detected with anti-XcpQ antibodies. **b) PlaF is an integral membrane protein of *P. aeruginosa***. The crude membranes of *P. aeruginosa p-plaF* were treated with sodium carbonate, urea, Triton X-100, or MES buffer control followed by ultracentrifugation (S, supernatant; M, membrane proteins). PlaF was detected as in panel a. **c) PlaF is a cytoplasmic-membrane protein of *P. aeruginosa***. The membrane fractions of *P. aeruginosa p-plaF*, and the EV strains were isolated, and separated by ultracentrifugation in a sucrose density gradient. The esterase activity was assayed as in panel a. *P. aeruginosa* SecG, and outer membrane lipid A were used as markers for cytoplasmic, and outer membranes, and detected by Western blotting using anti-SecG, and anti-Lipid A antibodies, respectively. Inlet: A model of PlaF cellular localization. All values are mean ± standard deviation (S.D.) of three independent experiments measured in triplicates. **d) The catalytic domain of PlaF is exposed to the periplasm.***P. aeruginosa p-plaF* cells with permeabilized outer membrane were treated with trypsin for the indicated periods, and PlaF was detected as described in figure 1a.

### PlaF is a PLA_1_ involved in the alteration of membrane GPL composition as determined by global lipidomics

The previously reported carboxylesterase activity of PlaF [31] was here further analyzed using different PLA substrates. PlaF purified with *n*-octyl-β-d-glucoside (OG) as described previously [30], showed PLA_1_ but no PLA_2_ activity towards the artificial substrates specific to each of these two phospholipase families (Fig. 2a). The substrate profile of PlaF against natural di-acyl GPLs commonly occurring in *P. aeruginosa* membranes [14] was determined with a spectrum of substrates (see legend to Fig. 2b). *In vitro*, purified PlaF preferably hydrolyzed GPLs containing medium-chain FAs (C12, C14), and showed comparable activities with phosphatidylethanolamine (PE), phosphatidylglycerol (PG), and phosphatidylcholine (PC) (Fig. 2b).

**Fig. 2.**
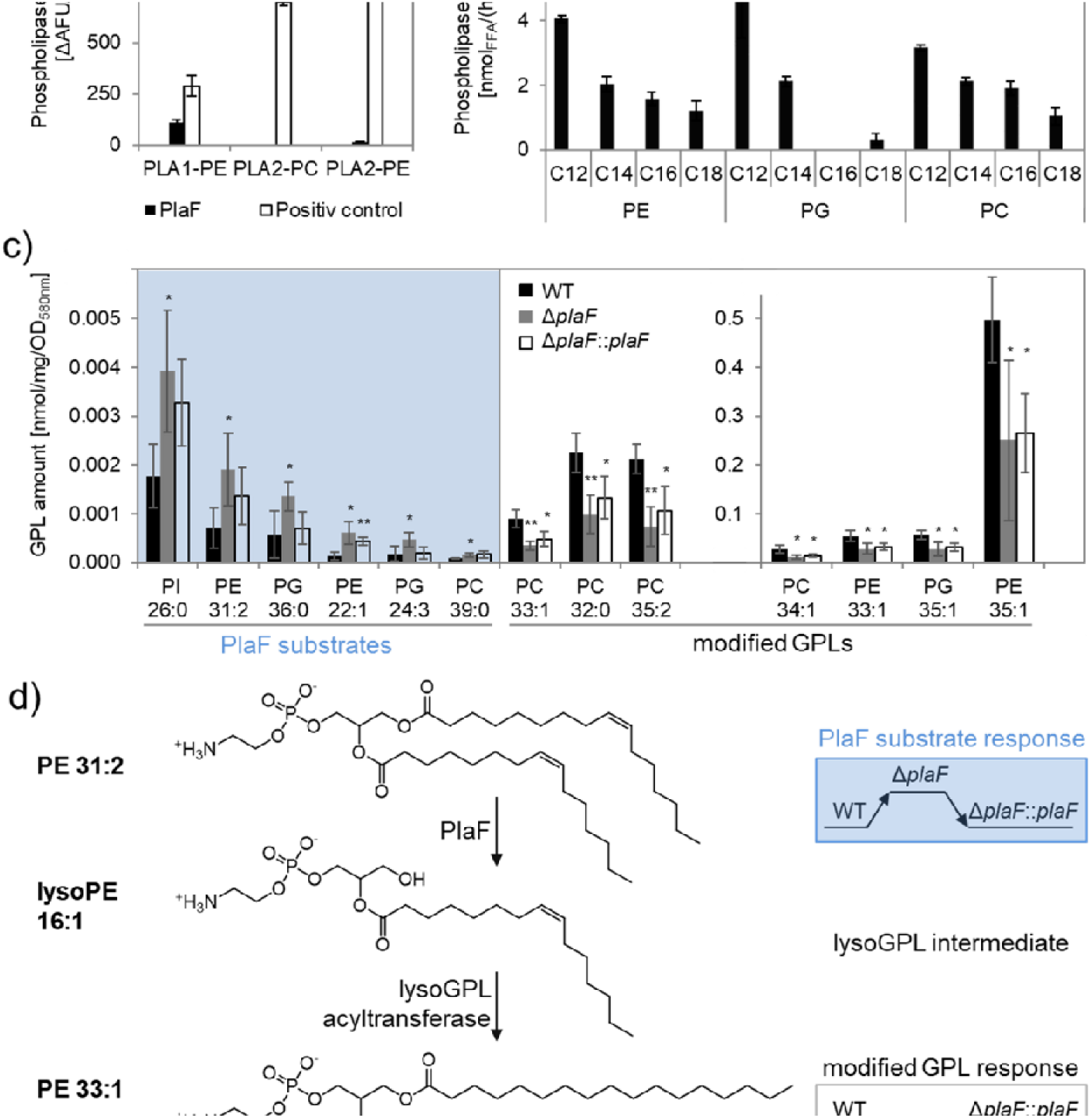
Phospholipolytic activity profiling of PlaF. **a) PlaF is a phospholipase A_1_.** Enzyme activities of PlaF were measured fluorimetrically using artificial PLA_1_, and PLA_2_ substrates containing either ethanolamine (PE) or choline (PC) head groups. The control enzymes were PLA_1_ of *T. lanuginosus, and* PLA_2_ of *N. mocambique*. Results are means ± S.D. of three independent measurements performed with at least three samples. **b) PlaF releases FAs from naturally occurring bacterial GPLs.** PLA activity of PlaF was measured by quantification of released FAs after incubation of PE, PG, and PC substrates containing FAs with different chain lengths (C12-C18). **c) PlaF changes GPL composition of *P. aeruginosa* cells.** Crude lipids extracted from *P. aeruginosa* wild-type, Δ*plaF*, and Δ*plaF::plaF* membranes were quantified by Q-TOF-MS/MS using an internal standard mixture of GPLs. The GPL amount (nmol) was normalized to mg of crude lipids, and optical density (Table S3). FA composition of GPL is depicted as XX:Y, where XX defines the number of carbon atoms, and Y defines the number of double bonds in FAs bound to GPL. Results are mean ± S.D. of four biological replicates of wildtype, Δ*plaF, and* three of the Δ*plaF::plaF*. T-test of normally distributed values, ** p < 0.01, * p < 0.05. **d) A putative Lands cycle of *P. aeruginosa* involving PlaF explained on the example of conversion of PE 31:2 to PE 35:1.** GPLs which concentrations show PlaF substrate response (elevation in Δ*plaF* and depleted by expressing *plaF* in Δ*plaF*) will be hydrolyzed in *sn*1 position to yield lysoGPL intermediate which will be acylated by still unknown lysoGPL acyltransferase to yield modified GPLs. Concentrations of modified GPLs show inverse response as GPL substrates.

To examine the role of membrane-bound PlaF in the regulation of the membrane GPL composition *in vivo, we* constructed the *P. aeruginosa* deletion mutant Δ*plaF* lacking the entire *plaF* gene by homologous recombination (Fig. S1a-c), and a complemented Δ*plaF::plaF* strain as a control (Fig. S1d). The activity assay showed ^~^90 % loss of PLA_1_ activity in the mutant strain, and restoration of activity in Δ*plaF::plaF* slightly above the wild-type level (Fig. S1e). These findings indicat that PlaF is a major but not the only intracellular PLA_1_ in *P. aeruginosa*.

The quantitative mass spectrometric (Q-TOF-MS/MS) analysis of total GPLs isolated from four biological replicates of *P. aeruginosa* wild-type, Δ*plaF*, and Δ*plaF::plaF* cells revealed significant differences in membrane GPL composition (Fig. 2c, Tables S1-3). Statistical analysis of 323 GPL molecular species identified six significantly (*p* < 0.05) accumulating GPLs, varying in the composition of head groups (PE, PG, PC, and phosphatidylinositol, PI), length, and unsaturation of acyl chains, in *P. aeruginosa* Δ*plaF*. Interestingly, these GPLs were present at low concentrations in the cells what may explain why they were not detected in the previous lipidomic analyses of *P. aeruginosa* GPLs [14, 34]. In the complemented strain (Δ*plaF::plaF*) these GPLs were depleted compared to the Δ*plaF*, although not to the level of the wild-type (Table S2). These results strongly indicate that PlaF specifically hydrolyses low abundant GPLs *in vivo*. We furthermore observed that the other seven PE, PG, and PC species, which belong among the most abundant *P. aeruginosa* GPLs [14, 34], were significantly depleted (Fig. 2c) in *P. aeruginosa* Δ*plaF*, and their concentrations were significantly elevated in complementation strain (Fig. 2c). This may explain why the net GPL contents of the wild-type and the Δ*plaF* strain were not significantly (*p* = 0.67) different. Significantly affected GPLs in the Δ*plaF* strain account for ^~^11 % (mol/mol) of the total GPL content, indicating the profound function of PlaF in membrane GPL remodeling.

Our quantitative lipidomics results which revealed that several PE, PG, and PC molecular species accumulated or were depleted in Δ*plaF*, together with *in vitro* PLA activity data of PlaF with various PE, PG, and PC substrates, strongly indicate that PlaF might be a major PLA involved in the Lands cycle (Fig. 2d). Thus, six low abundant PE, PG, and PC species which accumulated in Δ*plaF* might represent the substrates of PlaF. PlaF-mediated hydrolysis of these substrates yields lysoGPL intermediates. Acylation of these lysoGPLs by an unknown acyltransferase, will produce modified GPLs typical to *P. aeruginosa*. The absence of lysoGPL intermediates in Δ*plaF* will lead to the depletion of modified GPLs (Fig. 2d).

### *PlaF is a potent virulence factor of* P. aeruginosa *affecting in vivo toxicology*

We next addressed the question of whether PlaF contributes to the virulence of *P. aeruginosa* by using the *Galleria mellonella* infection model and the bone marrow-derived macrophages (BMDMs) viability assay. The results revealed a remarkable difference in the survival of *G. mellonella* larvae infected with *P. aeruginosa* wild-type or Δ*plaF*. While Δ*plaF* was avirulent during 20 h of infection, the majority of the larvae (^~^80 %) did not survive 20 h after infection with the *P. aeruginosa* wild-type (Fig. 3a). The viability assays with *P. aeruginosa* infected BMDMs showed a significantly (*p* < 0.01) stronger killing effect of *P. aeruginosa* wild-type compared to Δ*plaF* 6 h after infection (Fig. 3b). As expected, the complemented strain (Δ*plaF::plaF*) restored the loss of virulence of Δ*plaF* in *G. mellonella, and* BMDM assays (Figs. 3a, and 3b). Comparison of the growth curves of *P. aeruginosa* Δ*plaF*, and the wildtype in nutrient-rich medium (Fig. S2a) showed that PlaF most likely does not reduce the virulence by affecting the growth of *P. aeruginosa*.

**Fig. 3.**
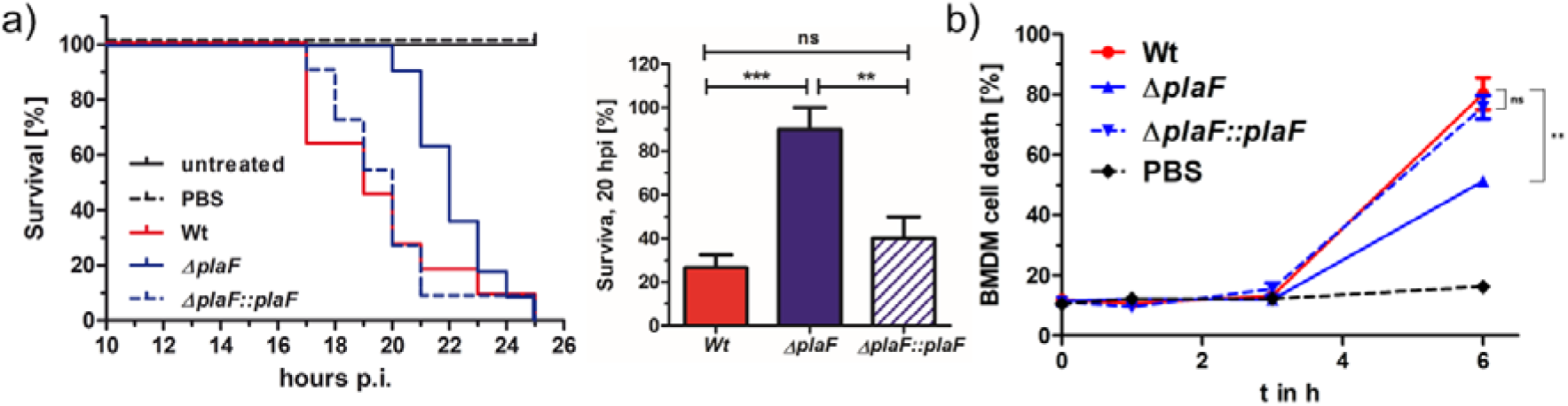
*PlaF is a novel virulence factor of* P. aeruginosa *PAO1*. **a) Left: *P. aeruginosa* Δ*plaF* strain is less virulent than the respective wild-type strain in a *G. mellonella* larvae virulence assay.** Kaplan-Meier plot of representative data of at least two experiments with 10 larvae per group. PBS treated, and untreated larvae served as infection, and viability controls, respectively. Right: Statistical analysis of the survival at the 20 h using three independent experiments with 10 larvae each. b) ***P. aeruginosa* Δ*plaF* strain is less cytotoxic to bone marrow-derived macrophages (BMDMs) than the wild-type strain in cell culture.** The BMDM cells (5×10^5^) were infected with 5×10^5^ bacteria in a 24-well plate, and lactate dehydrogenase activity in supernatants were determined as a measure of BMDM death. The Δ*plaF* phenotype could be complemented with *P. aeruginosa* Δ*plaF::plaF*. PBS or Triton-X100 (1 % v/v) treated cells served as viability or 100% killing controls, respectively. Results are the representative data of 2 independent experiments (n = 10). One-way Anova analysis, *** p < 0.001, ** p < 0.01, ns = not significant.

A BLAST search revealed PlaF orthologs in more than 90% of all sequenced *P. aeruginosa* genomes, including 571 clinical isolates (Table S4). Furthermore, we found PlaF homologs in pathogens from the *Pseudomonas* genus (*P. alcaligenes, P. mendocina, and P. otitidis*), and other high-priority pathogens (*Acinetobarter baumannii, Klebsiella pneumoniae, and Streptococcus pneumoniae*) (Fig. S3). These results indicate that PlaF is a novel, and very potent *P. aeruginosa* virulence factor, which is conserved in important pathogens, and therefore represents a promising target for development of novel broad-range antibiotics.

### Crystal structure of PlaF homodimer in the complex with natural ligands

To gain insights into the PlaF structure, we crystallized the OG-solubilized PlaF protein purified from *P. aeruginosa* membranes as described previously [31]. The structure was refined at a resolution of up to 2.0 Å (Table 1). The final model in the asymmetric unit consists of two protein molecules (PlaF_A_ and PlaF_B_), which are related by improper 2-fold non-crystallographic symmetry (Fig. 4a). Active site cavities of both the monomers reveal non-covalently bound ligands - myristic acid (MYR), OG, and isopropyl alcohol (IPA) in PlaF_A_; and undecyclic acid (11A), OG, and IPA in PlaF_B_ (Fig. 4a, Table S5). These FAs are the natural ligands from the homologous organism *P. aeruginosa* that were co-purified with PlaF, as confirmed by GC-MS analysis of organic solvent extracts of purified PlaF (Fig. S4).

**Table 1:**
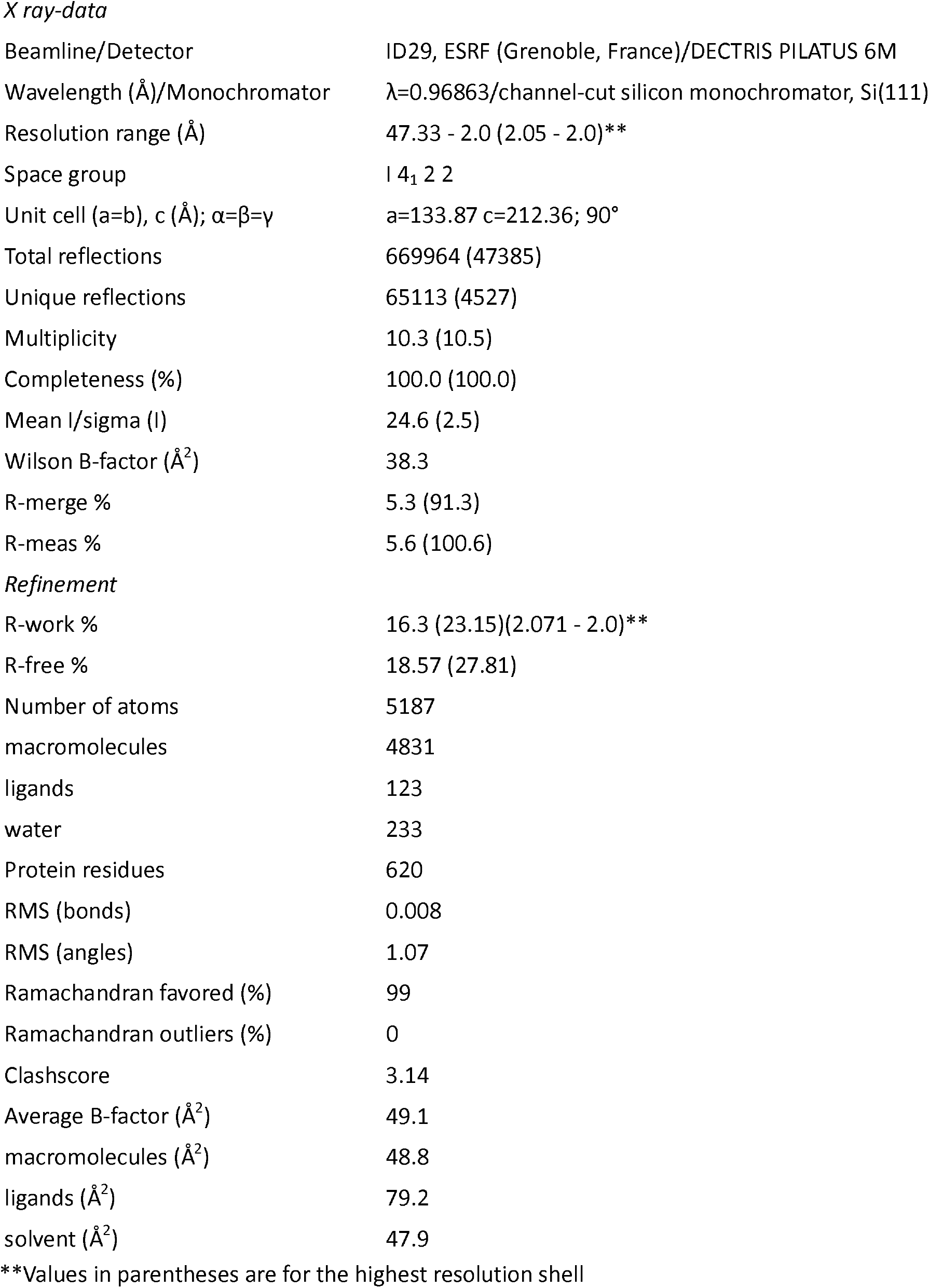
Data collection, and refinement statistics on PlaF

*X ray-data*
|  |  |
| --- | --- |
| Beamline/Detector | ID29, ESRF (Grenoble, France)/DECTRIS PILATUS 6M |
| Wavelength (Å)/Monochromator | $\lambda=0.96863$/channel-cut silicon monochromator, Si(111) |
| Resolution range (Å) | 47.33 - 2.0 (2.05 - 2.0)** |
| Space group | I 4<sub>1</sub> 2 2 |
| Unit cell (a=b), c (Å); $\alpha=\beta=\gamma$ | a=133.87 c=212.36; 90° |
| Total reflections | 669964 (47385) |
| Unique reflections | 65113 (4527) |
| Multiplicity | 10.3 (10.5) |
| Completeness (%) | 100.0 (100.0) |
| Mean I/sigma (I) | 24.6 (2.5) |
| Wilson B-factor (Å<sup>2</sup>) | 38.3 |
| R-merge % | 5.3 (91.3) |
| R-meas % | 5.6 (100.6) |
| *Refinement* |  |
| R-work % | 16.3 (23.15)(2.071 - 2.0)** |
| R-free % | 18.57 (27.81) |
| Number of atoms | 5187 |
| macromolecules | 4831 |
| ligands | 123 |
| water | 233 |
| Protein residues | 620 |
| RMS (bonds) | 0.008 |
| RMS (angles) | 1.07 |
| Ramachandran favored (%) | 99 |
| Ramachandran outliers (%) | 0 |
| Clashscore | 3.14 |
| Average B-factor (Å<sup>2</sup>) | 49.1 |
| macromolecules (Å<sup>2</sup>) | 48.8 |
| ligands (Å<sup>2</sup>) | 79.2 |
| solvent (Å<sup>2</sup>) | 47.9 |

\*\*Values in parentheses are for the highest resolution shell

**Fig. 4.**
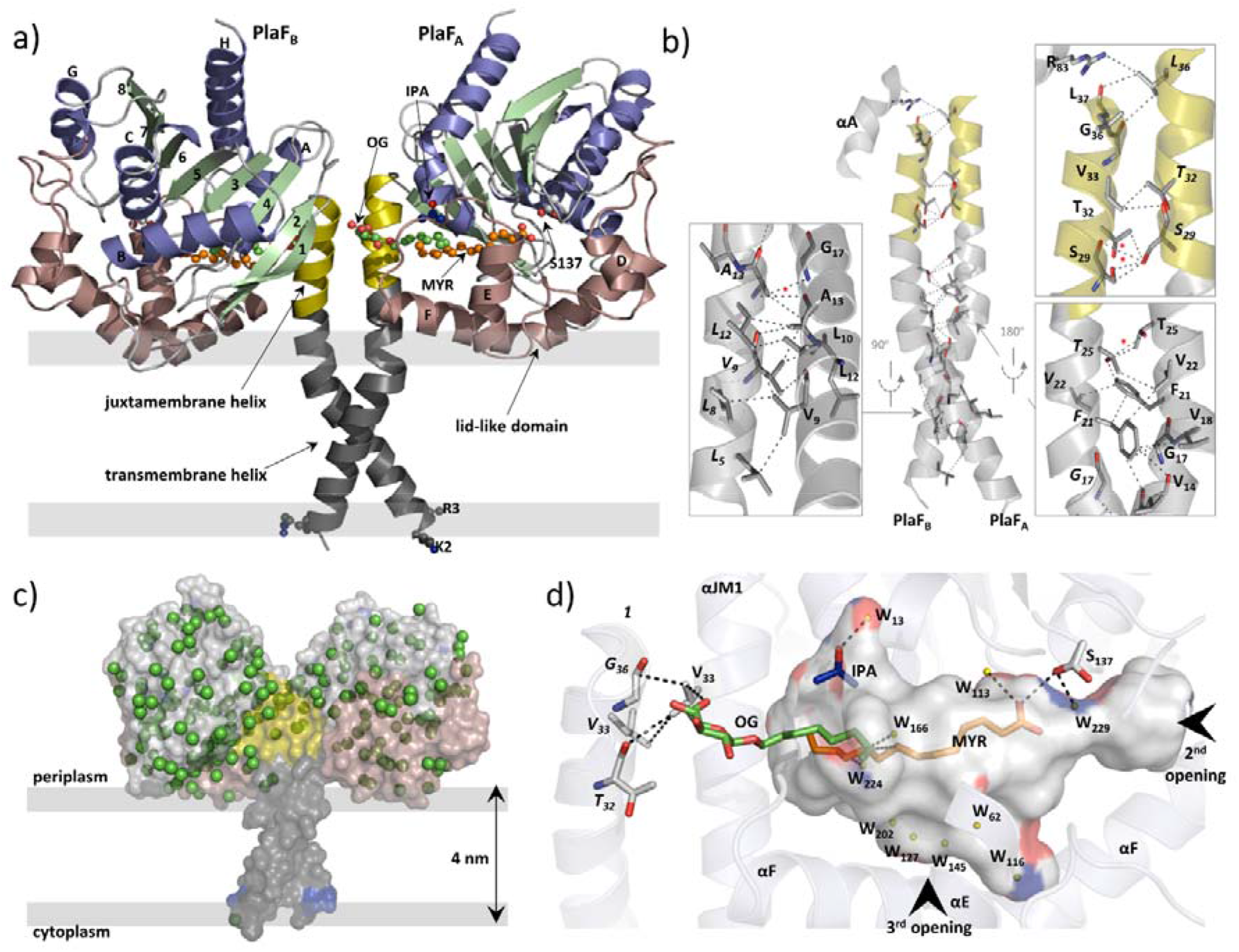
Overall structure of dimeric PlaF with bound endogenous FA ligands. **a)** A unique N-terminal helix comprising a putative transmembrane helix (αTM1, L_5_ - L_27_, grey) flanked by charged residues (K_2_, R_3_) on one side and, on another side, the juxtamembrane helix (αJM1, A_28_ - L_37_, yellow). αJM1 links the αTM1 with the catalytic domain which consists of an α/β-hydrolase (blue, α-helices; green, β-strands, and grey, loops), and a lid-like domain (brown). Ligands bound in the active site cleft are shown as ball-and-sticks (oxygen, red; carbon of OG, MYR, and IPA, green, orange, and blue, respectively). Thick grey lines roughly depict the membrane borders. **b) Dimer interface.** Interactions involving TM-JM helices are predominantly hydrophobic with four weak H-bonds (indicated by a red asterisk) detected mostly in the αJM1. R_83_ is the only residue outside of the JM-TM helix involved in interactions. Residues of the PlaF_B_ molecule are indicated in italics. A detailed list of interactions is provided in table S6. **c) A model suggesting the orientation of PlaF in the membrane.** The water molecules are indicated as green spheres. The transparent surface of PlaF was colored as in Fig. 4a. PlaF is rotated by 180<u>°</u> along the normal to the membrane compared with Fig. 4a. **d) Interaction network within the ligand-binding cleft of PlaF_A_.** MYR is linked *via* H-bond with the catalytic S_137_, and *via* hydrophobic interactions with OG. The sugar moiety of OG from PlaF_A_ forms H-bonds with V_33_ of PlaF_A_ which is interacting with V_33_, and G_36_ of PlaF_B_. The part of the cleft in direction of the opening 3 is occupied by several water molecules (W, yellow spheres). The cleft accommodates one IPA molecule bound to the water. Arrows indicate two openings not visible in this orientation. The cleft was calculated using the Pymol software and colored by elements: carbon, gray; oxygen, red; nitrogen, blue.

The N-terminal 38 amino acids form a long, kinked helix that comprises the putative TM (αTM1) and the JM (αJM1) helices, connecting the catalytic domain with the membrane (Fig. 5a). The kink angle in the TM-JM helices is the main difference between two monomers (Fig. S5a) and is likely caused by crystal packing effects (Fig. S5b). Dimerization is mediated primarily *via* hydrophobic interactions between symmetry-nonrelated residues from the TM-JM domains of two monomers (Fig. 4b, Table S6), consistent with the hydrophobic effects that dominate in the stabilization of dimeric TM domains [35]. In addition, four weak H-bonds (Fig. 4b) between JM residues stabilize the PlaF dimer. The TM-JM helices adopt a coiled-coil-like conformation (Fig. S5c), where the αTM1 crosses its counterpart at V14 to form an elongated X-shaped dimer interface with the buried surface area of 656 Å^2^ per monomer. PlaF dimerisation interface is slightly larger than the interface of glycophorin TM helix dimer (400 Å^2^) which is missing the JM region [35], however, it is relatively small compared with most protein-protein complexes (>1000 Å^2^) [36]. The full-length PlaF dimer represents a unique structure, as neither a relevant match to the TM-JM helix (Fig. S6) nor the membrane-spanning coiled-coil structure of the TM-JM dimer have been reported previously.

**Fig. 5.**
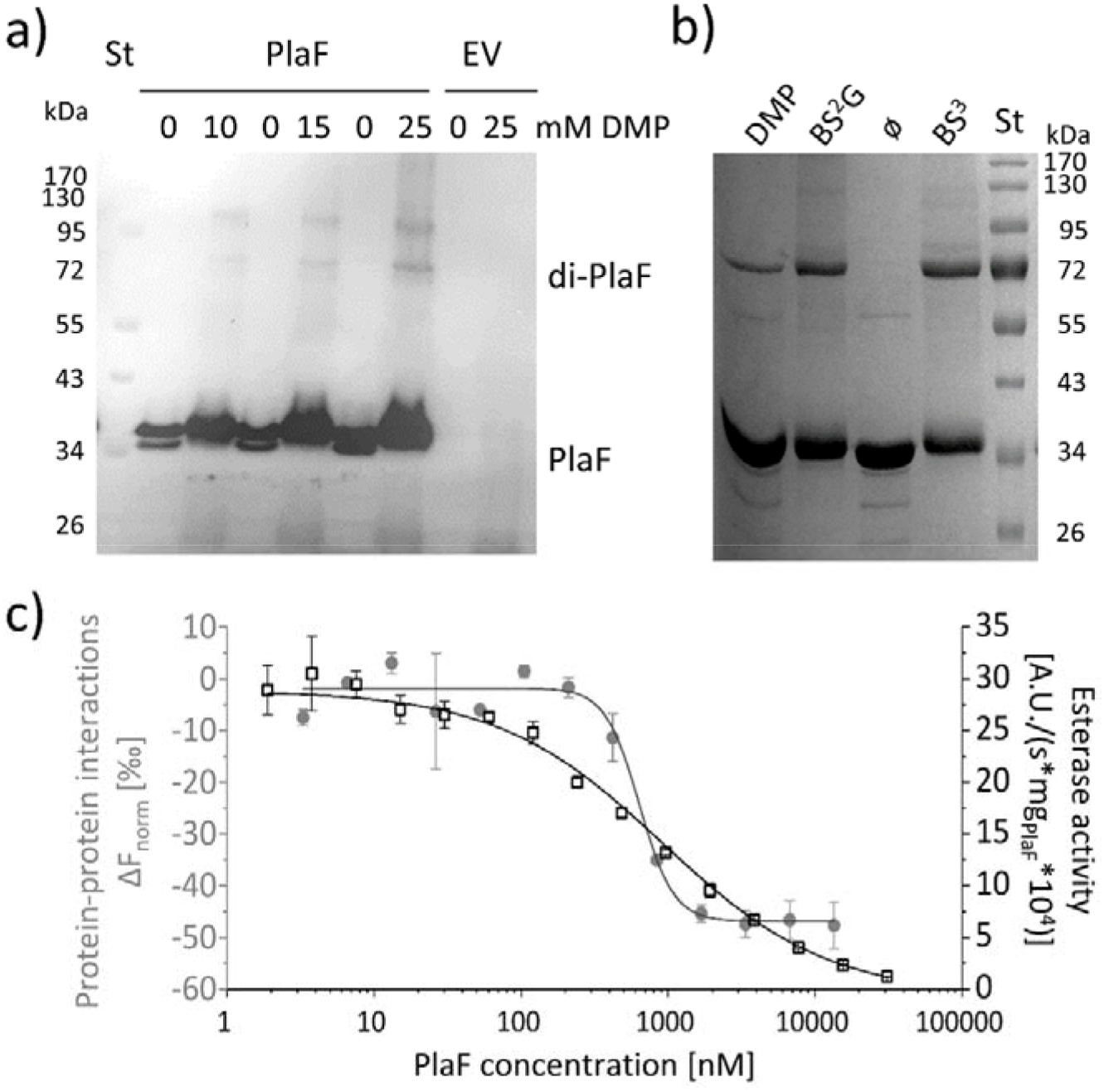
PlaF oligomeric states, and their enzymatic activity. **a) PlaF forms dimers in cell membranes.** *In vivo* cross-linking experiments were performed by incubating *P. aeruginosa* p-*plaF* or the empty vector control (EV) cells with different concentrations of DMP cross-linker followed by immunodetection of PlaF with anti-PlaF antiserum. **b) *In vitro* cross-linking of purified PlaF.** Purified PlaF was incubated with DMP, BS^2^G, and BS^3^ cross-linking reagents or buffer control (ø) for 90 min, and the samples were analyzed by SDS-PAGE. Molecular weights of protein standard in kDa are indicated. **c) PlaF homodimerization, and activity are concentration-dependent.** Protein-protein interactions of purified PlaF were monitored by measuring the changes in thermophoresis (Δ*F*_norm_, grey circles) using the MST method. The MST results are mean ± S.D. of two independent experiments with PlaF purified with OG. Esterase activity (black squares) of PlaF was measured in three independent experiments using 4-methylumbelliferyl palmitate substrate. Dissociation (*K*_D_), and activation (*K*_act_) constants were calculated using a logistic fit of sigmoidal curves.

### The crystal structure of PlaF is indicative of a specific orientation in the membrane

The catalytic domain of PlaF adopts a canonical α/β-hydrolase fold [37] (Fig. 4a) with three α- helices forming a distinct lid-like domain that covers the active site (Fig. 4a). Despite the high homology of the catalytic domain, the lid-like domain varies significantly between PlaF homologs (Fig. S7), as observed previously for other lipolytic enzymes (Fig. S7) [38]. Furthermore, the lid-like domain shows a less ordered structure, as suggested by comparatively higher B-factors (Fig. S8a). This is likely a consequence of the lack of stabilizing interactions between the charged residue-rich (23 of 77 residues) lid-like domain and the hydrophilic head groups of membrane GPLs in the native membrane environment. The TLS (translation-libration-screw-rotation) model revealed higher disorder in the TM-JM domains, presumably also due to the missing interactions with the hydrophobic membrane (Fig. S8b). No ordered water molecules in the vicinity of αTM1 (Fig. 4c), and the presence of several charged and polar residues adjacent to αTM1 suggest a model where the TM-JM domain spans through the membrane with the catalytic domain localized on the membrane surface (Fig.4c).

### Ligand-mediated interaction network connects dimerization and active sites

The active site of PlaF comprises the typical serine-hydrolase catalytic triad with S_137_, D_258_, and H_286_ interacting through H-bonds [39] (Table S7). Interestingly, S_137_ shows two side-chain conformations, where one conformer is within the hydrogen bond distance of FA ligand (Fig. 4d, Tables S5 and S7). Additionally, S137 forms H-bonds with residues I_160_, D_161_, and A_163_ located in the lid-like domain. The active site cleft in PlaF is formed by residues from the helix αJM1, the α/β-hydrolase and the lid-like domains (Fig. 4d, Table S8). In PlaF, the large T-shaped active site cleft formed by residues from the JM helix, the α/β-hydrolase, and the lid-like domains is amphiphilic in nature, making it compatible to bind the bulky GPL substrates. Three openings are observed in the cleft - one, close to the catalytic S_137_, lined with residues from the loops preceding αE, and αF; second, in the middle pointing towards the putative membrane, lined mostly with polar residues of the loops preceding αB, and αF; and third, at the dimer interface, comprising residues from αJM1, and the loop preceding αF of the lid-like domain. The third opening accommodates an OG molecule (Fig. 4d), with the pyranose ring of OG interacting with residue V_33_ of PlaF_A_, which in turn participates in dimerization *via* interactions with V_33_ and T_32_ of PlaF_B_ (Fig. 4b). The alkyl chains of OG and MYR bound in the active site cleft are stabilized *via* hydrophobic interactions (Fig. 4d). Finally, the H-bond interaction of catalytic S_137_ with the carboxyl group of MYR completes an intricate ligand-mediated interaction network bridging the catalytic (S_137_) and dimerization (V_33_) sites in PlaF (Fig. 4d). The crystal structure presented thus suggests a role of dimerization and ligand binding in the regulation of the PlaF function, which was subsequently analyzed biochemically.

### The PlaF activity is affected by dimerization

To investigate the oligomeric state of PlaF *in vivo*, we performed cross-linking experiments in which intact *P. aeruginosa* p-*plaF* cells were incubated with the cell-permeable bi-functional cross-linking reagent dimethyl pimelimidate (DMP). Western blot results revealed the presence of monomeric and dimeric PlaF in DMP-treated cells, whereas dimers were absent in untreated cells (Figs. 5a, and S9). Size exclusion chromatography showed that PlaF was extracted from the membranes with detergent and purified by IMAC elutes as a monomer (Fig. S10). Incubation of this purified PlaF for 90 min with bi-functional cross-linkers of different lengths (DMP; bis(sulfosuccinimidyl) glutarate, BS^2^G or bis(sulfosuccinimidyl) suberate, BS^3^) resulted in the formation of a substantial amount of PlaF dimers, suggesting spontaneous dimerization in the solution (Fig. 5b). Microscale thermophoresis (MST) measurements were performed in which the fluorescence-labeled PlaF was titrated with an equimolar concentration of non-labeled PlaF to quantify spontaneous dimerization. The results revealed a sigmoidal binding curve from which was calculated a dissociation constant *K*_D_ = 637.9 ± 109.4 nM indicating weak binding, (Fig. 5c). Measurements of the esterase activity of PlaF samples used for MST experiments revealed that the specific activity of PlaF strongly decreased with increasing PlaF concentrations (Fig. 5c). Enzyme activity measurements were employed to calculate the activation constant *K*_act_ = 916.9 ± 72.4 nM. The similar dissociation and activation constants support a model in which PlaF activity is regulated through reversible dimerization *in vitro*.

### Fatty acids induce dimerization and inhibit PlaF

To investigate the effect of FA ligands on the activity of PlaF, we used mM concentrations of FAs with different chain lengths (C5 – C15) in a competitive inhibition assay. PlaF was strongly inhibited (>80%) with FAs containing 10 to 14 carbon atoms (Fig. 6a), while the shorter and longer FAs showed only moderate to weak inhibition (Fig. 6a). To explore the underlining mechanism, we performed kinetic inhibition studies with increasing concentrations of decanoic acid (C10). The results showed that C10 FA lowered maximal hydrolysis rates (*v*_max_) as expected for a competitive inhibitor.

**Fig. 6.**
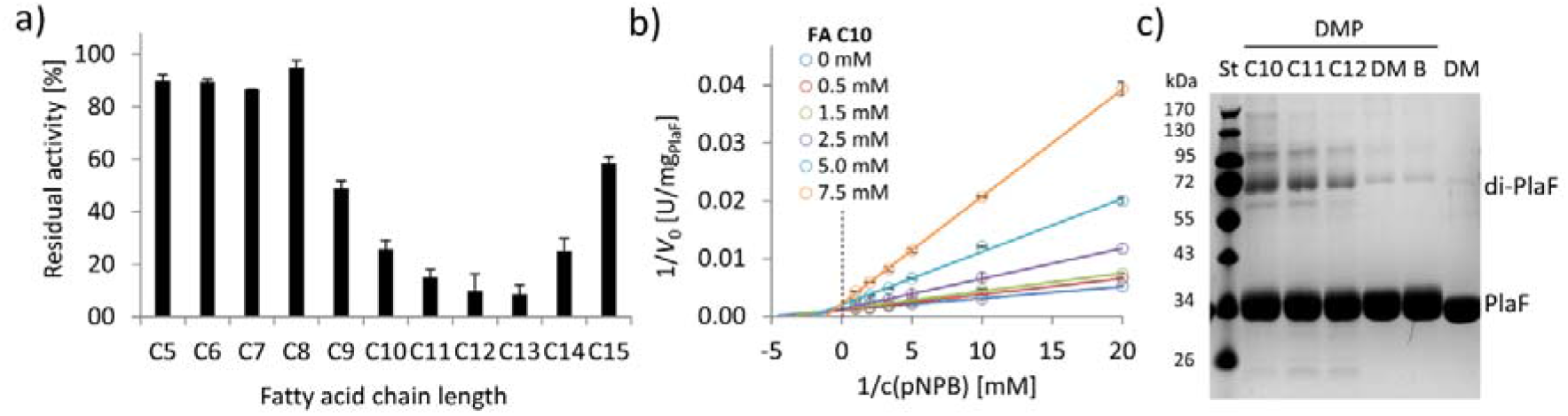
FAs exert an inhibitory effect on PlaF and trigger dimerization. **a) Inhibition of PlaF with FAs.** Esterase activity of PlaF was measured in the presence of 7.5 mM FA (C5 – C15); an untreated PlaF sample was set as 100 %. The results are mean ± S.D. of three experiments with three samples each. **b) Kinetic studies with FA C10 show evidence of mixed-inhibition.** Double-reciprocal plots of initial reaction velocities measured with the *p*-NPB substrate, and FA C10 inhibitor at concentrations in a range 0-7.5 mM. **c) The effect of FAs on PlaF dimerization.** PlaF samples incubated with FAs (C10, C11, C12), dimethyl sulfoxide (DM, DMSO used to dissolve FAs), and purification buffer (B, dilution control) were cross-linked with dimethyl pimelimidate (DMP).

Yet, elevated binding constants (*k*_m_) in the presence of higher concentrations of C10 FA indicate that PlaF undergoes allosteric changes affecting the binding of FAs (Fig. 6b, Table S9). We examined whether inhibitory FAs affect dimerization by cross-linking of PlaF in the presence of C10, C11, and C12 FAs. The results of SDS-PAGE revealed a significantly higher amount of dimeric PlaF in FA-treated than in untreated samples (Fig. 6c). These results suggest a potential regulatory role of FAs on PlaF activity *via* FA-induced dimerization, which is in agreement with previously demonstrated lower activity of the PlaF dimer compared to the monomer (Fig. 5).

### The tilt of monomeric PlaF in a lipid bilayer permits direct GPL access to the active site

To better understand the molecular mechanism of PlaF activation through monomerization, we performed a set of ten independent, unbiased 2 μs long MD simulations starting from dimeric or monomeric PlaF embedded in an explicit membrane with a GPL composition similar to the native *P. aeruginosa* membrane (Fig. 7a). The simulations revealed only minor intramolecular structural changes in monomeric and dimeric PlaF compared to the initial structure (RMSD_all atom_ < 4.0 Å) (Fig. S11a, Table S10). Spontaneous monomerization was not observed during the MD simulations (Fig. S11b), in line with the sub-nanomolar dissociation constant and the simulation timescale. However, in 8 and 6 out of 10 simulations started, respectively, from PlaF_A_ or PlaF_B_, a tilting of the monomer towards the membrane was observed (Fig 7b, left and Fig. S11c). This tilting motion results in the active site cleft of the catalytic domain being oriented perpendicularly to the membrane surface, such that GPL substrates can have direct access to the active site through the opening at the dimer interface (Fig 7a, right). In dimeric PlaF, this opening is, according to the model suggested from the X-ray structure, at > 5 Å above the membrane surface (Fig. 4a) so that the diffusion of a GPL from the membrane bilayer to the cleft entrance in this configuration is thermodynamically unfavorable. In all MD simulations started from the tilted PlaF monomer, the protein remains tilted (Figs. 7b, right, and S11c), which corroborates the notion that the tilted orientation is preferred over the respective configuration in di-PlaF.

**Fig. 7.**
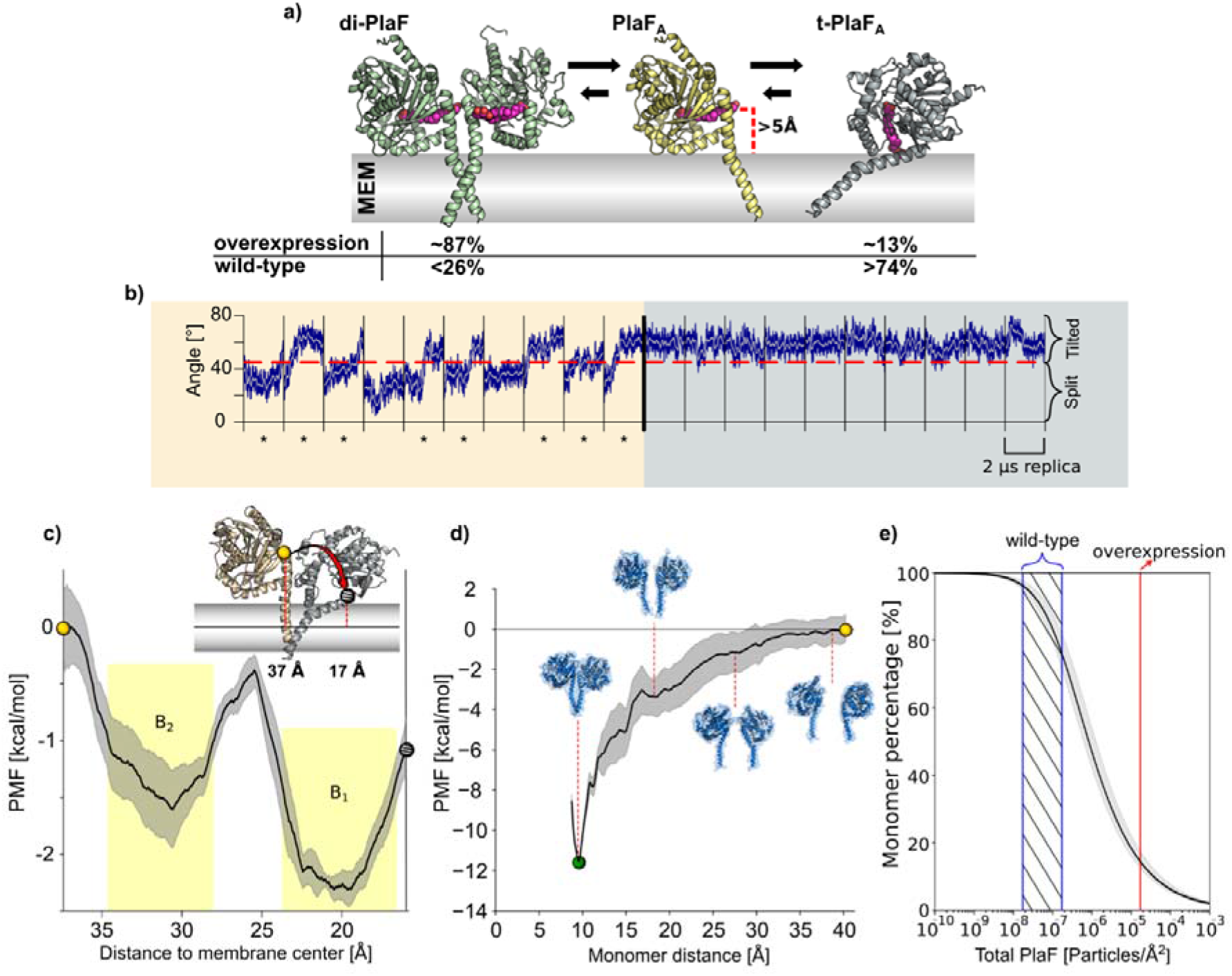
MD simulations and PMF computations of PlaF in the lipid bilayer. **a) Structures used for MD simulations.** di-PlaF: Crystal structure oriented in the membrane by the PPM method. **PlaF_A_:** Chain A from PlaF dimer oriented as in the dimer. The entrance of the active site cleft is more than 5 Å above the membrane bilayer surface. t-PlaF_A_: Extracted monomer A oriented using the PPM method. Cocrystallized MYR, 11A, and OG (not included in the simulations) are depicted in pink to highlight the orientation of the active site cleft. Arrows between the structures reflect the predicted states of equilibria under physiological conditions in *P. aeruginosa*. Percentages of the different states are obtained from the molecular simulations (see main text and panel e) **b) Molecular dynamics simulations of monomeric PlaF.** Time course of the orientation of monomeric PlaF with respect to the membrane starting from the PlaF_A_ configuration as observed in the structure (left). In 80% of the trajectories, the monomer ends in a tilted configuration (marked with *). When starting from t-PlaF_A_ (right), in all cases the structure remains tilted. This shows a significant tendency of the monomer to tilt (McNemar’s *X*^2^ = 6.125, *p* = 0.013). **c) Potential of mean force of monomer tilting.** The distance between the COM of C_α_ atoms of residues 33 to 37 (yellow, and grey spheres), and the COM of the C_18_ of the oleic acid moieties of all lipids in the membrane (continuous horizontal line in the membrane slab) was used as a reaction coordinate. The gray shaded area shows the S.D. of the mean. The yellow shaded regions are the integration limits used to calculate *K*_tilting_ (eq. S5). The spheres in the PMF relate to monomer configurations shown in the inset. **d) Potential of mean force of dimer separation.** The distance between the COM of C_α_ atoms of residues 25 to 38 of each chain was used as the reaction coordinate. The shaded area shows the standard error of the mean obtained by dividing the data into four independent parts of 50 ns each. Insets show representative structures at intermediate reaction coordinate values. **e) Percentage of PlaF monomer as a function of total PlaF concentration in the membrane according to the equilibria shown in c) and d).** The monomer percentage was computed according to eqs. S7-11 (see methods and SI for details). The red line shows the experimentally determined PlaF concentration under overexpressing conditions in *P. aeruginosa p-plaF*, while the blue-dashed region shows an estimated span for the PlaF concentration in *P. aeruginosa* wild-type (see methods for details). Calculated percentages are shown in panel a).

To further explore the transition of the monomeric PlaF_A_ to its tilted orientation (t-PlaF_A_), we calculated the free energy profile or potential of mean force (PMF) for the tilting process by using umbrella sampling and post-processing the distributions with the WHAM method [40, 41], As reaction coordinate, the distance (*d*) of the top of the JM domain (residues 33-37) to the membrane center was chosen. Distances of ^~^37 Å and ^~^17 Å were calculated for nontilted PlaF_A_ using the crystal structure and t-PlaF_A_ using the structure obtained from the unbiased MD simulations where tilting spontaneously occurred, respectively. The converged and precise (Fig. S11d; SEM < 0.4 kcal mol^−1^) PMF revealed two minima at *d* = 19.6 and 30.6 Å, with t-PlaF_A_ favored over PlaF_A_ by 0.66 kcal mol^−1^ (Fig. 7c). The free energy barrier of ^~^1.2 kcal mol^−1^ explains the rapid transition from PlaF_A_ to t-PlaF_A_ observed in the unbiased MD simulations. The equilibrium constant and free energy of PlaF tilting are *K*_tilting_ = 3.35 and a Δ*G_tilting_* = −0.8 ± 0.2 kcal mol^−1^. These results suggest a model in which PlaF is activated after monomerization by tilting with respect to the membrane surface, which allows substrate access to its catalytic site.

### Estimating the ratio of monomeric and dimeric PlaF in the cell

To investigate if dimerization-mediated PlaF inhibition is dependent on PlaF concentration in the GPL bilayer, we calculated the free energy profile of dimerization, similarly as for the tilting process. For this, the distance (*r*) between C_α_ atoms of the JM region of the two chains was used as a reaction coordinate. The converged (Fig. S11) and precise (SEM < 1.4 kcal mol^−1^) PMF revealed that di-PlaF is strongly favored at *r* = 9.5 Å (−11.4 kcal mol^−1^) over the monomer (Fig. 7d), fitting with the distance of 9.9 Å observed in the crystal structure of PlaF. From the PMF, the equilibrium constants (*K*_a_ = 1.57 × 10^7^Å^2^; *K_X_* = 2.58 × 10^5^) and free energy (Δ*G* = −7.5 ± 0.7 kcal mol^−1^) of PlaF dimerization were computed (eqs. S1-S3), taking into consideration that *K*_x_ and Δ*G* relate to a state of one PlaF dimer in a membrane of 764 lipids, according to our simulation setup. Experimentally, a concentration of one PlaF dimer per ^~^3786 lipids in *P. aeruginosa plaF*-overexpressing cells [31] was determined. However, the concentration in *P. aeruginosa* wild-type is likely 100 – 1000 fold lower, as we could not detect PlaF by Western blot (Fig. S12a). Under such physiological conditions and considering that the equilibria for dimer-to-monomer transition and titling are coupled (Fig. 7a), between 74 and 96 % of the PlaF molecules are predicted to be in a monomeric, tilted, catalytically active state in *P. aeruginosa* (Fig. 7e). Our quantitative real-time-PCR results revealed that *plaF* is constitutively expressed in *P. aeruginosa* wild-type at a much lower level than sigma factors *rpoD* and *rpoS* [42] (Fig. S12b). This is in agreement with previous global proteomic and transcriptomic results [43]. As a catalytically active form of PlaF is favored in the wild-type, PlaF is likely involved in the constant remodeling of membrane GPLs.

## Discussion

### PlaF catalyzed remodeling of membrane GPLs is a part of the bacterial Lands cycle

Employing lipidomic profiling of *P. aeruginosa* wild-type and the *plaF gene* deletion mutant, we found substantial changes in membrane GPL composition consistent with *in vitro* PLA_1_ activity of PlaF and its integral cytoplasmic membrane-localization. The present understanding of bacterial PLAs is limited to extracellular (ExoU, YplA, SlaA [26, 44]) and outer membrane (PlaB, OMPLA [45, 46]) enzymes with a proposed role in host-pathogen interactions, but, so far, bacterial PLA proteins tethered to the cytoplasmic membrane were not described [16].

Although bacterial enzymes catalyzing *de novo* GPL synthesis, their physiological functions and biochemical mechanisms are becoming increasingly well understood [16], information about GPL turnover enzymes remains largely obscure. Several of our findings indicate that PlaF plays a hitherto unexplored role in the membrane remodeling (Fig. 8) that becomes especially apparent during virulence adaptation:

i. Deletion of *plaF gene* in *P. aeruginosa* leads to accumulation of several low abundant PE, PG, and PC molecular species (Fig. 2c). PE, PG, and PE with different acyl chain lengths (C12 – C18) were hydrolyzed by PlaF *in vitro* (Fig. 2b). A low *in vitro* PLA_1_ activity of PlaF (μ*U*/mg) is expected for an enzyme that could irreversibly damage the membrane.
ii. *P. aeruginosa* Δ*plaF* strain revealed several depleted GPLs (Fig. 2c), which may be explained assuming that lysoGPLs generated by PlaF activity are further acylated to yield modified GPLs.
iii. FAs with 10 – 14 carbon atoms inhibit PlaF activity *in vitro* (Fig. 6a). As PlaF can produce such FAs *in vivo* (Fig. 2c), it is reasonable to assume that their cellular function is related to the regulation of PlaF activity by product feedback inhibition. This phenomenon is well known for lipolytic [47, 48], and other central metabolic enzymes [49–51].
iv. PlaF is constitutively expressed (Fig. S12 and reference [43]) at low levels suggesting that PlaF-catalyzed GPL remodeling may have general importance for *P. aeruginosa* physiology.
v. The *P. aeruginosa* Δ*plaF* strain shows strongly impaired killing of *G. mellonella* and human macrophages compared to wild-type (Fig. 3), thus revealing the important function of PlaF-mediated GPL remodeling for *P. aeruginosa* virulence.

**Fig. 8:**
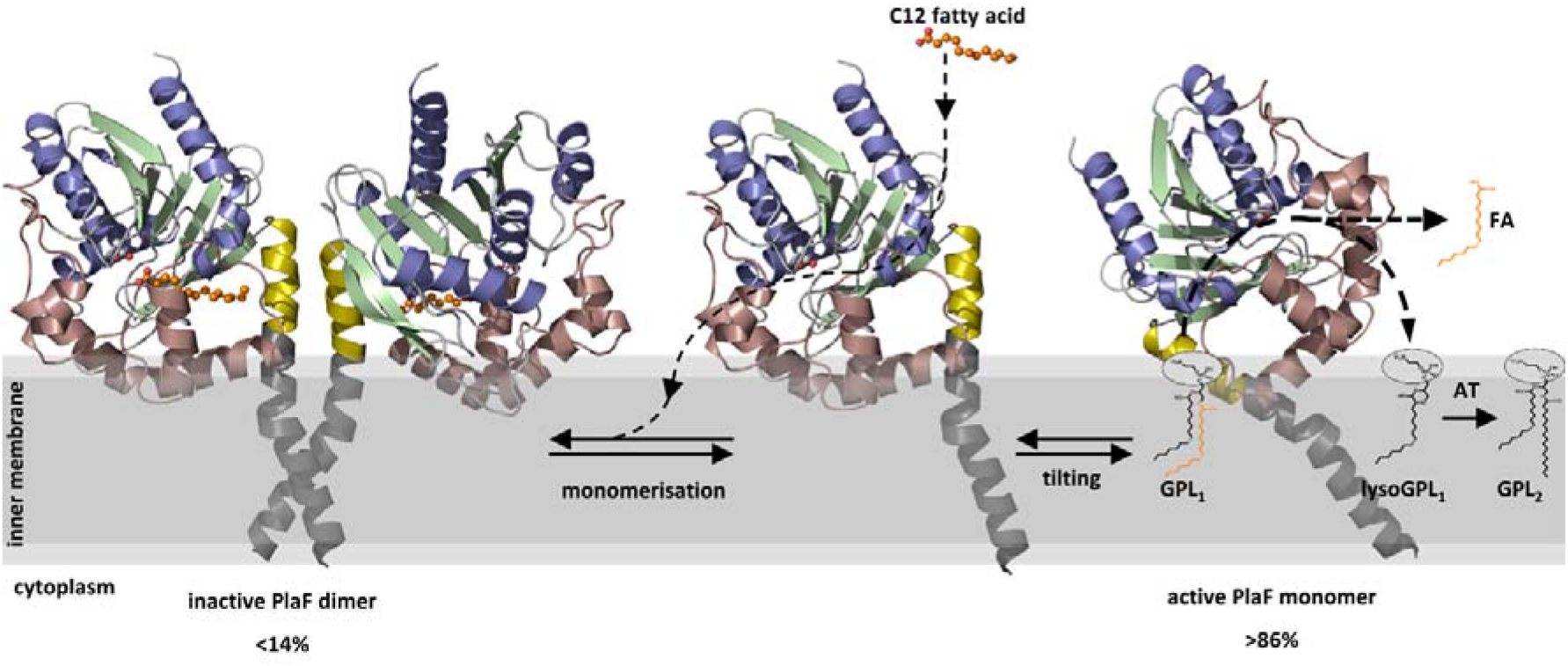
A model of PlaF-mediated membrane GPL remodeling. PlaF is anchored with the TM helix to the inner membrane of *P. aeruginosa* (Figs. 1c and 4c), where it forms an inactive dimer (Fig. 5c). Monomerization (Fig. 5c) and subsequent spontaneous tilting (Fig. 7) leads to activation. Binding of dodecanoic acid (C12) to monomeric PlaF triggers dimerization (Fig. 6c) and inhibits enzymatic activity (Fig. 6a). Tilting constrains the active site cavity of PlaF to the membrane surface such that GPL substrates can enter (GPL_1_, Fig. 2), which are hydrolyzed to FA and lysoGPL_1_. A yet unknown acyl transferase possibly acylates lyso-GPL_1_ to yield modified GPL_2_ (Fig. 2).

It is well known that the global diversity of GPL acyl chains in eukaryotes derives from *de novo* synthesis (Kennedy pathway) and remodeling (Lands cycle) pathways which are differentially regulated [52]. In the Lands cycle, GPLs are targeted by PLA and acyltransferases that remove and replace acyl chains in GPLs, respectively. We suggest that PlaF is a bacterial Lands cycle PLA which alters membranes by hydrolysis of main classes of GPLs, namely PE, PG, and PC. The exact molecular function of PlaF in GPL-remodeling and the regulation of virulence of *P. aeruginosa* remains unknown. One possibility is that PlaF tunes the concentration of low abundant GPLs species in the membrane which in turn affects the function of membrane-embedded virulence-related proteins interacting with these lipid effectors. Similar was demonstrated for ABHD6, human membrane-bound PLA, which controls the membrane concentration of lipid transmitter 2-arachidonoylglycerol involved in the regulation of the endocannabinoid receptor [53]. Human ABHD6 and PlaF share ^~^50% sequence similarity and accept similar substrates [31].

Although PlaF is an important enzyme involved in GPL metabolism, future research should reveal (i) which acyltransferase is involved in the acylation of lysoGPLs produced by PlaF, (ii) if PlaF has acyltransferase activity as described for cPLA_2_γ involved in the Lands cycle in humans [24], and (iii) if periplasmic lysophospholipase TesA [28] and the recently discovered intracellular PLA PlaB [54] are involved in the Lands cycle.

### Structural insights into dimerization and ligand-mediated regulation of PlaF activity

The high-resolution structure of PlaF with the natural ligands (fatty acids) bound to its active site represents the first dimeric structure of a full-length, single-pass TM protein (Fig. 4). It contributes to our understanding of the role of TM-JM domain-mediated dimerization for the biological activity of single-pass TM proteins which is undisputed in bacteria and eukaryotes, yet, poorly understood at the atomic level due to the lack of full-length dimeric structures [55, 56]. The present structure-function relationship of single-pass TM dimers derives from structural data of isolated TM helices in the absence of their soluble domains. Therefore, their biological relevance remains questionable [55].

The crystal structure of PlaF reveals unprecedented details of interactions between the membrane-spanning TM-JM domains and underlines the role of PlaF for degradation of membrane GPLs. The TM and JM domains are not distinct but fold into a long kinked α-helix (Fig. 4a). This is different from the structure of a human epidermal growth factor receptor (EGFR), the only structure of an isolated TM-JM domain, in which TM and JM helices are connected by an unstructured loop [57, 58]. The mechanism undergoing PlaF dimerization likely differs from the EGFR family, although it is not excluded that the truncation of soluble domains might destabilize the TM-JM dimer of EGFR leading to structural changes. We identified intramolecular interactions of at least 13 residues from the catalytic domain of PlaF with the JM domain, which clearly demonstrates the stabilizing role of the soluble domain on the TM-JM helix. Sole interactions of TM-JM helices result in the formation of a coiled-coil structure that stabilizes the PlaF dimer (Fig. 4b). The biological relevance of PlaF dimerization is corroborated by crosslinking experiments with *P. aeruginosa* cells, which revealed the *in vivo* occurrence of PlaF dimer (Fig. 5a). Furthermore, enzyme activity measurements and MST analysis of protein-protein interactions revealed that the activity decreases and dimerization increases as a function of increasing PlaF concentration *in vitro* (Fig. 5c). These findings open the question of regulation of dimerization-mediated PlaF inhibition *in vivo* and the role of membrane GPLs and their hydrolytic products in this process. Homodimerization mediated *via* TM-JM interactions was previously shown to be required for activation of single-span TM proteins from receptor tyrosine kinase [59] and ToxR-like transcriptional regulator [60]. However, structural and mechanistic details remained unknown.

A model in which PlaF-catalyzed hydrolysis of membrane GPL substrates yields FAs and lysoGPLs which are acylated to yield modified GPLs suggests that GPLs, lysoGPLs, and FAs act as potential effectors of PlaF function. A dimer interface with mainly hydrophobic interactions and a few H-bonds detected in the JM region (Fig. 4b) seems to be designed to interact with amphipathic GPLs and lysoGPLs. However, it remains to be elucidated if PlaF-GPL interactions regulate PlaF dimerization and its activity as shown for interactions of SecYEG with cardiolipin and bacteriorhodopsin with sulfated tetraglycosyldiphytanylglycerol [61, 62].

C10 – C14 FAs exert competitive inhibition as *in vitro* effectors of PlaF (Fig. 6a) and enhance dimerization (Fig. 6c). Interestingly, the range in which inhibitory activity of FA on PlaF was observed (0.5 – 7.5 mM) is the same as the intracellular concentration of FAs in bacteria [63]. The dimerization triggering function of FAs is strengthened by an observation of the mixed type inhibition (Fig. 6b), which indicates that FAs affect PlaF not only by binding to an active site but also by modulating the oligomerization equilibrium [64]. Interestingly, we identified FA ligands in the PlaF structure bound to the PlaF active site cleft (Fig. 4), although these compounds were not added exogenously prior to crystallization. Hence, they were copurified with PlaF from *P. aeruginosa* as shown by GC-MS analysis (Fig. S4). Furthermore, we identified an OG molecule, used for purification, in the active site of PlaF. These ligands form an intricate interaction network connecting the catalytic (S_137_) with the dimerization site (S_29_, T_32_, and V_33_) in the JM domain (Fig. 4e). Assuming that the OG molecule is replaced by a natural ligand in the cell, this interaction network provides a possible explanation of how FAs can mediate dimerization of the two TM-JM helices, thereby inhibiting PlaF activity.

### Atomistic model of PlaF catalyzed hydrolysis of membrane GPLs

The question remains of how does the PlaF dimer-to-monomer transition activates PlaF in the GPL bilayer? The active sites in the crystal structure of di-PlaF already adopt catalytically active conformations (Fig. 4a), suggesting that the activation of PlaF most likely does not involve structural rearrangements of the active site. To unravel a possible effect of the structural dynamics of PlaF in the membrane on enzyme regulation by dimerization, we performed extensive MD simulations, and configurational free energy computations on dimeric and monomeric PlaF embedded into a GPL bilayer mimicking the bacterial cytoplasmic membrane. While structural changes within di-PlaF, and monomeric PlaF were moderate (Table S10), monomeric PlaF spontaneously tilted as a whole towards the membrane, constraining the enzyme protein in a configuration with the opening of the active site cleft immersed into the GPL bilayer (Figs. 7a and 7b). A configuration similar to t-PlaF was observed for monomeric *Saccharomyces cerevisiae* lanosterol 14α-demethylase, a single TM spanning protein acting on a membrane-bound substrate [65]. In t-PlaF, GPL can likely access the active site cleft directly from the membrane, in contrast to di-PlaF, in which the opening of the active site cleft is > 5 Å above the membrane (Fig. 7e). There, a GPL would need to undergo the transition from the bilayer to the water milieu prior to entering the active site cleft, which is thermodynamically unfavorable.

Based on the experimental evidence, we propose a hitherto undescribed mechanism by which the transition of PlaF between a dimeric, not-tilted to a monomeric, tilted configuration is intimately linked to the modulation of the PlaF activity. This mechanism, to the best of our knowledge, expands the general understanding of mechanisms of inactivation of integral single-pass TM proteins and differs from suggested allosteric mechanisms implying structural rearrangements (even folding), mostly in the JM domain, upon ligand binding as underlying causes for functional regulation [55]. Rather, for PlaF, monomerization followed by a global reorientation of the single-pass TM protein in the membrane is the central, function-determining element.

Based on computed free energies of association (Fig. S11), and tilting (Fig. 7c), and taking into account the concentration range of PlaF in *P. aeruginosa* (Fig. 7d), PlaF preferentially exists as t-PlaF in the cytoplasmic membrane (Fig. 7d). Increasing the PlaF concentration in the membrane will thus shift the equilibrium towards the di-PlaF. This result can explain the observations that PlaF, an enzyme with membrane-disruptive activity, is found in only very low amounts (Fig. S12) in wild-type *P. aeruginosa* cells, and that overproduction of PlaF in *P. aeruginosa* is not harmful to the cells.

### Implications for the development of drugs targeting membrane remodeling

Based on our observation that *P. aeruginosa* Δ*palF* shows strongly attenuated virulence, we suggest that the bacterial Lands cycle involving PlaF might be a promising target for the development of an entirely new class of antibiotics. This class of antibiotics might be very potent if we assume that GPL remodeling plays a global role in virulence adaptation in bacteria through simultaneous regulation of several virulence-related processes.

The latest progress in the development of lipidomics methods established a clear link between alteration of membrane GPL composition and bacterial virulence and antibiotic resistance [14, 34, 66, 67]. Therefore, lipid metabolic enzymes have emerged as an intensively studied class of proteins with the potential for drug targeting [68–71]. For example, the approved antibiotic isoniazid [70] targets the fatty acid biosynthesis enzyme FabI and daptomycin most likely disturbs the assembly of GPLs in the membrane through interactions with 1-palmitoyl-2-oleoylphosphatidylglycerol. Although more compounds with similar modes of action are used [72, 73] or are in clinical trials [11] PLA-mediated GPL remodeling (Lands cycle) has not been considered as an antibiotic target.

Our results revealed that PlaF has at least two well-defined sites (active site pocket and dimerisation interface) to which drug-like compounds can bind to and modulate its activity [64, 74]. A complex active site cavity of PlaF, which is unlikely to be found in identical form in other enzymes, provides information for *in silico* design of drugs binding to PlaF with high affinity. In this process, fatty acid ligands which are bound to the active site and inhibit PlaF activity represent a good starting point. Hence, the antibacterial properties of fatty acids are well documented, although their mode of action is still poorly understood [75]. Furthermore, the activation mechanism of PlaF by monomerization indicates that compounds targeting the dimerization interface may act as agonists that dysregulate the function of the membrane by hyperactivating PlaF-mediated remodeling. PlaF seems to be suitable for the design of bivalent antibiotics which through binding to dimerisation and active sites show even higher affinity and specificity [76]. Such novel PlaF-targeting drugs might have a broad range application because PlaF homologs are conserved in several emerging bacterial pathogens (Fig. S3). Therefore, these results open up novel avenues for the development of potential drugs to extenuate *P. aeruginosa* virulence during infections through inhibition of PlaF.

## Materials and Methods

**Table 2:**
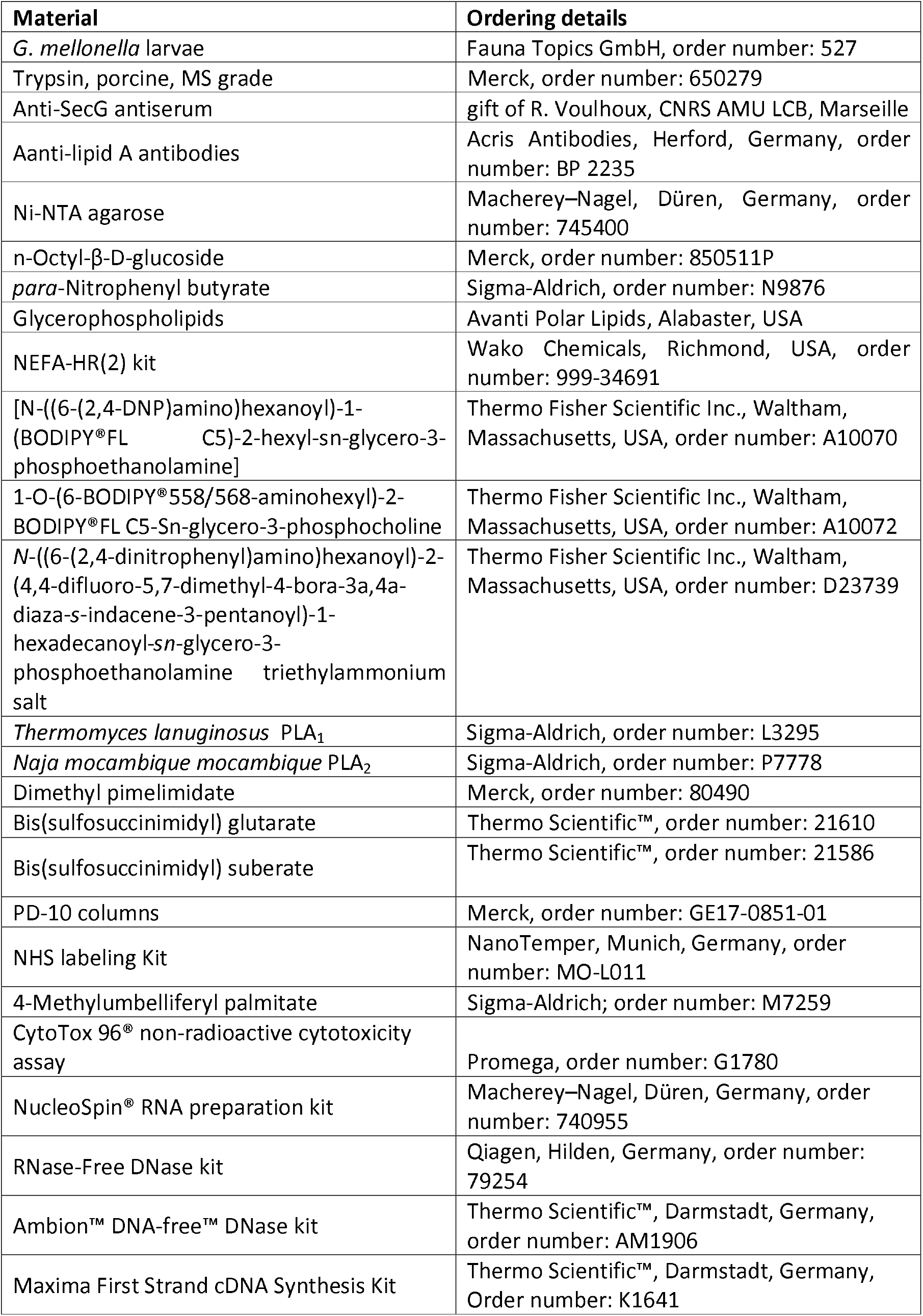
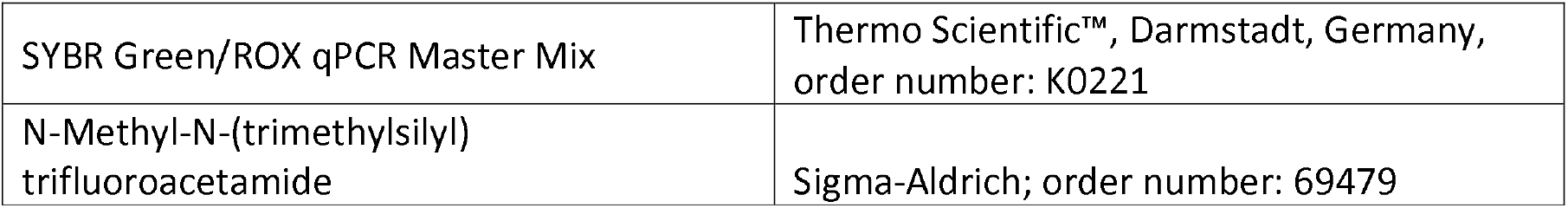
Material used in this work

### Cloning, protein expression, and purification

Molecular biology methods, DNA purification, and analysis by electrophoresis were performed as described previously [30]. For the expression of PlaF, *P. aeruginosa* PAO1 (wild-type) cells transformed [77] with plasmid pBBR-*pa2949* [30], here abbreviated as *p-plaF*, were grown overnight at 37°C in lysogeny broth (LB) medium supplemented with tetracycline (100 μg/ml) [31]. The total membrane fraction of *P. aeruginosa p-plaF* was obtained by ultracentrifugation, membranes were solubilized with Triton X-100, and PlaF was purified using Ni-NTA IMAC and buffers supplemented with 30 mM OG [31]. For biochemical analysis, PlaF was transferred to Tris-HCl (100 mM, pH 8) supplemented with 30 mM OG.

### SDS-PAGE, zymography, and immunodetection

The protein analysis by electrophoresis under denaturation conditions [78], in-gel esterase activity (zymography), and immunodetection by Western blotting were performed as described previously [30]. The protein concentration was determined by UV spectrometry using a theoretical extinction coefficient of PlaF containing a C-terminal His_6_-tag of 22,920 M ^−1^ cm^−1^ [31].

### Enzyme activity assays, inhibition, and enzyme kinetic studies

Esterase activity assays with p-nitrophenyl fatty acid esters as substrates were performed in 96-well microtiter plates as described previously [30]. Phospholipid substrates purchased from Avanti Polar Lipids (Alabaster, USA) were prepared for PLA activity assays (25 μL enzyme + 25 μL substrate) performed as described previously [79]. The amount of fatty acids released by the PLA activity of PlaF was determined using the NEFA-HR(2) kit (Wako Chemicals, Richmond, USA) [31]. PLA_1_ and PLA_2_ activities of PlaF were measured using fluorescent substrates purchased from Thermo Fisher Scientific Inc. (Waltham, Massachusetts, USA): PLA1-PE, [N-((6-(2,4-DNP)amino) hexanoyl)-1-(BODIPY® FL C5)-2-hexyl-sn-glycero-3-phosphoethanolamine]; PLA2-PC, 1-O-(6-BODIPY®558/568-aminohexyl)-2-BODIPY®FL C5-Sn-glycero-3-phosphocholine; and PLA2-PE, *N*-((6-(2,4-dinitrophenyl)amino)hexanoyl)-2-(4,4-difluoro-5,7-dimethyl-4-bora-3a,4a-diaza-s-indacene-3-pentanoyl)-1-hexadecanoyl-*sn*-glycero-3-phosphoethanolamine triethylammonium as described by da Mata Madeira and coworkers [80]. Measurements were performed using a plate reader in 96-well plates at 25°C by combining 50 μL of the substrate with 50 μL PlaF (0.7 μg/mL), or control enzymes, the PLA_1_ of *Thermomyces lanuginosus* (5 (*U*/mL) and the PLA_2_ or *Naja mocambigue mocambigue* (0.7 (*U*/mL).

#### Inhibition

The inhibition of PlaF by FAs was assayed by combining FA dissolved in DMSO (20 fold stock solution) with para-*n*itrophenyl butyrate (*p*-NPB) substrate solution followed by the addition of the PlaF sample (8 nmol) and spectrophotometric enzyme activity measurement using p-NPB substrate [81]. In control experiments, all compounds except FA were combined to assess PlaF activity in absence of FA. Inhibition constants were calculated by fitting enzyme kinetic parameters obtained by varying FA concentration (0, 0.5, 1.5, 2.5, 5 and 7.5 mM) for different substrate concentrations (0.05, 0.1, 0.2, 0.3, 0.5 and 1 mM) [82].

### Subcellular localization

Membranes from *P. aeruginosa* wild-type and *p-plaF* (PlaF overproduction strain) were isolated as described previously [30]. To separate integral from peripheral membrane proteins, total cell membranes were incubated for 30 min at room temperature with: 10 mM Na_2_CO_3_ (pH = 11), 4 M urea (in 20 mM MES buffer pH = 6.5) or 2 % (w/v) Triton X-100 (in 20 mM MES buffer pH = 6.5). After the incubation, the samples were centrifuged for 30 min at 180,000 *g* to separate membranes from solubilized proteins.

The separation of the inner and outer membrane was performed with a discontinuous sucrose gradient by ultracentrifugation at 180,000 *g* for 72 h and 4°C [83]. The sucrose gradient consisted of 1.5 ml fractions with 35, 42, 46, 50, 54, 58, 62 and 65 % (w/v) sucrose in 100 mM Tris-HCl pH 7.4. Isolated membranes from *P. aeruginosa* wild-type were suspended in buffer containing 35 % (w/v) sucrose and loaded on the top of the discontinuous sucrose gradient. Fractions were collected from the bottom (pierced tube) and sucrose concentration was determined with a refractometer (OPTEC, Optimal Technology, Baldock UK). To determine the orientation of catalytic PlaF domain *P. aeruginosa p-plaF* cells (10 ml culture with OD_580nm_ 1 grown in LB medium at 37°C) were harvested by centrifugation (4,000 *g*, 4°C, 5 min) and suspended in 1 ml Tris-HCl buffer (50 mM, pH 7.5, 10 % sucrose (w/v)) followed by shock freezing with liquid nitrogen [84]. Cells were thawed to room temperature and centrifuged (4,000 *g*, 4°C, 5 min) followed by incubation of the pellet for one hour on ice in Tris-HCl buffer (30 mM, pH 8.1, sucrose 20 % (w/v) EDTA 10 mM). Trypsin (20 μL, 1 mg/ml) was added to the suspension containing the cells with the permeabilized outer membrane and incubated at room temperature up to 5 h. The proteolytic reaction was stopped with 1 fold SDS-PAGE sample buffer and by incubation for 10 min at 99°C. Immunodetection of SecG with anti-SecG antiserum (gift of R. Voulhoux, CNRS AMU LCB, Marseille) and-lipid A antibodies (BP 2235, Acris Antibodies, Herford, Germany) was performed as described above for PlaF using the respective antisera at 1/2,000 and 1/1,000 dilutions.

### Cross-linking assays

*In vitro* cross-linking using the bifunctional cross-linking reagents dimethyl pimelimidate (DMP) was performed as previously described [85] with the following modifications. PlaF (10 μL, 15.5 μM) purified with OG was incubated with 6 μL freshly prepared DMP (150 mM in 100 mM Tris-HCl pH 8.4), BS^2^G (5 mM in 100 mM Tris-HCl pH 8.0) and BS^3^ (5 mM in 100 mM Tris-HCl pH 8.0) for 90 min. The cross-linking reaction was terminated with a 5 μLstop solution (50 mM Tris-HCl, 1M glycine, NaCl 150 mM, pH 8.3). For *in vivo* cross-linking, *P. aeruginosa p-plaF* and EV strains were grown in LB medium at 37°C to OD_580nm_ 1. Cells were harvested by centrifugation (10 min, 4,000 *g*, 4°C), suspended in 1/20 volume of Tris-HCl (pH 8.3, 100 mM, NaCl 150 mM), and treated with the same volume of freshly prepared cell-permeable cross-linking reagent DMP (0, 20, 30 and 50 mM in Tris-HCl buffer 100 mM, pH 8.4) for 2 h. The cross-linking reaction was terminated with the same volume of stop solution (50 mM Tris-HCl, 1 M glycine, 150 mM NaCl, pH 8.3).

### Analysis of concentration-dependent dimerization

Purified PlaF (20 μL, 50-60 μM) was transferred from the purification buffer into the labeling buffer (Na-PO_4_ 20 mM, pH 8.3) supplemented with OG (30 mM) using PD-10 columns (GE Healthcare, Solingen, Germany) according to the manufacturer’s protocol. Labeling was performed by incubating PlaF with 15 μL dye (440 μM stock solution) for 2.5 h using the NHS labeling Kit (NanoTemper, Munich, Germany). PlaF was then transferred into a purification buffer using PD-10 columns. Nonlabeled PlaF was diluted with the same buffer in 16 steps by combining the same volume of the protein and buffer yielding samples with concentrations from 26.9 μM to 1.6 nM. Samples containing 100 nM labeled PlaF were incubated for 16 h at room temperature in the dark and microscale thermophoresis (MST) experiments were performed using the Monolith® NT.115 device (NanoTemper, Munich, Germany) with the following setup: MST power, 60 %; excitation power 20 %; excitation type, red; 25°C. Constants were calculated according to the four-parameter logistic, nonlinear regression model using Origin Pro 2018 software.

The enzymatic activity of PlaF samples used for MST analysis was assayed by combing 15 μL of enzyme and 15 μL 4-methylumbelliferyl palmitate (4-MUP, 2 mM) dissolved in purification Tris-HCl (100 mM, pH 8) containing 10 % (v/v) propan-2-ol. Fluorescence was measured during 10 min (5 seconds steps) using a plate reader in black 96-well microtiter plates at 30°C.

### *Construction of a* P. aeruginosa Δ*plaF*, *and ΔplaF::plaF strains*

The detailed procedure for generation, and validation of Δ*plaF* deletion mutant, and Δ*plaF::plaF* complemented strains by homologous recombination is provided in SI methods and in the legend of Fig. S1 [86, 87].

### G. mellonella *virulence model*

*G. mellonella* larvae (Fauna Topics GmbH) were sorted according to size and split into groups of ten in Petri dishes. *P. aeruginosa* wild-type, the Δ*plaF*, and the Δ*plaF::plaF* strains were grown overnight and sub-cultured to mid-log phase in LB media at 37°C. The bacteria were washed twice with PBS and adjusted to OD_600_ 0.055, which equals 5 × 10^4^ bacteria/μl. This suspension was diluted in PBS to the infection dose of 500 bacteria per 10 μl, which were injected into the hindmost left proleg of the insect. Hereby, PBS injections were used as infection control and untreated larvae as viability control. If more than one larvae were dying within the control group, the experiment was repeated. The survival of larvae incubated at 30°C was monitored [88].

### Cytotoxicity assay

Bone-marrow-derived macrophages (BMDMs) were isolated from the bones of C57BL/6 mice and cultured in RPMI supplemented with 20% (v/v) conditioned L929 medium to allow for differentiation into macrophages for at least 7 days. BMDMs were seeded at a concentration of 5 × 10^5^ cells in a 24-well plate. The BMDMs cells (n = 10) were infected with 5 × 10^5^ bacteria (cultivated overnight in LB medium at 37°C), which accounts for MOI 1 [89]. PBS treated cells served as viability control. Supernatants were taken at 0, 1, 3, and 6 h post-infection. LDH levels were determined (n = 6) using the CytoTox 96® Non-Radioactive Cytotoxicity Assay according to the manufacture’s protocol. As 100% killing control, uninfected cells were lysed with 1% (v/v) Triton-X100. Statistical analysis was performed using a one-way ANOVA to determine significant changes of normally distributed values obtained from two independent experiments with 10 samples each.

### Growth curves

The growth of *P. aeruginosa* wild-type and Δ*plaF* cultures in Erlenmeyer flasks (agitation at 160 rpm) was monitored by measuring OD_580nm_ during 24 h. OD_580nm_ was converted to colony-forming units (CFU) by multiplying with the factor 8 × 10^8^ experimentally determined for *P. aeruginosa* PAO1 strain from our laboratory.

### Quantitative real-time-PCR (qRT-PCR)

RNA was isolated from *P. aeruginosa* PA01 and Δ*plaF* grown overnight (37<u>°</u>C, LB medium) with the NucleoSpin® RNA preparation kit (Macherey–Nagel, Düren) and genomic DNA was quantitatively removed using RNase-Free DNase kit (Qiagen, Hilden) and Ambion™ DNA-free™ DNase kit (Thermo Scientific™, Darmstadt) according to the manufacturer’s recommendations. One μg of RNA was transcribed into cDNA using the Maxima First Strand cDNA Synthesis Kit (Thermo Scientific™, Darmstadt). For the qRT-PCR 50 ng of cDNA was mixed with SYBR Green/ROX qPCR Master Mix (Thermo Scientific™, Darmstadt) to a total volume of 20 μl and q rt-PCR was performed as described previously. [90] Following primers were used for *rpoD* (3’-CAGCTCGACAAGGCCAAGAA-5”, CCAGCTTGATCGGCATGAAC), *rpoS* (3’-CTCCCCGGGCAACTCCAAAAG-5’, 3’-CGATCATCCGCTTCCGACCAG-5’) and *plaF* (3’-CGACCCTGTTGCTGATCCAC-5’, 3’-ACGTCGTAGCTGGCCTGTTG-5’).

### Lipidomic analysis of GPLs extracted from cell membranes

The cells of *P. aeruginosa* wildtype, Δ*plaF*, and Δ*plaF::plaF* cultures grown overnight in 15 ml LB medium (Table S3) at 37°C were harvested by centrifugation at 4,000 *g* and 4°C for 15 min and suspended in 2 ml ddH_2_O followed by boiling for 10 min to inactivate phospholipases. Cells were harvested by centrifugation (4,000 *g*, 4°C, 15 min) and total lipids were extracted from the cell pellet [91]. Briefly, after boiling the water was removed by centrifugation (4,000 *g*, 4°C, 15 min). Lipids were extracted with CHCl_3_: CH_3_OH = 1:2 (v/v) and the organic phase collected. The extraction was repeated with CHCl_3_ : CH_3_OH = 2:1 (v/v) and the organic phases were combined. One volume of CHCl_3_ and 0.75 volumes of an aqueous solution containing 1 M KCl and 0.2 M H_3_PO_4_ were added to the combined chloroform/methanol extracts. Samples were vortexed and centrifuged (2,000 *g*, 5 min). The organic phase was withdrawn and the solvent of the lipid extract was evaporated under a stream of N_2_. Total lipids were dissolved in CHCl_3_: CH_3_OH = 2:1 (v/v). GPLs were quantified by Q-TOF mass spectrometry (Q-TOF 6530; Agilent Technologies, Böblingen, Germany) as described elsewhere [91]. Statistical analysis of the GPL amount was performed using the T-test and the Shapiro-Wilk method to determine significant changes of normally distributed values obtained from four *P. aeruginosa* wildtype lipidome and four Δ*plaF* samples. Ratio of PlaF and GPLs was calculated knowing GPLs extraction yield of 40 μg GPLs per 1 ml *P*. *aeruginosa p-plaF* (OD_580nm_ 1) and PlaF purification yield of ^~^1 μg from 1 ml *P. aeruginosa p-plaF* culture with OD_580nm_ 1 [31].

### Gas chromatography-mass spectrometric (GC-MS) analysis of FA extracted from PlaF

FAs were extracted from PlaF purified from 13 g *P. aeruginosa p-plaF* cells with OG using four parts of organic solvent (CHCl_3_ : CH_3_OH = 2:1). Extraction was repeated three times, the chloroform extracts were combined, chloroform was evaporated and FAs were dissolved in 200 μL chloroform. The chloroform extract was mixed with ten volumes of acetonitrile and filtered through a 0.2 μm pore size filter. For GC-MS analysis, FA extracts and standards (C10-, C11-, C14-, C15-, C16- and C18-fatty acid; C16-, C18- and C20-primary fatty alcohol) were converted into their trimethylsilylesters and trimethylsilylethers, respectively. 900 μL of the sample or standard solution (CHCl_3_ : acetonitrile = 1:5) was mixed with 100 μL N-methyl-N-(trimethylsilyl) trifluoroacetamide and heated to 80°C for 1 h. The GC-MS system consisted of a Trace GC Ultra gas chromatograph, TriPlus autosampler, and an ITQ 900 mass spectrometer (ThermoFisher Scientific, Waltham, MA, USA). Analytes were separated on a Zebron-5-HT Inferno column (60 m × 0.25 mm i.d., 0.25 μm film thickness, Phenomenex, USA). Helium was used as carrier gas at a constant gas flow of 1.0 ml/min. The oven temperature program was as follows: 80°C; 5°C/min to 340°C, held for 5 min. The injector temperature was held at 290°C, and all injections (1 μL) were made in the split mode (1:10). The mass spectrometer was used in the electron impact (EI, 70 eV) mode and scanned over the range m/z 25 - 450 with an acquisition rate of 3 microscans. The transfer line and ion source were both kept at 280°C. Data processing was performed by the use of the software XCalibur 2.0.7 (ThermoFisher Scientific). Fatty acids from the PlaF sample were identified by comparison of their retention times and mass spectra with fatty acid standards.

### Crystallization, data collection, structure determination, and analysis

PlaF purified with OG was crystallized as described previously [31]. The X-ray diffraction data were recorded at beamline ID29 of the European Synchrotron Radiation Facility (ESRF, Grenoble, France) and processed as described [31]. The structure was determined by molecular replacement using the automated pipeline “MrBUMP” from the CCP4 package [92]. In detail, a combination of PHASER [93], REFMAC [94], BUCCANEER [95], and SHELXE [96] resulted in an interpretable electron density map to expand the placed model by molecular replacement using the model built with HsaD from *Mycobacterium tuberculosis* (PDB code: 2VF2) [97]. Phase improvement was achieved by running several cycles of automated model building (ARP/wARP, CCP4) and refinement using the PHENIX [98] package. The model was further corrected by manual rebuilding using the program COOT [99]. Detailed statistics on data collection and refinement are provided in table 1. None of the residues are in disallowed regions according to Ramachandran plots generated with MolProbity (PHENIX) [100]. The secondary structure was defined according to Kabsch and Sander [101]. Interaction surface area was determined by PISA server [102]. Coordinates and structure factors for PlaF have been deposited in the Protein Data Bank under accession code 6I8W.

#### Identification of structural homologs of PlaF

PlaF structural homologs were defined as protein structures from a non-redundant subset of PDB structures with less than 90 % sequence identity to each other (PDB90 database, 12.10.2020) with a Z-score higher than 2 according to the DALI server [103]. Sequence alignment based on structural superimposition of all 357 homologs of PlaF_B_ (all 340 homologs of PlaF_A_ were among PlaF_B_ homologs) was used to identify proteins with homology in TM-JM helix of PlaF (residues 1-38). To evaluate homology, thirty-nine 3D structures with partial conservation of TM-JM helix were superimposed with the PlaF structure using Pymol (http://www.pymol.org) (Fig. S3).

### Sequence analysis

A protein sequence of PlaF was used for a BLAST search of Pseudomonas Genome Databank (https://www.pseudomonas.com/) to identify PlaF orthologs in 4660 sequenced *P. aeruginosa* genomes. Pseudomonas Genome Databank BLAST search was extended to all pathogenic *Pseudomonas* species which were designated as those with assigned risk group 2 according to the German classification of prokaryotes into risk groups. NCBI BLAST (https://blast.ncbi.nlm.nih.gov/Blast.cgi) was used to identify PlaF homologs in other pathogenic bacteria.

### Molecular dynamics simulations of dimer and monomers

The crystal structure of the PlaF dimer was used as the starting point for building the systems for MD simulations. Five missing C-terminal residues on both chains were added by using MODELLER.[104] The dimer was oriented into the membrane using the PPM server.[105] From the so oriented dimer structure, chain B was deleted to obtain a PlaF_A_ monomer in a dimer-oriented configuration; in the same way, chain A was deleted to keep PlaF_B_. Additionally, the PlaF_A_ and PlaF_B_ monomers were oriented by themselves using the PPM server, yielding tilted configurations (t-PlaF_A_ and t-PlaF_B_). These five starting configurations, di-PlaF, PlaF_A_, PlaF_B_, t-PlaF_A_ and t-PlaF_B_, were embedded into a DOPE : DOPG = 3:1 membrane with CHARMM-GUI v1.9, [106] resembling the native inner membrane of Gram-negative bacteria,[14],[107]. In each case, 10 replicas of 2 μs were calculated. For details on the molecular dynamics setup and the thermalization process, see the SI.

### Potential of mean force calculations of dimer dissociation

For calculating a configurational free energy profile (potential of mean force, PMF) of the process of dimer dissociation, 36 intermediate states were generated by separating one chain of the dimer along the membrane plane by 1 Å steps, resulting in a minimum and maximum distance between the chain centers of mass (COM) of 40.8 and 68 Å, respectively. The generated structures represent the separation process of the PlaF dimer. To sample configurations along the chain separation in a membrane environment, each intermediate state was embedded into a membrane of approximately 157 × 157 Å^2^ by using PACKMOL-Memgen,[108] and independent MD simulations of 300 ns length each, with a total simulation time of 10.8 μs. Umbrella sampling simulations were performed by restraining the initial distance between chains in every window with a harmonic potential, using a force constant of 4 kcal mol^−1^ Å^−2^; [109] the distance between the COM of C_α_ atoms of residues 25 to 38 of each monomer was used as a reaction coordinate, being restrained in every simulation. The obtained profile was integrated along the reaction coordinate to obtain an association constant (*K*_a_; SI eq. 1), which is further used to calculate the free energy of dimerization (eq. SI 3). For details on the PMF calculation, its error estimation, and the integration for the association constant and free energy, see the SI.

### Potential of mean force calculations of monomer tilting

The initial conformations used in every window for calculating the PMF of the monomer tilting were obtained from a representative trajectory of an unbiased MD simulation where spontaneous tilting occurred. The distance between the COM of C_α_ atoms of residues 33 to 37 of the monomer with the membrane center, which correlates with the tilting of the protein in the membrane environment was used to select 22 intermediate tilting configurations. The distance to the membrane center was used as a reaction coordinate and was restrained for every configuration by a harmonic potential. Based on the energy profile obtained, an equilibrium constant that describes the change from the non-tilted and tilted states (*K*_tilting_; SI eq. 5) was calculated by integrating the energy minima. For details on the PMF calculation, its error estimation and the integration of the profile, see the SI.

### PlaF dimer versus monomer proportion under in vivo conditions

The dimer to monomer equilibrium of PlaF in the membrane results from the coupling of the following equilibria:

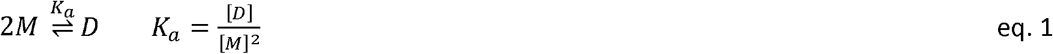

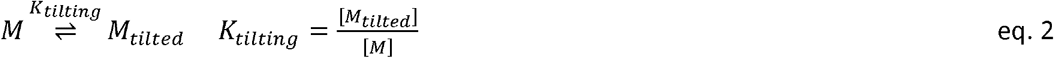

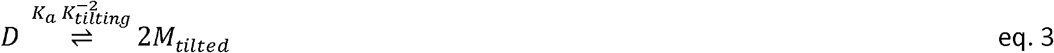

where D, M, and M_tilted_ represent the PlaF dimer, “split” monomer and tilted monomer, respectively, with *K_a_* and *K*_tilting_ being the dimer association and monomer tilting equilibrium constants, obtained from the PMF calculations. Using a total concentration of PlaF in the membrane of ^~^ 5.28 × 10^−4^ (see main text and SI for the derivation; egs. S7-S11), the fraction of PlaF in the tilted monomeric (dimeric) state was computed to be between 74 and 96% (26 and 4%) in *P. aeruginosa* (eq. S7-11). A graphical representation of the percentage of protein as a tilted monomer with respect to the protein concentration in the membrane is shown in Figure 7e. For details on the calculation of the different states, see the SI.

## Acknowledgments

This study was funded by the Deutsche Forschungsgemeinschaft (DFG, German Research Foundation), project CRC 1208 (number 267205415) to FK and KEJ (subproject A02), and HG (subproject A03). We are grateful to the beamline scientists at the European Synchrotron Radiation Facility (Grenoble, France) for assisting with the use of beamline ID29. We thank R. Voulhoux (CNRS AMU LCB, Marseille) for providing anti-SecG antiserum; P. Dollinger (HHU Düsseldorf) for help with MST measurements; M. Dick (HHU Düsseldorf) for assistance in setting up biased MD simulations. We are grateful for computational support by the “Zentrum für Informations und Medientechnologie” at the Heinrich-Heine-Universität Düsseldorf, and the computing time provided by the John von Neumann Institute for Computing (NIC) to HG on the supercomputer JUWELS at Jülich Supercomputing Centre (JSC) (user IDs: HKF7; HDD18; plaf).

## Author contributions

F.K. conceptualization, supervision, analysis, visualization, writing; K.-E.J. supervision, writing; H.G. conceptualization, supervision, analysis, writing; F.B., S.F., and M.C. investigation, analysis, visualization; S.N.S.V. investigation, analysis, visualization, writing; B.T. analysis, visualization; P.D. supervision and analysis; M.S., analysis; R.B.S. conceptualization, supervision, analysis, visualization, writing, and J.G. investigation, analysis, visualization, writing. All authors have read the final manuscript.

## Additional information

This article contains supporting information.

## References

1. Eickhoff, M.J. and B.L. Bassler, Snapshot: bacterial quorum sensing. Cell, 2018. 174(5):p. 1328–1328.

2. Parsons, J.B. and C.O. Rock, Bacterial lipids: metabolism and membrane homeostasis. Prog. Lipid Res., 2013. 52(3):p. 249–276.

3. Higgins, C.F., ABC transporters: from microorganisms to man. Annu. Rev. Cell Biol., 1992. 8(1): p. 67–113.

4. Venturi, V. and C. Fuqua, Chemical signaling between plants and plant-pathogenic bacteria. Annu. Rev. Phytopathol., 2013. 51: p. 17–37.

5. Krampen, L., et al., Revealing the mechanisms of membrane protein export by virulence-associated bacterial secretion systems. Nat. Commun., 2018. 9(1): p. 3467.

6. Mackenzie, K.R., Folding and stability of alpha-helical integral membrane proteins. Chem. Rev., 2006. 106(5):p. 1931–1977.

7. Baxter, A.A., M.D. Hulett, and I.K. Poon, The phospholipid code: a key component of dying cell recognition, tumor progression and host-microbe interactions. Cell Death. Differ., 2015. 22(12): p. 1893–1905.

8. Garcia-Fernandez, E., et al., Membrane microdomain disassembly inhibits MRSA antibiotic resistance. Cell, 2017. 171(6):p. 1354–1367.

9. Rowlett, V.W., et al., *Impact of membrane phospholipid alterations in* Escherichia coli *on cellular function and bacterial stress adaptation*. J. Bacteriol., 2017. 199(13): p. e00849–16.

10. Schniederjans, M., M. Koska, and S. Haussler, *Transcriptional and mutational profiling of an aminoglycoside-resistant* Pseudomonas aeruginosa *small-colony variant*. Antimicrob. Agents Chemother., 2017. 61(11):e01178–17.

11. Zhang, Y.M. and C.O. Rock, Membrane lipid homeostasis in bacteria. Nat. Rev. Microbiol., 2008. 6(3): p. 222–233.

12. Sinensky, M., *Homeoviscous adaptation - a homeostatic process that regulates the viscosity of membrane lipids in* Escherichia coli. Proc. Natl. Acad. Sci. U S A, 1974. 71(2): p. 522–525.

13. Cossins, A.R. Temperature adaptation of biological membranes: proceedings of the meeting held in Cambridge under the auspices of the Society for Experimental Biology in conjunction with its US/Canadian counterparts. London; Chapel Hill, NC: Portland Press, xiv, 227 (1994).

14. Benamara, H., et al., *Characterization of membrane lipidome changes in* Pseudomonas aeruginosa *during biofilm growth on glass wool*. PLoS One, 2014. 9(9): p. e108478.

15. Kanonenberg, K., et al., Shaping the lipid composition of bacterial membranes for membrane protein production. Microb. Cell Factories, 2019. 18(1): p. 131.

16. Jeucken, A., et al., *A comprehensive functional characterization of* Escherichia coli *lipid genes*. Cell Rep., 2019. 27(5):p. 1597–1606.e2.

17. Lands, W.E., Metabolism of glycerolipides; a comparison of lecithin and triglyceride synthesis. J. Biol. Chem., 1958. 231(2):p. 883–888.

18. Shindou, H. and T. Shimizu, Acyl-CoA:lysophospholipid acyltransferases. J. Biol. Chem., 2009. 284(1):p. 1–5.

19. Hishikawa, D., et al., Discovery of a lysophospholipid acyltransferase family essential for membrane asymmetry and diversity. Proc. Natl. Acad. Sci. U S A, 2008. 105(8): p. 2830–2835.

20. Clark, J.D., N. Milona, and J.L. Knopf, Purification of a 110-kilodalton cytosolic phospholipase A2 from the human monocytic cell line U937. Proc. Natl. Acad. Sci. U S A, 1990. 87(19): p. 7708–7712.

21. Song, C., et al., Molecular characterization of cytosolic phospholipase A2-beta. J. Biol. Chem., 1999. 274(24):p. 17063–17067.

22. Underwood, K.W., et al., A novel calcium-independent phospholipase A2, cPLA2-gamma, that is prenylated and contains homology to cPLA2. J. Biol. Chem., 1998. 273(34): p. 21926–21932.

23. Ohto, T., et al., Identification of novel cytosolic phospholipase A(2)s, murine cPLA(2){delta}, {epsilon}, and {zeta}, which form a gene cluster with cPLA(2}{beta}. J. Biol. Chem., 2005. 280(26):p. 24576–24583.

24. Asai, K., et al., Human group IVC phospholipase A2 (cPLA2gamma). Roles in the membrane remodeling and activation induced by oxidative stress. J. Biol. Chem., 2003. 278(10):p. 8809–8814.

25. Murakami, M., H. Sato, and Y. Taketomi, Updating Phospholipase A(2) Biology. Biomolecules, 2020. 10(10).

26. Istivan, T.S. and P.J. Coloe, Phospholipase A in Gram-negative bacteria and its role in pathogenesis. Microbiology, 2006. 152(Pt 5): p. 1263–1274.

27. Flores-Diaz, M., et al., Bacterial sphingomyelinases and phospholipases as virulence factors. Microbiol Mol Biol Rev, 2016. 80(3):p. 597–628.

28. Kovacic, F., et al., *Structural and functional characterisation of TesA - a novel lysophospholipase A from* Pseudomonas aeruginosa. PLoS One, 2013. 8(7): p. e69125.

29. Lescic Asler, I., et al., Probing enzyme promiscuity of SGNH hydrolases. Chembiochem, 2010. 11(15): p. 2158–2167.

30. Kovacic, F., et al., *A membrane-bound esterase PA2949 from* Pseudomonas aeruginosa *is expressed and purified from* Escherichia coli. FEBS Open Bio, 2016. 6(5): p. 484–493.

31. Bleffert, F., et al., Pseudomonas aeruginosa *esterase PA2949, a bacterial homolog of the human membrane esterase ABHD6: expression, purification and crystallization*. Acta Crystallogr. F Struct. Biol. Commun., 2019. 75 (Pt 4): p. 270–277.

32. Bleves, S., A. Lazdunski, and A. Filloux, *Membrane topology of three Xcp proteins involved in exoprotein transport by* Pseudomonas aeruginosa. J. Bacteriol., 1996. 178(14): p. 4297–4300.

33. Matsushita, K., et al., Isolation and characterization of outer and inner membranes from Pseudomonas aeruginosa and effect of EDTA on the membranes. J. Biochem., 1978. 83(1): p. 171–181.

34. Le Senechal, C., et al., *Phospholipid content of* Pseudomonas aeruginosa *PAO1 is modulated by the growth phase rather than the immobilization state*. Lipids, 2019. 54(9): p. 519–529.

35. MacKenzie, K.R., J.H. Prestegard, and D.M. Engelman, A transmembrane helix dimer: structure and implications. Science, 1997. 276(5309): p. 131–133.

36. Chen, J., N. Sawyer, and L. Regan, Protein-protein interactions: general trends in the relationship between binding affinity and interfacial buried surface area. Protein Sci., 2013. 22(4):p. 510–515.

37. Ollis, D.L., et al., The alpha/beta hydrolase fold. Protein Eng, 1992. 5(3): p. 197–211.

38. Chow, J., et al., The metagenome-derived enzymes LipS and LipT increase the diversity of known lipases. PLoS One, 2012. 7(10): p. e47665.

39. Jaeger, K.E., et al., Bacterial lipases. FEMS Microbiol. Rev., 1994. 15(1): p. 29–63.

40. Kumar, S., et al., The weighted histogram analysis method for free-energy calculations on biomolecules. I. The method. J. Comput. Chem., 1992. 13: p. 1011–1021.

41. Grossfield, A. WHAM: the weighted histogram analysis method. 2016. Available from: http://membrane.urmc.rochester.edu/content/wham.

42. Savli, H., et al., *Expression stability of six housekeeping genes: a proposal for resistance gene quantification studies of* Pseudomonas aeruginosa *by real-time quantitative* RT-PCR. 2003. 52(5): p. 403–408.

43. Erdmann, J., et al., Environment-driven changes of mRNA and protein levels in Pseudomonas aeruginosa. 2018. 20(11):p. 3952–3963.

44. Sawa, T., et al., Pseudomonas aeruginosa Type III secretory toxin ExoU and its predicted homologs. Toxins (Basel). 2016. 8(11): p. 307.

45. Snijder, H.J., et al., Structural evidence for dimerization-regulated activation of an integral membrane phospholipase. Nature, 1999. 401(6754): p. 717–721.

46. Schunder, E., et al., *Phospholipase PlaB is a new virulence factor of* Legionella pneumophila. Int. J. Med. Microbiol., 2010. 300(5): p. 313–323.

47. Ruiz, C., et al., *Activation and inhibition of* Candida rugosa *and* Bacillus-related *lipases by saturated fatty acids, evaluated by a new colorimetric microassay*. Biochim. Biophys. Acta, 2004. 1672(3):p. 184–191.

48. Markweg-Hanke, M., S. Lang, and F. Wagner, *Dodecanoic acid inhibition of a lipase from* Acinetobacter *sp. OPA 55*. Enzyme Microb. Technol., 1995. 17(6): p. 512–516.

49. Rose, I.A., Regulation of human red cell glycolysis: a review. Exp. Eye Res., 1971. 11(3):p. 264–272.

50. Van Schaftingen, E. and H.G. Hers, Inhibition of fructose-1,6-bisphosphatase by fructose 2,6-biphosphate. Proc. Natl. Acad. Sci. U S A, 1981. 78(5): p. 2861–2863.

51. Alam, M.T., et al., The self-inhibitory nature of metabolic networks and its alleviation through compartmentalization. Nat. Commun., 2017. 8: p. 16018.

52. Jacquemyn, J., A. Cascalho, and R.E. Goodchild, The ins and outs of endoplasmic reticulum-controlled lipid biosynthesis. 2017. 18(11): p. 1905–1921.

53. Marrs, W.R., et al., The serine hydrolase ABHD6 controls the accumulation and efficacy of 2-AG at cannabinoid receptors. Nat. Neurosci., 2010. 13(8): p. 951–7.

54. Weiler, A.J., et al., *Novel intracellular phospholipase B from* Pseudomonas aeruginosa *with activity towards endogenous phospholipids affects biofilm assembly*. BBA - Molecular and Cell Biology of Lipids, 2021. PREPRINT: https://doi.org/10.1101/2021.06.15.448513.

55. Bocharov, E.V., et al., Helix-helix interactions in membrane domains of bitopic proteins: specificity and role of lipid environment. Biochim. Biophys. Acta. Biomembr., 2017. 1859(4):p. 561–576.

56. Fink, A., et al., Transmembrane domains interactions within the membrane milieu: principles, advances and challenges. Biochim. Biophys. Acta, 2012. 1818(4): p. 974–983.

57. Bragin, P.E., et al., HER2 transmembrane domain dimerization coupled with self-association of membrane-embedded cytoplasmic juxtamembrane regions. J. Mol. Biol., 2016. 428(1):p. 52–61.

58. Endres, N.F., et al., Conformational coupling across the plasma membrane in activation of the EGF receptor. Cell, 2013. 152(3):p. 543–556.

59. Li, E. and K. Hristova, Receptor tyrosine kinase transmembrane domains: function, dimer structure and dimerization energetics. Cell. Adh. Migr., 2010. 4(2): p. 249–254.

60. Buchner, S., et al., Structural and functional analysis of the signal-transducing linker in the pH-responsive one-component system CadC of Escherichia coli. J. Mol. Biol., 2015. 427(15): p. 2548–2561.

61. Essen, L., et al., Lipid patches in membrane protein oligomers: crystal structure of the bacteriorhodopsin-lipid complex. Proc. Natl. Acad. Sci. U S A, 1998. 95(20):p. 11673–11678.

62. Corradi, V., et al., Emerging diversity in lipid–protein interactions. Chem. Rev., 2019. 119(9):p. 5775–5848.

63. Lennen, R.M., et al., Identification of transport proteins involved in free fatty acid efflux in Escherichia coli. J. Bacteriol., 2013. 195(1):p. 135–144.

64. Gabizon, R. and A. Friedler, Allosteric modulation of protein oligomerization: an emerging approach to drug design. Front. Chem., 2014. 2:p. 9–9.

65. Monk, B.C., et al., Architecture of a single membrane spanning cytochrome P450 suggests constraints that orient the catalytic domain relative to a bilayer. Proc. Natl. Acad. Sci. U S A, 2014. 111(10): p. 3865–3870.

66. El Khoury, M., et al., Targeting bacterial cardiolipin enriched microdomains: an antimicrobial strategy used by amphiphilic aminoglycoside antibiotics. Sci. Rep., 2017. 7(1): p. 10697.

67. Blanka, A., et al., *Constitutive production of c-di-GMP is associated with mutations in a variant of* Pseudomonas aeruginosa *with altered membrane composition*. Sci. Signal., 2015. 8(372): p. ra36.

68. Valentine, W.J., et al., Biosynthetic enzymes of membrane glycerophospholipid diversity as therapeutic targets for drug development. Adv. Exp. Med. Biol., 2020. 1274:p. 5–27.

69. Singh, K.S., et al., IspH inhibitors kill Gram-negative bacteria and mobilize immune clearance. Nature, 2021. 589(7843): p. 597–602.

70. Belete, T.M., Novel targets to develop new antibacterial agents and novel alternatives to antibacterial agents. Hum. Microbiome J., 2019. 11: p. 100052.

71. Brown, E.D. and G.D. Wright, Antibacterial drug discovery in the resistance era. Nature, 2016. 529(7586): p. 336–43.

72. McMurry, L.M., M. Oethinger, and S.B. Levy, Triclosan targets lipid synthesis. Nature, 1998. 394(6693): p. 531–2.

73. Zheng, C.J., et al., Cephalochromin, a FabI-directed antibacterial of microbial origin. Biochem. Biophys. Res. Commun., 2007. 362(4): p. 1107–12.

74. Hopkins, A.L. and C.R. Groom, The druggable genome. Nat. Rev. Drug. Discov., 2002. 1(9): p. 727–30.

75. Desbois, A.P. and V.J. Smith, Antibacterial free fatty acids: activities, mechanisms of action and biotechnological potential. Appl. Microbiol. Biotechnol., 2010. 85(6): p. 1629–1642.

76. Domalaon, R., et al., Antibiotic hybrids: the next generation of agents and adjuvants against Gram-negative pathogens? Clin Microbiol Rev, 2018. 31(2).

77. Choi, K.H., A. Kumar, and H.P. Schweizer, A 10-min method for preparation of highly electrocompetent Pseudomonas aeruginosa cells: application for DNA fragment transfer between chromosomes and plasmid transformation. J. Microbiol. Methods., 2006. 64(3): p. 391–397.

78. Laemmli, U.K., Cleavage of structural proteins during the assembly of the head of bacteriophage T4. Nature, 1970. 227(5259): p. 680–685.

79. Jaeger, K.E. and F. Kovacic, Determination of lipolytic enzyme activities. Methods Mol Biol, 2014. 1149: p. 111–34.

80. da Mata Madeira, P.V., et al., Structural basis of lipid targeting and destruction by the type V secretion system of Pseudomonas aeruginosa. J Mol Biol, 2016. 428 (9 Pt A): p. 1790–1803.

81. Tian, W.X. and C.L. Tsou, Determination of the rate constant of enzyme modification by measuring the substrate reaction in the presence of the modifier. Biochemistry, 1982. 21(5): p. 1028–1032.

82. Kenakin, T.P., Enzymes as Drug Targets. in Pharmacology in Drug Discovery 105–124 (2012).

83. Viarre, V., et al., HxcQ liposecretin is self-piloted to the outer membrane by Its N-terminal lipid anchor. J. Biol. Chem., 2009. 284(49): p. 33815–33823.

84. Eichler, J. and W. Wickner, The SecA subunit of Escherichia coli preprotein translocase is exposed to the periplasm. J. Bacteriol., 1998. 180(21): p. 5776–5779.

85. de Jong, L., et al., In-culture cross-linking of bacterial cells reveals large-scale dynamic protein-protein interactions at the peptide level. J. Proteome. Res., 2017. 16(7): p. 2457–2471.

86. Martinez-Garcia, E. and V. de Lorenzo, Engineering multiple genomic deletions in Gramnegative bacteria: analysis of the multi-resistant antibiotic profile of Pseudomonas putida KT2440. Environ Microbiol, 2011. 13(10): p. 2702–2716.

87. Choi, K.-H., et al., A Tn7-based broad-range bacterial cloning and expression system. Nature methods, 2005. 2(6): p. 443–448.

88. Koch, G., et al., Assessing Pseudomonas virulence with nonmammalian host: Galleria mellonella. Methods in molecular biology (Clifton, N.J.), 2014. 1149: p. 681–688.

89. Mittal, R., et al., Otopathogenic Pseudomonas aeruginosa Enters and Survives Inside Macrophages. Frontiers in microbiology, 2016. 7: p. 1828–1828.

90. Savli, H., et al., Expression stability of six housekeeping genes: A proposal for resistance gene quantification studies of Pseudomonas aeruginosa by real-time quantitative RT-PCR. J Med Microbiol, 2003. 52 (Pt 5): p. 403–408.

91. Gasulla, F., et al., The role of lipid metabolism in the acquisition of desiccation tolerance in Craterostigma plantagineum: a comparative approach. Plant J., 2013. 75(5): p. 726–741.

92. Keegan, R.M., et al., Evaluating the solution from MrBUMP and BALBES. Acta Crystallogr. D, 2011. 67: p. 313–323.

93. Mccoy, A.J., et al., Phaser crystallographic software. J. Appl. Crystallogr., 2007. 40: p. 658–674.

94. Murshudov, G.N., A.A. Vagin, and E.J. Dodson, Refinement of macromolecular structures by the maximum-likelihood method. Acta Crystallogr. D, 1997. 53: p. 240–255.

95. Cowtan, K., The Buccaneer software for automated model building. 1. Tracing protein chains. Acta Crystallogr. D, 2006. 62 (Pt 9): p. 1002–1011.

96. Hubschle, C.B., G.M. Sheldrick, and B. Dittrich, ShelXle: a Qt graphical user interface for SHELXL. J. Appl. Crystallogr., 2011. 44(Pt 6): p. 1281–1284.

97. Lack, N.A., et al., Characterization of a carbon-carbon hydrolase from Mycobacterium tuberculosis involved in cholesterol metabolism. J. Biol. Chem., 2010. 285(1):p. 434–443.

98. Adams, P.D., et al., The Phenix software for automated determination of macromolecular structures. Methods, 2011. 55(1): p. 94–106.

99. Emsley, P. and K. Cowtan, Coot: model-building tools for molecular graphics. Acta Crystallogr. D, 2004. 60(Pt 12 Pt 1): p. 2126–2132.

100. Adams, P.D., et al., PHENIX: building new software for automated crystallographic structure determination. Acta Crystallogr. D, 2002. 58(Pt 11): p. 1948–1954.

101. Kabsch, W. and C. Sander, Dictionary of protein secondary structure: pattern recognition of hydrogen-bonded and geometrical features. Biopolymers, 1983. 22(12):p. 2577–2637.

102. Krissinel, E. and K. Henrick, Inference of macromolecular assemblies from crystalline state. J. Mol. Biol., 2007. 372(3): p. 774–797.

103. Holm, L. and P. Rosenstrom, Dali server: conservation mapping in 3D. Nucleic Acids Res., 2010. 38(Web Server issue): p. W545–549.

104. Sali, A. and T.L. Blundell, Comparative protein modelling by satisfaction of spatial restraints. J. Mol. Biol., 1993. 234(3): p. 779–815.

105. Lomize, M.A., et al., OPM database and PPM web server: resources for positioning of proteins in membranes. Nucleic Acids Res, 2012. 40(Database issue): p. D370–376.

106. Jo, S., et al., CHARMM-GUI membrane builder for mixed bilayers and its application to yeast membranes. Biophys. J., 2009. 97:p. 50–58.

107. Murzyn, K., T. Rog, and M. Pasenkiewicz-Gierula, Phosphatidylethanolamine-phosphatidylglycerol bilayer as a model of the inner bacterial membrane. Biophys. J., 2005. 88(2): p. 1091–1103.

108. Schott-Verdugo, S. and H. Gohlke, PACKMOL-Memgen: a simple-to-use, generalized workflow for membrane-protein-lipid-bilayer system building. J. Chem. Inf. Model., 2019. 59(6):p. 2522–2528.

109. Torrie, G.M. and J.P. Valleau, Nonphysical sampling distributions in Monte Carlo free-energy estimation: umbrella sampling. J. Comput. Phys., 1977. 23(2): p. 187–199.

