## Supporting Information for "Evidence for a bacterial Lands cycle phospholipase A: Structural and mechanistic insights into membrane phospholipid remodeling"

This PDF file includes:

SI Materials and Methods

Figures S1 to S12

Tables S1 to S10

SI References

**SI Materials and Methods**

**Construction of a *P. aeruginosa* ∆*plaF* and *P. aeruginosa* ∆*plaF*:*plaF* strains.** The mutagenesis vector pEMG-*ΔplaF* (Fig S1a) was generated with upstream and downstream regions of *plaF* gene amplified by standard PCR using Phusion DNA polymerase, a genomic DNA of *P. aeruginosa* PAO1 as a template, and primer pairs 5`-ATATATGAATTCTCTGCTCGGCGCGAAACGCAGCGP-3`/5`-ATATATACGCGTGGGTGTCCGAAGGCTTCAGGAAAAAAGGGGC-3` and 5`-ATATATACGCGTAAACGCGAACCGGCGCCTGGG-3`/5`-CTGGATGAATTCTGGCCTGGACACCGACAAGGAAGTGATCAAGG-3`, respectively. DNA fragments upstream and downstream of the *plaF* gene were cloned into the pEMG vector by ligation of DNA fragments hydrolyzed with *Eco*RI restriction endonuclease. *P. aeruginosa* PAO1 (wild-type) cells were transformed with the pEMG-*ΔplaF* and *P. aeruginosa* *ΔplaF* mutant strain was generated by homologous recombination [1]. Generation of pUC18T-mini-Tn7T-Gm-*plaF* plasmid (Fig. S1d) for recombination of *plaF* gene containing 128-bp upstream region of *plaF* with a chromosome of *P. aeruginosa* Δ*plaF*. A DNA fragment containing the upstream region and *plaF* gene was amplified using primer pair 5’-AATAGAGCTCACCGCCGTCCTTAGGTTC-3’/5’-AATAGAGCTCCGTTTTCAGCGACCGGC-3’ from the genomic DNA of *P. aeruginosa* PAO1. Both primers contained the restriction site *Sac*I for cloning into the pUC18T-mini-Tn7T-Gm (gifts from Herbert Schweizer, Addgene plasmids #63121, #64968, and #64946). *P. aeruginosa* ∆*plaF* was transformed with pUC18T-mini-Tn7T-*plaF*-Gm and helper plasmid pTNS2 encoding the Tn7 site-specific transposase ABCD by tri-parental conjugation and the positive clones were identified by PCR using primer pair 5’-GCACATCGGCGACGTGCTCTC-3’/5’-CACAGCATAACTGGACTGATTTC-3’. The gentamycin-resistance gene was excised from *P. aeruginosa* ∆*plaF*::*plaF*-Gm by Flp-recombinase produced from pFLP3 plasmid [2].

**System construction for unbiased MD simulations**

The molecular systems for unbiased molecular dynamics simulations were built using CHARMM-GUI v1.9 [3], ensuring a distance of at least 15 Å between the protein or membrane and the solvation box boundaries. KCl at a concentration of 0.15 M was included in the solvation box to obtain a neutral system. The GPU particle mesh Ewald implementation from the AMBER16 molecular simulation suite[4, 5] with the ff14SB [6] and Lipid17 [7-9] parameters for the protein and the membrane lipids, respectively, were used; water molecules were added using the TIP3P model [10]. For each protein configuration, ten independent molecular dynamics (MD) simulations of 2 µs length were performed. Covalent bonds to hydrogens were constrained with the SHAKE algorithm[11] in all simulations, allowing the use of a time step of 2 fs. Details of the thermalization of the simulation systems are given below. All unbiased simulations showed stable protein structures (Fig. S11a) and membrane phases, evidenced by electron density and order parameter calculations (Figs. S11e and S11f). The area per lipid through all simulations calculated for the leaflet opposite to the one where PlaF was embedded was 61.3 ± 0.13 Å^2^ (mean ± SEM), similar to values reported previously [12]. All analyses were performed by using CPPTRAJ [13]. For further details on structural analyses of the MD trajectories, see below.

**Thermalization and relaxation of simulated systems**

Initially, systems were energy-minimized by three mixed steepest descent/conjugate gradient calculations with a maximum of 20,000 steps each. First, the initial positions of the protein and membrane were restrained, followed by a calculation with restraints on the protein atoms only, and finally a minimization without restraints. The temperature was maintained by using a Langevin thermostat[14], with a friction coefficient of 1 ps^‑1^. The pressure, when required, was maintained using a semi-isotropic Berendsen barostat[15], coupling the membrane (x-y) plane. The thermalization was started from the minimized structure, which was heated by gradually increasing the temperature from 10 to 100 K for 5 ps under NVT conditions, and from 100 to 300 K for 115 ps under NPT conditions at 1 bar. The equilibration process was continued for 5 ns under NPT conditions, after which production runs were started using the same conditions.

**Structural analysis of MD trajectories**

The distance between the centers of mass (COM) of residues 25 to 38 C_α_ atoms of the chains in the dimer structure was evaluated; this residue range corresponds to the solvent-accessible half of helix TM-JM (Figure 8a). The distance between the COM of C_α_ atoms of the transmembrane portion (residues 1 to 25; the first half of TM-JM helix) of each chain was also evaluated. For the monomer structures, the angle with respect to the membrane normal was assessed. For this, the angle between the membrane normal and the vector between the COM of residues 21 to 25 and residues 35 to 38 was calculated.

**Estimation of dimerization free energy from PMF**

To calculate the PlaF homodimerization PMF, values for the reaction coordinate, representing the intermonomer distance, were recorded every 2 ps and post-processed with the Weighted Histogram Analysis Method implementation of A. Grossfield (WHAM 2.0.9),[16, 17] removing the first 100 ns as an equilibration of the system. The kernel densities showed a median overlap of 8.2% between contiguous windows (Fig. S11g, upper panel), well suited for PMF calculations.[18] The error was estimated by separating the last 200 ns of data in four independent parts of 50 ns each and then calculating the standard error of the mean of the independently determined energy profiles.

The association free energy was estimated from the obtained PMF following the membrane two-body derivation from Johnston *et al.* (2012) [19] and our previous work [20]. The PMF of dimer association is integrated along the reaction coordinate to calculate an association constant (*K*_a_), which is transformed to the mole fraction scale (*K*_x_) taking into account the number of lipids *N*_L_ per surface area *A*, and this value is used to calculate the difference in free energy between dimer and monomers (*ΔG*), according to SI eqs. 1-3:

$K_{a}=\frac{\left| \left| \Omega\right| \right|}{(2{\pi)}^{2}}\int_{0}^{D} re^{\frac{-w\left( r \right)}{k_{B}T}}dr K_{x}=K_{a}\frac{N_{L}}{A} \Delta G=-RT ln(K_{x})$	(SI eqs. 1-3)

where *r* is the value of the reaction coordinate, *w*(*r*) is the PMF at value *r*, *D* is the maximum distance at which the protein is still considered a dimer, *k*_B_ is the Boltzmann constant, and *T* is the temperature at which the simulations were performed. The factor $\frac{\left| \left| \Omega\right| \right|}{(2{\pi)}^{2}}$ considers the restriction of the configurational space of the monomers upon dimer formation in terms of the sampled angle between the two chains in the dimeric state (SI eq. 4) and the accessible space for the monomers, (2π)^2^.

$\left| \left| \Omega\right| \right|=\left[ \max\left( \theta_{a} \right)-\min\left( \theta_{a} \right) \right]*\left[ \max\left( \theta_{b} \right)-\min\left( \theta_{b} \right) \right]$	(SI eq. 4)

In SI eq. 4, the angle *θ*_a_ is defined as the angle formed between the vectors connecting the COM of chain *b* with the COM of the chain *a* and with the COM of residues 25 to 38 of the latter chain; *θ*_b_ is defined analogously starting from the COM of chain *a*. A value for ||𝛺|| of 0.55 computed from SI eq. 4 indicates the fraction of the accessible space that the PlaF monomers have in the dimeric state compared to when both chains rotate independently [(2π)^2^].

**Estimation of tilting free energy**

The initial conformations used in every window for calculating the PMF of the monomer tilting were obtained from the first microsecond of MD simulations of replica 10 of PlaF_A_ (oriented as in the di-PlaF crystal structure). The distance *d* between the COM of the C_α_ atoms of residues 33 to 37 of the monomer along the z-axis and the membrane center, represented by the COM formed by the C_18_ atoms of the membrane phospholipids, was used. *d* significantly correlates (*R*^2^ = 0.997, *p* < 0.001) with the angle formed by the second half of helix αJM1 of the monomer (residues 25 to 38) and the normal vector of the membrane (Fig. S11h). Twenty-two conformations were extracted from the representative trajectory, taking the respective snapshots where *d* and the angle showed the least absolute deviation to the average value obtained by binning *d* in windows of 2 Å width and with an evenly distributed separation of 1 Å. The initial distance for every configuration was restrained by a harmonic potential with a force constant of 4 kcal mol^‑1^ Å^‑2^, and sampling was performed for 300 ns per window. The data were obtained every 2 ps and analyzed as described above, resulting in 8.6% of median overlap between kernel densities of contiguous windows (Fig. S11g, lower panel). The error was estimated in the same way as for the dimerization (see above).

For calculating the free energy difference between the obtained basins, the PMF of monomer tilting was integrated using SI eq. 5 and 6 [21]:

$K_{tilting}=\frac{\int_{B_{1}} e^{- \frac{w\left( d \right)}{k_{B}T}}dr}{\int_{B_{2}} e^{- \frac{w\left( d \right)}{k_{B}T}}dr} \Delta G_{tilting}=-RT ln K_{tilting}$	(SI eqs. 5,6)

where *d* is defined as above, *w*(*d*) is the value of the PMF at that distance, and *B*_1_ and *B*_2_ represent the basins for the tilted and split configurations, respectively. The integration limits *B*_1_ and *B*_2_ included each basin portion below half of the value between the basin minimum and the energy barrier separating the basins, respectively (Fig. 7c, yellow shaded regions).

**PlaF dimer *versus* monomer proportion under *in vivo* conditions**

Based on the association constant computed according to SI eq. 1, *K_a_* = [*D*] / [*M*]^2^ = 1.57×10^7^ Å^2^, with [*D*] and [*M*] as area concentrations of dimer and monomer, respectively, the proportion of PlaF dimer *versus* monomer in a live cell *of* *P*. *aeruginosa* can be computed. Experimentally, 40 µg GPLs per 1 ml *P. aeruginosa* p-plaF (OD_580nm_ 1) were extracted, and a PlaF purification yield of ca. 1 µg from 1 ml *P. aeruginosa* p-*plaF* culture with OD_580nm_ was obtained [22] (Tab. S3). Considering the molecular weight of PlaF of 35.5 kDa and assuming 750 Da as the average molecular weight of membrane GPL, this relates to a concentration under overexpressing conditions of ~5.28 x 10^-4^ PlaF monomers per lipid. Under non-overexpressing conditions, the concentration of PlaF monomers is estimated to be at least 100-1000 fold lower, i.e., 5.28 x 10^-6^ – 5.28 x 10^-7^ PlaF monomers per lipid. Considering that the area per lipid in a PE : PG = 3 : 1 membrane at 300 K is approximately 61 Å^2^ per leaflet (or 30.5 Å^2^ in a bilayer, computed in this work and ref. [23]), the total area concentration of PlaF molecules then is

$T = 2\left[ D \right]+ \left[ M \right]=[1.73 x {10}^{-8}, 1.73 x {10}^{-7}] \frac{\mathrm{PlaF}}{Å^{2}}$	(SI eq. 7).

Expressing the association constant in terms of the monomer concentration using SI eq. 1 yields

$K_{a}=\frac{\frac{T-\left[ M \right]}{2}}{\left[ M \right]^{2}} \Leftrightarrow2K_{a}\left[ M \right]^{2}+\left[ M \right]-T=0$	(SI eq. 8),

and solving the quadratic equation then results in

$\left[ M \right]=\frac{-1+\sqrt{1+8K_{a}T}}{4K_{a}}=\left[ 1.25\times{10}^{-8},6.00\times{10}^{-8} \right]\frac{\mathrm{PlaF}}{Å^{2}}$	(SI eq. 9)

and

$\left[ D \right]=\frac{T-\left[ M \right]}{2}=\left[ 2.43\times{10}^{-9},5.66\times{10}^{-8} \right]\frac{PlaF dimer}{Å^{2}}$	(SI eq. 10),

These results show that in live cells the fraction of PlaF in the monomeric (dimeric) state is between 35 and 72 % (65 and 28%), where the PlaF monomer is considered to be in the “split” configuration with respect to the membrane normal.

As the tilting of the PlaF monomer is energetically favorable compared to the “split” configuration and, hence, depletes the concentration of “split” PlaF monomers, the dimeric PlaF concentration will decrease (Figure 8a). To quantitatively consider the effect of the tilting, we express the overall equilibrium constant for the processes shown in Figure 8a as

$K=K_{a} K_{tilting}^{-2}=\frac{\left[ D \right]}{\left[ M_{tilted} \right]^{2}}$	(SI eq. 11),

where

$K_{tilting}=\frac{\left[ M_{tilted} \right]}{\left[ M \right]}=3.35$, equivalent to $\Delta G_{\mathrm{tilting}} = -0.72\frac{\mathrm{kcal}}{\mathrm{mol}},$computed according to SI eq. 5.

Following the same procedure as before then yields

$$\left[ M_{tilted} \right]=\left[ 1.66\times{10}^{-8},1.28\times{10}^{-7} \right]\frac{\mathrm{PlaF}}{Å^{2}}$$

$\left[ D \right]=\left[ 3.83\times{10}^{-10},2.28\times{10}^{-8} \right]\frac{PlaF dimer}{Å^{2}}$,

showing that in live cells the fraction of PlaF in the tilted monomeric (dimeric) state is between 74 and 96% (26 and 4%). A graphical representation of the percentage of protein as a tilted monomer with respect to the protein concentration in the membrane is shown in Figure 7e.

**SI Figures**

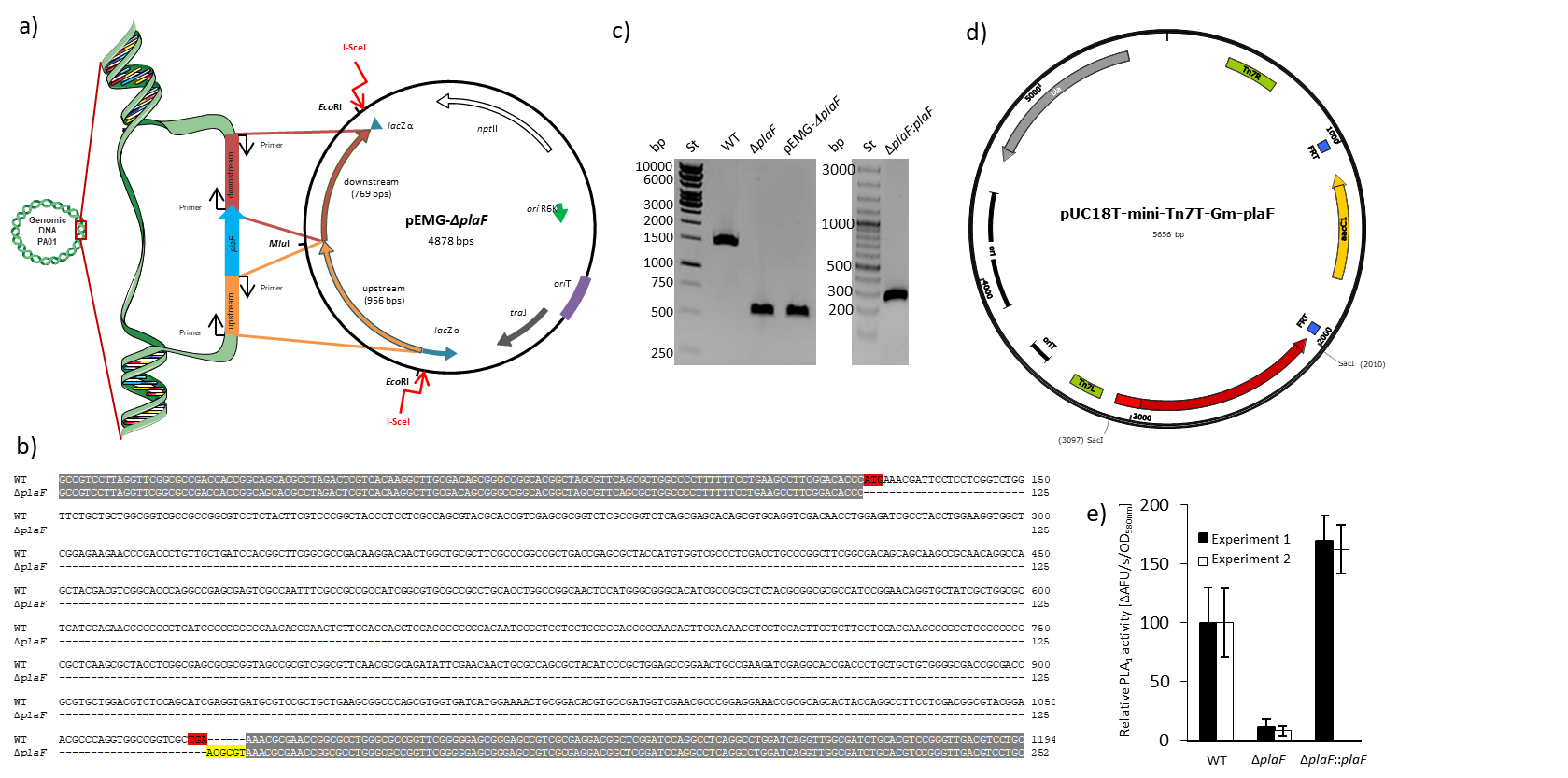

**Fig. S1: Generation of *P. aeruginosa* Δ*plaF* deletion mutant and *P. aeruginosa* Δ*plaF*::*plaF* complemented strain*.* a)** Generation of the mutagenesis vector pEMG-*ΔplaF* and homologous recombinant with a chromosome of *P. aeruginosa* PAO1 [1]. Upstream and downstream regions of *plaF* gene were amplified by standard PCR using Phusion DNA polymerase, a genomic DNA of *P. aeruginosa* PAO1 as a template, and primer pairs 5`-ATATATGAATTCTCTGCTCGGCGCGAAACGCAGCGP-3`/5`-ATATATACGCGTGGGTGTCCGAAGGCTTCAGGAAAAAAGGGGC-3` and 5`-ATATATACGCGTAAACGCGAACCGGCGCCTGGG-3`/5`-CTGGATGAATTCTGGCCTGGACACCGACAAGGAAGTGATCAAGG-3`, respectively. DNA fragments upstream and downstream of *plaF* gene were cloned into the pEMG vector by ligation of DNA fragments hydrolyzed with *Eco*RI restriction endonuclease. The white arrow indicates the neomycin phosphotransferase II gene (*npt*II) necessary for plasmid selection in the presence of kanamycin antibiotic. DNA recognition sequence for I-*Sce*I restriction endonuclease is indicated with the red arrows. **b)** Sequence alignment of DNA products of *P. aeruginosa* Δ*plaF* and the wild-type (WT) strain obtained by PCR as described in Fig S1b. The start and stop codons of *plaF* gene are indicated in red, identical nucleotides are indicated in grey, the recognition site for *Mlu*I restriction endonuclease inserted on the chromosome using the mutagenesis vector pEMG-Δ*plaF* in yellow. The company Eurofins Genomics (Ebersberg, Germany) performed DNA sequencing. **c)** Verification of *P. aeruginosa* Δ*plaF* and Δ*plaF*::*plaF* strains by PCR. Standard PCR was performed using Phusion DNA polymerase and the primers (5`-AAGGTCGCCGCCAGGAATTTCCG-3`/5`-CCGCCTGCGTGCCGACTACAAGG-3`) that bind to the up- and downstream regions of *plaF* gene. As PCR templates were used genomic DNA of *P. aeruginosa* PAO1 (WT), pEMG‑Δ*plaF* plasmid and DNA obtained from the colony of *P. aeruginosa* Δ*plaF* or Δ*plaF*::Δ*plaF* scratched from the LB agar plate and suspended in water. The expected sizes of DNA fragments were 1494 bp, 552 bp, 552 bp, and 322 bp for WT, Δ*plaF*, and pEMG‑Δ*plaF,* and Δ*plaF*::*plaF* respectively. DNA products were analyzed by electrophoresis on agarose gel (1 % w/v). The sizes of standard DNA fragments are indicated on the left. **d)** Generation of pUC18T-mini-Tn7T-Gm-plaF plasmid for recombination of *plaF* gene containing 128-bp upstream region of *plaF* with a chromosome of *P. aeruginosa* Δ*plaF*. DNA fragment containing upstream region and *plaF* gene was amplified using primer pair 5’-AATAGAGCTCACCGCCGTCCTTAGGTTC-3’/5’-AATAGAGCTCCGTTTTCAGCGACCGGC-3’ from the genomic DNA of *P. aeruginosa* PAO1. Both primers contained the restriction site *Sac*I for cloning into the pUC18T-mini-Tn7T-Gm (gifts from Herbert Schweizer, Addgene plasmids #63121, #64968 and #64946) which consists of β-lactamase resistance gene (bla), ColE1 origin of replication (ori), the origin of conjugative transfer (oriT), left and right ends of Tn7 (Tn7L and Tn7R), Flp recombinase target (FRT) and gentamycin acetyltransferase gene (aacC1).

**e)** Phospholipase A1 activity of *P. aeruginosa* Δ*plaF* and Δ*plaF*::*plaF*. *P. aeruginosa* PAO1, Δ*plaF* and Δ*plaF*::*plaF* strains were grown overnight in LB-medium at 37°C, and cells were harvested at O.D._580nm_ ~ 1 by centrifugation. Cells were suspended in Tris‑HCl buffer (100 mM, pH 8) to equal cell count, and enzyme activities of these samples were measured using PLA1 assays with N-((6-(2,4-DNP)amino)hexanoyl)-1-(BODIPY®FL C5)-2-hexyl-sn-glycero-3-phosphoethanolamine substrate. Activities are normalized on the activity of the WT which was set as 100%. The results are mean ± S.D. of two experiments with three biological replicates each measured three times.

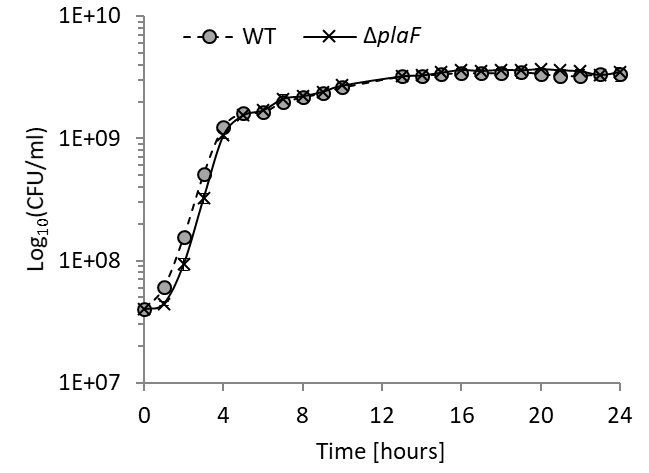

**Fig. S2:** **The growth of *P. aeruginosa* PA01 and Δ*plaF* do not differ*.*** *P. aeruginosa* strains (n = 3) were grown in LB medium in Erlenmeyer flasks at 37 °C.

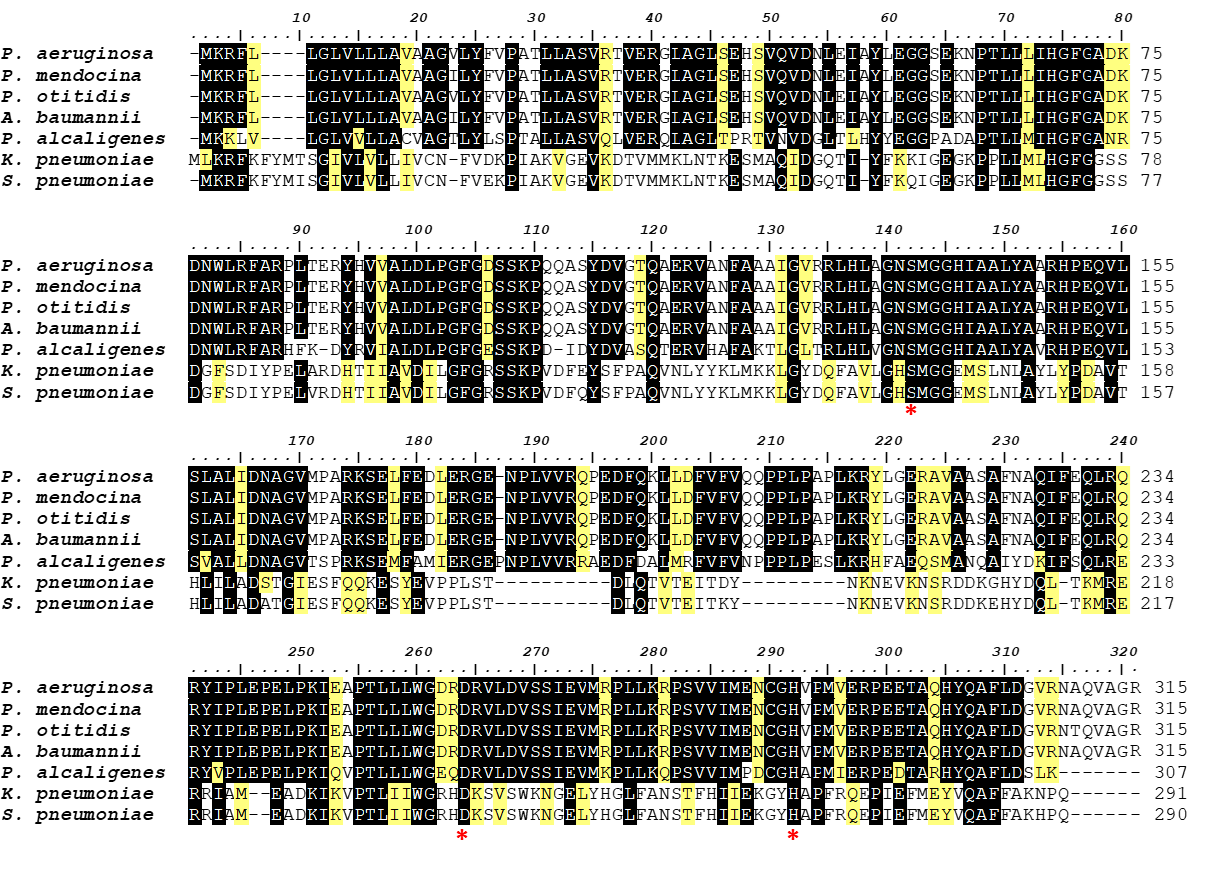

**Fig. S3: Sequence alignment of PlaF and its homologs** from *P. alcaligenes* (Pal-PlaF; sequence ID: A0A142IQQ0; sequence identity 68%, coverage 97%), *P. mendocina* (sequence ID: A0A291K7Z5; sequence identity 63%, coverage 100%), and *P. otitidis* (sequence ID: A0A1I0U5Q8; sequence identity 62%, coverage 100%), *A. baumannii* (sequence ID: [A0A1G5KTV2](https://www.uniprot.org/uniprot/A0A1G5KTV2?version=*); sequence identity 99.7%, coverage 100%), *K. pneumoniae* (sequence ID: OON65700.1; sequence identity 27%, coverage 90%) and *S. pneumonia* (sequence ID: CGG52681.1; sequence identity 28%, coverage 90%)*.* Residues identical and similar in at least five sequences were shaded in black and yellow, respectively. The catalytic triad residues of PlaF are indicated with an asterisk. The figure was prepared using BioEdit software. [3]

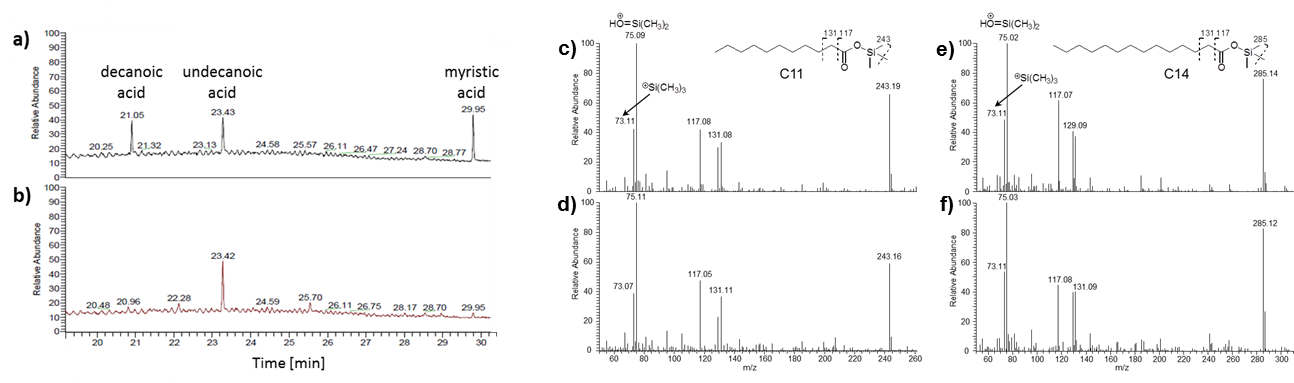

**Fig. S4: Identification of fatty acid ligands co-purified with PlaF.** Fatty acids were extracted from purified PlaF samples with organic solvent followed by silylation and gas chromatographic separation with mass spectrometric detection (GC-MS). GC chromatograms of pure C10, C11, and C14 fatty acids as standards (a) were compared to PlaF extracts (b). Mass spectrometric analysis of compounds with retention times 23.43/23.42 min and 29.95 min for fatty acid standards (c and e) and PlaF extracts (d and f) revealed the presence of undecanoic and myristic acid trimethylsilylesters, respectively. The chemical structure of undecanoic and myristic acid trimethylsilylester and characteristic fragments [4] (molecular weights in Da are indicated above) identified in mass spectra are shown.

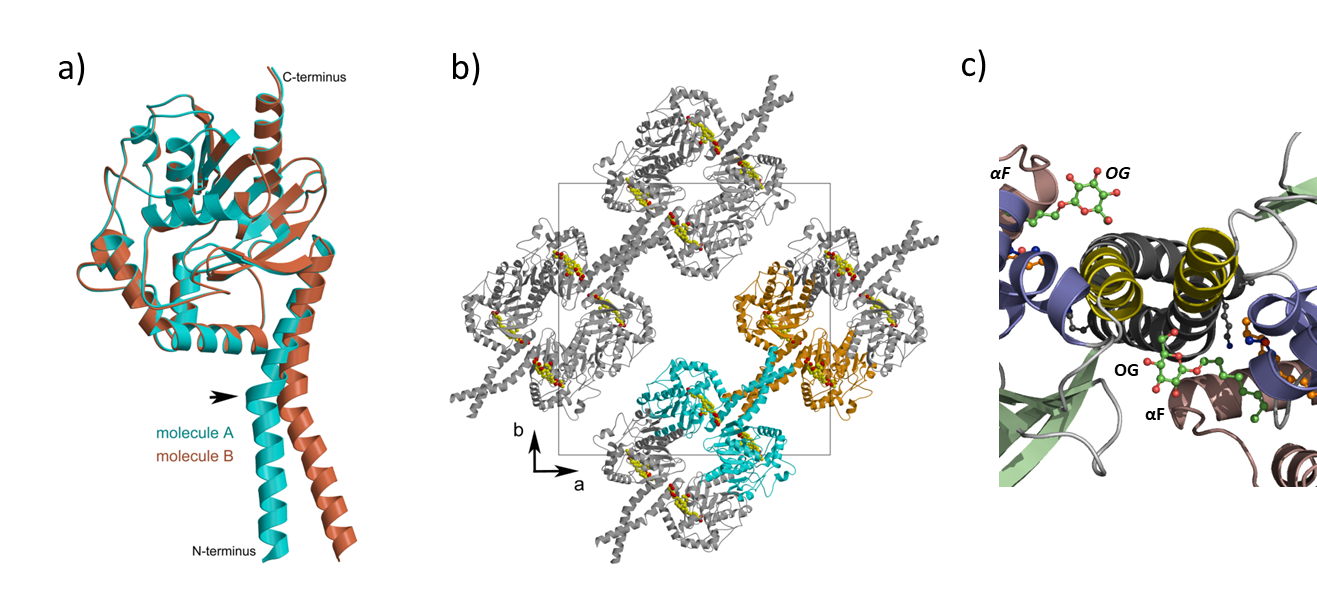

**Fig. S5: a)** Superposition of PlaF_A_ (cyan) and PlaF_B_ (orange). The arrow indicates the kink in TM-JM helix. **b)** Crystal packing of PlaF showing a four-helix bundle formed by TM-JM helices. For clarity, two dimers are shown in cyan and orange colors. The ligands present are shown as filled circles and are colored by element (carbon yellow, oxygen red). The unit cell (black square) and the a-b axes are labeled. **c)** Coiled-coil structure of the TM-JM. Enlarged section of PlaF viewed from the periplasmic side showing the coiled-coil organization of the TM-JM helix and the OG ligands located between JM and the loop preceding αF in both PlaF monomers. Elements of the PlaF_B_ molecule are indicated in italics.

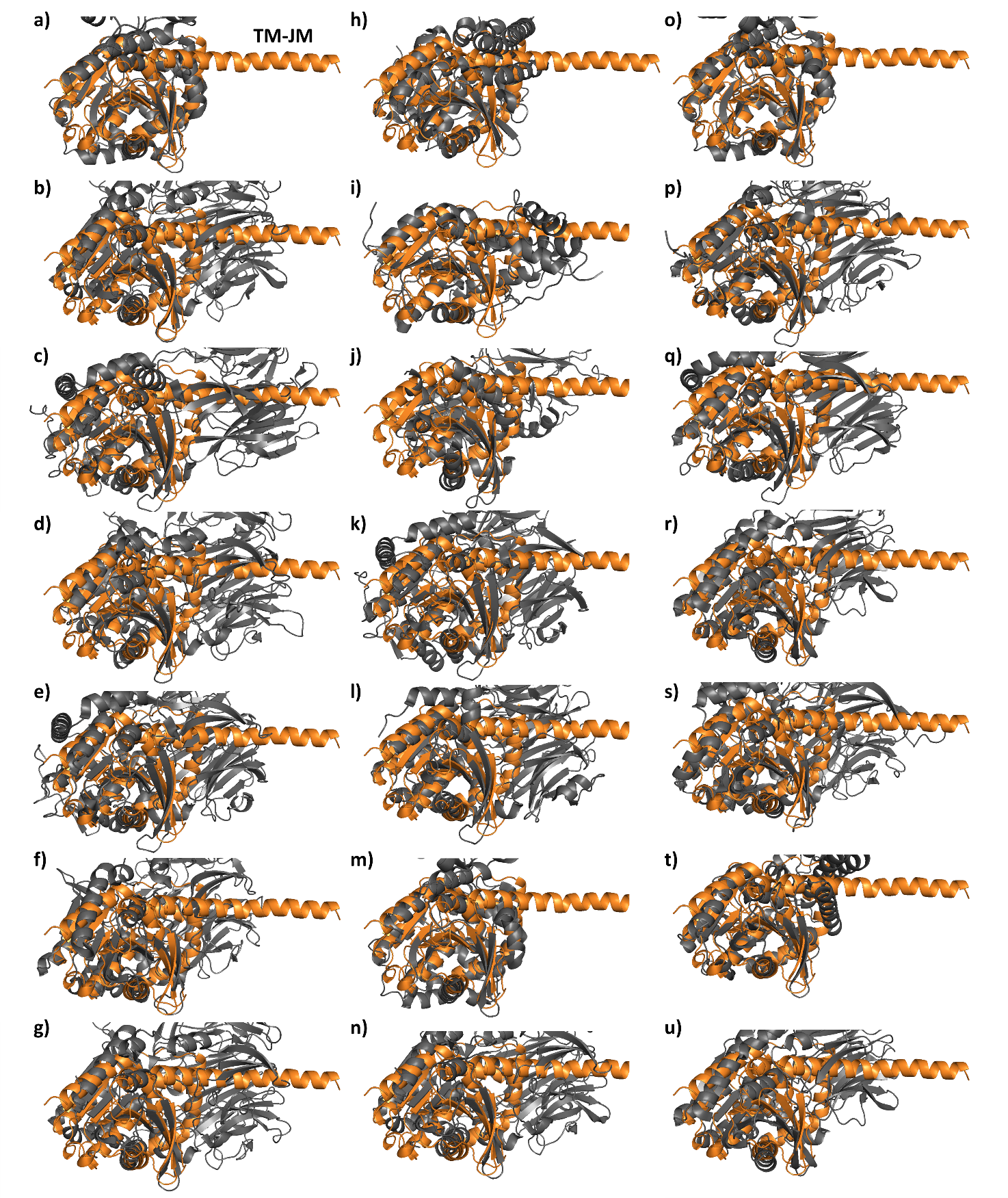

**Fig. S6: TM-JM helix of PlaF is not detected among PlaF structural homologs.** Pairwise structural alignment of PlaF_B_ (colored orange) with 1cqz (a), 1orw (b), 1yr2 (c), 1z68 (d), 2bkl (e), 2ecf (f), 2bgc (g), 2jbw (h), 2roq (i), 3mga (j), 3mun (k), 3o4j (l), 4hai (m), 4n8e (n), 5alj (o), 5l8s (p), 5t88 (q), 5yzm (r), 6eop (s), 6hxa (t), 6igp (u). PlaF structural homologs (colored gray) containing conserved sequence in TM-JM helix region of PlaF (yellow highlighted in Table S6) and in catalytic domain as identified by Dali server.

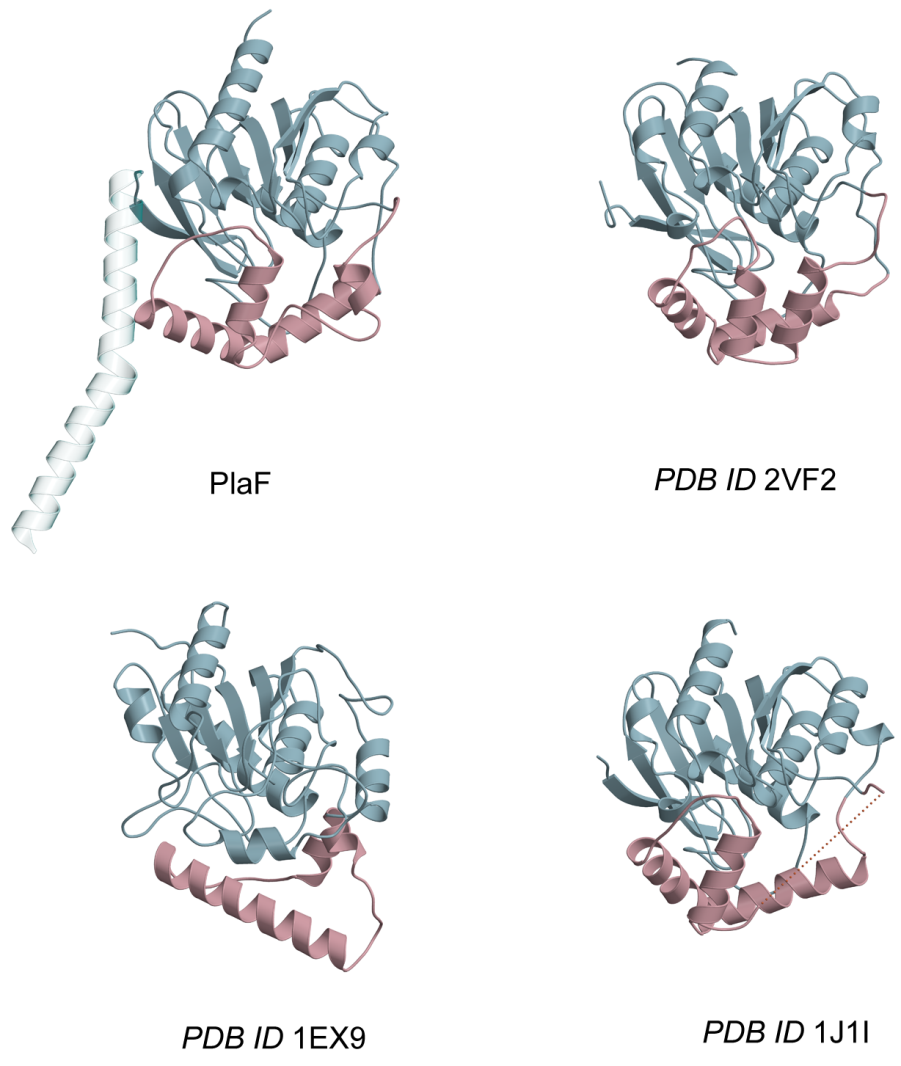

*P. aeruginosa* LipA	*Janthinobacterium* sp. CarC

PlaF	*M. tuberculosis* HsaD

**Fig. S7:** The lid-like domains shown in pink vary considerably among PlaF and hydrolytic enzymes (hydrolase HsaD from *Mycobacterium tuberculosis,* PDB ID 2VF2 [5]; lipase LipA from *P. aeruginosa*, PDB ID 1EX9 [6]; and hydrolase CarC from *Janthinobacterium* sp. strain J3, PDB ID 1J1I [7]) with structurally conserved α/β-hydrolase domains (light blue).

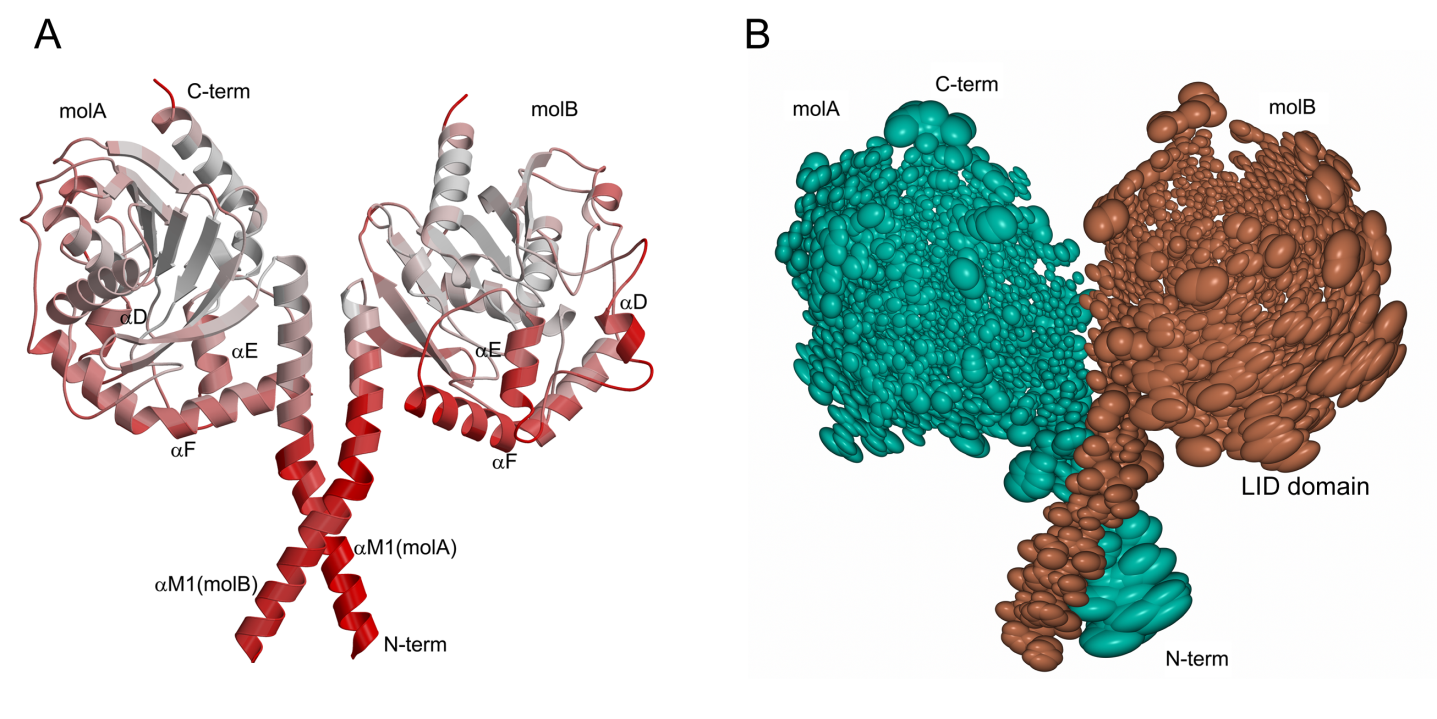

a)	b)

**Fig. S8:** **a)** Ribbon representation of the dimer structure colored according to B-factor. The lid domain and the N-terminal helices show significantly higher B-factors (color spectrum white - low B-factor, to red - high B-factor). The average B-factors of the αTM1 helix and the lid domain in PlaF dimer are ~74 and ~55 Å^2^, respectively. **b)** Thermal ellipsoid representation of the dimer scaled by the B factors combined with TLS (translation-libration-screw-rotation model) displays comparatively higher B-factors in the lid domain in molecule B and in the αTM1 helix in molecule A. The figure was prepared using CCP4mg [8].

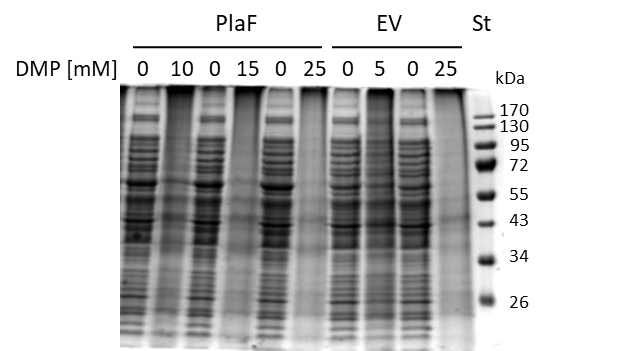

**Fig. S9: *In vivo* crosslinking and in-gel activity of PlaF.** The SDS-PAGE analysis of *in vivo* cross-linking samples of *P. aeruginosa* strain carrying pBBR1mcs-3 (empty vector control, EV) or p-*plaF* obtained in the experiment shown in Fig. 5a. Molecular weights of protein standard (St) in kDa are indicated.

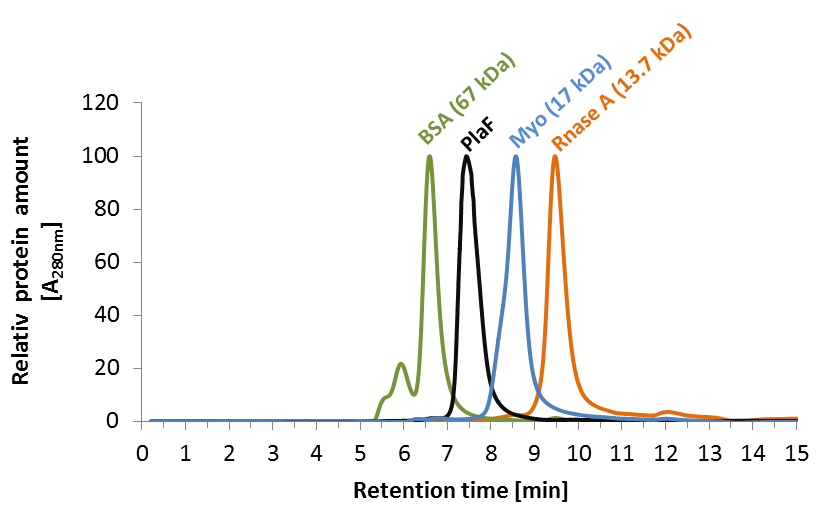

**Fig. S10: Size exclusion chromatography of PlaF showed a monomer.** PlaF (10 µl, 2 mg/ml) purified with OG and standard proteins, bovine serum albumin (BSA), equine myoglobin (Myo), and bovine ribonuclease A (RNase A) were separately analyzed using Biosep-SEC-S2000 column. Proteins were detected by measuring absorbance at a wavelength of 280 nm, and values were normalized to the maximal A_280nm_ of each protein.

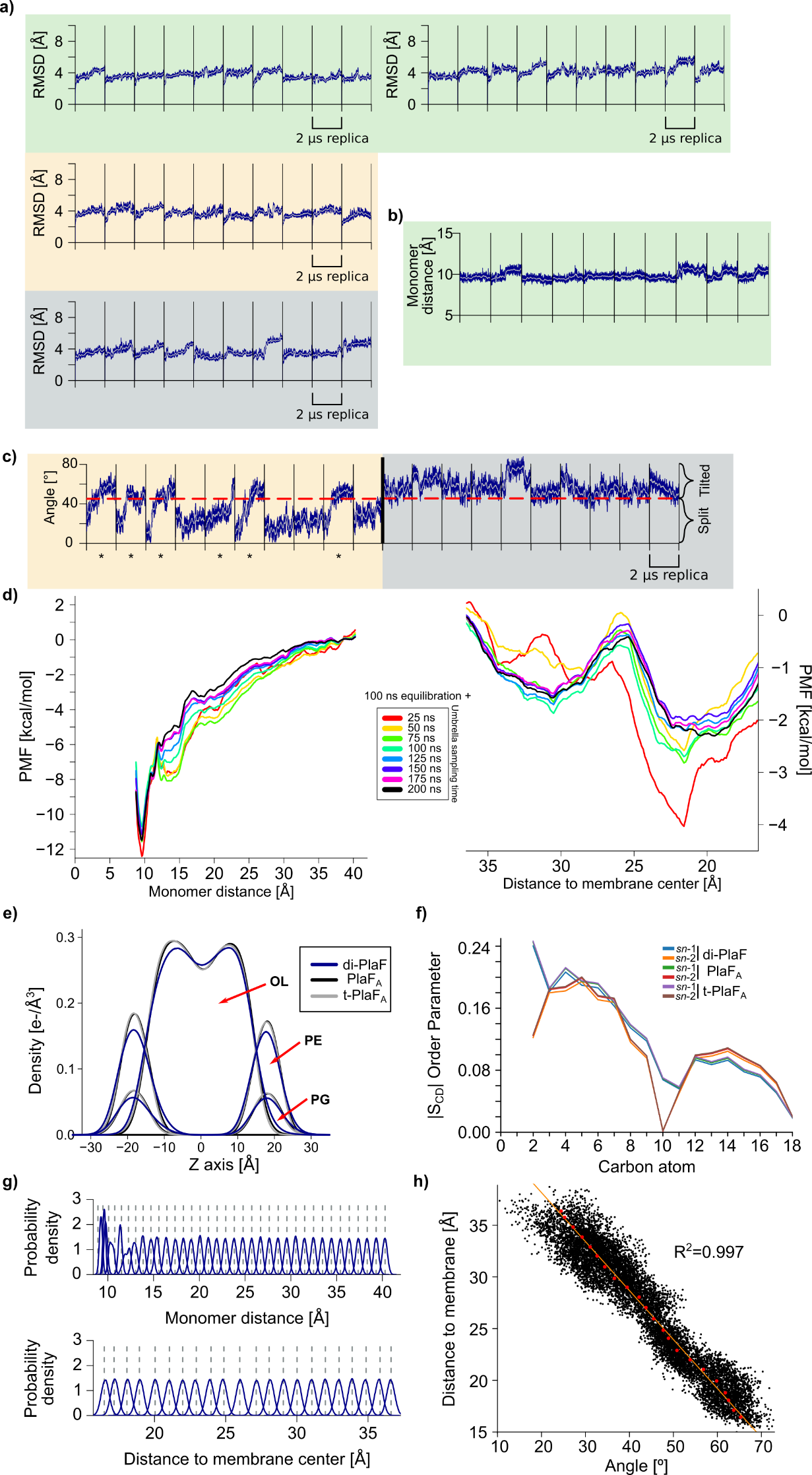

**Figure S11: Structural variations, membrane parameters, and tests for PMF convergence. a) Root mean square deviations during 10 independent, unbiased MD simulations** of I) the di-PlaF, where each chain was measured independently (green, left and right), II) monomeric PlaF started in the PlaF_A_ configuration (beige), and III) monomeric PlaF started in the tilted configuration (t-PlaF_A_, gray), computed with respect to the respective initial structure. Most simulations reached a plateau at ~ 4 Å. **b) di-PlaF monomer distance during molecular dynamics simulations.** In 10 independent replicas of 2 µs, the dimer does not show a tendency to separate on the time scale of the MD simulations according to the distance between the COM of C_α_ atoms of residues 25 to 38 of each monomer. **c) Time course of the orientation of monomeric PlaF starting from the PlaF_B_ (left) and t-PlaF_B_ (right) configurations.** Six out of ten PlaF_B_ replicas show tilting of the monomer in less than 2 μs, while all ten replicas that started in the t-PlaF_B_ configuration stayed tilted, similar to simulations started from PlaF_A_ and t-PlaF_A_**. d) Convergence of the PMFs for dimer separation (left) and monomer tilting (right).** The plots show PMFs computed every 25 ns of umbrella simulations for each window; the first 100 ns of umbrella simulations were considered equilibration phase and removed. **e) Electron density profiles of membrane components averaged over 10 independent, unbiased MD simulations of di-PlaF, PlaF_A_ or t-PlaF_A_ configurations** for the phospholipid head groups (PE and PG) and oleic acid tails (OL). The obtained shapes correspond with those generally found by experiment and MD simulations for biomembranes [7, 30, 31]. **f) S_CD_ order parameter of the lipid phase averaged over 10 independent, unbiased MD simulations of di-PlaF, PlaF_A_, or t-PlaF_A_ configurations**. The carbon atoms are numbered according to their position in the phospholipid tail. A clear dip is seen at the unsaturated position of the oleic acid tail, and the overall shape resembles a structured membrane bilayer, as previously described [7, 32]. **g) Distribution of reaction coordinate values obtained by umbrella sampling of dimer separation (top) and monomer tilting (bottom).** The dashed lines represent the restrained distance used for each window. In both cases, a force constant of 4 kcal mol^-1^ Å^-2^ was used, obtaining distributions with a median overlap of 8.2% and 8.6%, respectively. For details, see the main text. **h)** **Distance to membrane center *versus* tilting angle.** The scatter plot shows the distance of the COM of residues 33 to 37 to the membrane center *versus* the tilting angle during the first microsecond of MD simulations of the tenth replica, starting from the **PlaF_A_** configuration (*R*^2^ = 0.997, *p* < 0.001). The red dots represent the structures used as starting conformations for calculating the PMF of the tilting process.

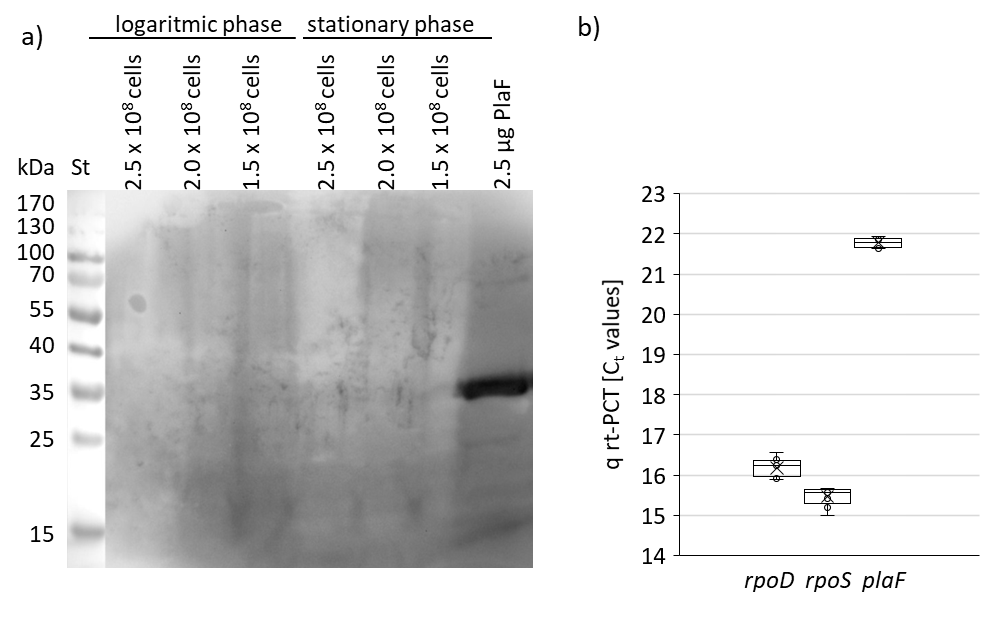

**Fig. S12:** Detection of PlaF by western blotting and q-RT-PCR. a) Western blot was performed using anti-PlaF antiserum and *P. aeruginosa* cells were from logarithmic (OD_580nm_ 1) and stationary phase (OD_580nm_ 3). Cell count was determined by counting colonies formed on LB agar plates. Purified PlaF was used as a positive control. Dimensions of *P. aeruginosa* cells in the exponential phase are 3.14 x 0.75 µm [13] which yields the surface area of 3.12 * 10^6^ nm^2^. b) Q RT-PCR analysis of transcription of *rpoD*, *rpoS*, and *plaF* genes in *P. aeruginosa* PA01 overnight culture.

**SI Tables**

**Table S1**: All phospholipid species identified in *P. aeruginosa* PA01 and ∆*plaF* by Q-TOF MS/MS.

- Excel file “**Table S1** All phospholipid species”

*Phospholipid nomenclature XX:Y. XX, the sum of carbon atoms in fatty acids bound to

**Table S2:** Phospholipid species significantly differentially abundant in *P. aeruginosa* wild-type, ∆*plaF,* and ∆*plaF*::*plaF*.

| GPL^#^ | *P. aeruginosa* PAO1 nmol/mg(GPL)/OD_580nm_ ± SD | | | *P. aeruginosa* Δ*plaF*  nmol/mg(GPL)/OD_580nm_ ± SD T-test^*^ | | | | *P. aeruginosa* Δ*plaF*::*plaF*  nmol/mg(GPL)/OD_580nm_ ± SD T-test^*^ | | | |
| --- | --- | --- | --- | --- | --- | --- | --- | --- | --- | --- | --- |
| PI 26:0 | 0.0018 | ± | 0.0007 | 0.0039 | ± | 0.0013 | 0.032 | 0.0033 | ± | 0.0009 | 0.076 |
| PE 31:2 | 0.0007 | ± | 0.0004 | 0.0019 | ± | 0.0007 | 0.040 | 0.0014 | ± | 0.0006 | 0.189 |
| PG 36:0 | 0.0006 | ± | 0.0005 | 0.0014 | ± | 0.0003 | 0.042 | 0.0007 | ± | 0.0003 | 0.699 |
| PE 22:1 | 0.0001 | ± | 0.0001 | 0.0006 | ± | 0.0002 | 0.019 | 0.0004 | ± | 0.0001 | 0.007 |
| PG 24:3 | 0.0002 | ± | 0.0001 | 0.0005 | ± | 0.0001 | 0.033 | 0.0002 | ± | 0.0001 | 0.907 |
| PC 39:0 | 0.0001 | ± | 0.0000 | 0.0002 | ± | 0.0000 | 0.040 | 0.0002 | ± | 0.0001 | 0.181 |
| PC 36:1 | 0.0005 | ± | 0.0001 | 0.0003 | ± | 0.0001 | 0.021 | 0.0004 | ± | 0.0001 | 0.175 |
| PG 23:2 | 0.0004 | ± | 0.0001 | 0.0001 | ± | 0.0001 | 0.009 | 0.0003 | ± | 0.0001 | 0.318 |
| PE 26:3 | 0.0007 | ± | 0.0000 | 0.0003 | ± | 0.0001 | 0.004 | 0.0005 | ± | 0.0002 | 0.255 |
| PC 33:1 | 0.0009 | ± | 0.0002 | 0.0004 | ± | 0.0001 | 0.004 | 0.0005 | ± | 0.0002 | 0.021 |
| PE 37:4 | 0.0024 | ± | 0.0004 | 0.0014 | ± | 0.0004 | 0.016 | 0.0013 | ± | 0.0003 | 0.015 |
| PC 32:0 | 0.0022 | ± | 0.0004 | 0.0010 | ± | 0.0004 | 0.004 | 0.0013 | ± | 0.0004 | 0.042 |
| PC 35:2 | 0.0021 | ± | 0.0003 | 0.0007 | ± | 0.0004 | 0.002 | 0.0011 | ± | 0.0005 | 0.047 |
| PC 32:1 | 0.0022 | ± | 0.0009 | 0.0007 | ± | 0.0002 | 0.033 | 0.0012 | ± | 0.0004 | 0.101 |
| PG 32:0 | 0.0033 | ± | 0.0009 | 0.0016 | ± | 0.0001 | 0.031 | 0.0032 | ± | 0.0007 | 0.806 |
| PC 35:1 | 0.0071 | ± | 0.0010 | 0.0035 | ± | 0.0018 | 0.017 | 0.0045 | ± | 0.0022 | 0.161 |
| PE 33:0 | 0.0092 | ± | 0.0009 | 0.0052 | ± | 0.0016 | 0.009 | 0.0061 | ± | 0.0018 | 0.082 |
| PC 34:1 | 0.0281 | ± | 0.0077 | 0.0117 | ± | 0.0045 | 0.015 | 0.0140 | ± | 0.0034 | 0.028 |
| PE 33:1 | 0.0546 | ± | 0.0117 | 0.0283 | ± | 0.0116 | 0.019 | 0.0324 | ± | 0.0065 | 0.026 |
| PG 35:1 | 0.0570 | ± | 0.0092 | 0.0269 | ± | 0.0144 | 0.016 | 0.0319 | ± | 0.0086 | 0.016 |
| PE 35:1 | 0.4968 | ± | 0.0869 | 0.2503 | ± | 0.1635 | 0.049 | 0.2656 | ± | 0.0804 | 0.017 |
| **Total^€^** | **0.6711** | **±** | **0.1223** | **0.3409** | **±** | **0.2021** | **-** | **0.3705** | **±** | **0.1080** | **-** |

^#^ Phospholipid nomenclature XX:Y; XX, the sum of carbon atoms in fatty acids bound to phospholipid; Y, the number of double bonds in fatty acids bound to phospholipid.

* Significance compared to the *P. aeruginosa* PAO1.

^€^ Total amount of significantly changed GPLs.

**Table S3:** Properties of the cultures used for lipid extraction.

|  | Biological replicate | *P. aeruginosa* PA01 | *P. aeruginosa* ∆*plaF* | *P. aeruginosa* ∆*plaF*::*plaF* |
| --- | --- | --- | --- | --- |
| Optical density [OD_580nm_] | 1 | 5.93 | 5.99 | - |
|  | 2 | 7.11 | 4.24 | 5.07 |
|  | 3 | 6.54 | 6.03 | 5.02 |
|  | 4 | 6.13 | 4.96 | 4.75 |
| GPL [mg]* | 1 | 54.7 | 64.1 | - |
|  | 2 | 65.5 | 63.4 | 64.6 |
|  | 3 | 65.1 | 63.8 | 64.7 |
|  | 4 | 64.9 | 63.9 | 64.5 |
| T-test** |  | - | 0.67 | 0.49 |

* Isolated from 15 ml culture.

** Significance compared to the *P. aeruginosa* PAO1.

**Table S4:** PlaF homologs in P. aeruginosa species with sequenced genomes.

🡪 Excel file “Table S4”

**Table S5:** List of interactions involving the ligand molecules.

| Ligand | Interacting residue in PlaFA | | | | Ligand | Interacting residue in PlaFB | | | |
| --- | --- | --- | --- | --- | --- | --- | --- | --- | --- |
| Atom | Distance / Å | Atom | Residue | # | Atom | Distance / Å | Atom | Residue | # |
| MYR | | | | | 11A | | | | |
| O1 | 3.28 | OD1 | ASN | 136 | C3 | 3.62 | OD1 | ASN | 77 |
| O1 | 3.47 | O | HOH | 113 | C3 | 3.34 | O | HOH | 90 |
| O1 | 3.43 | OGB | SER | 137 | C4 | 3.51 | C8' | BOG | 502 |
| C1 | 3.64 | O | PHE | 71 | C4 | 3.65 | O | HOH | 90 |
| C3 | 3.20 | OD1 | ASN | 77 | C5 | 3.57 | C5' | BOG | 502 |
| C3 | 3.89 | CB | ALA | 73 | C5 | 3.74 | C8' | BOG | 502 |
| C3 | 3.67 | CG1 | VAL | 287 | C6 | 3.04 | O | HOH | 90 |
| C4 | 3.47 | OD1 | ASN | 77 | C7 | 3.54 | O | HOH | 90 |
| C7 | 3.66 | ND2 | ASN | 77 | C10 | 3.67 | CA | ARG | 31 |
| C8 | 3.83 | C7' | BOG | 502 | C10 | 3.76 | CG | ARG | 31 |
| C10 | 3.11 | O | HOH | 166 | C10 | 3.64 | CD1 | LEU | 79 |
| O2 | 3.43 | O | PHE | 71 | O1 | 3.12 | ND2 | ASN | 77 |
| OG | | | | | O1 | 2.99 | O2 | IPA | 504 |
| O1 | 3.48 | NH2 | ARG | 80 | C | 3.45 | O2 | IPA | 504 |
| C5' | 3.73 | CD1 | LEU | 206 | O | 3.39 | O2 | IPA | 504 |
| C8' | 3.85 | CE2 | PHE | 200 | C1 | 3.56 | OD1 | ASN | 77 |
| O6 | 2.58 | OE2 | GLU | 34 | C2 | 3.62 | OD1 | ASN | 77 |
| O6 | 3.56 | CD | GLU | 34 | IPA503 | | | | |
| C3' | 3.73 | NH2 | ARG | 80 | C1 | 3.16 | OD1 | ASN | 136 |
| C3' | 3.69 | O | HOH | 28 | C1 | 3.49 | O | HOH | 22 |
| C1' | 3.63 | CG1 | VAL | 30 | C1 | 3.25 | OGB | SER | 137 |
| C7' | 3.72 | CE2 | PHE | 200 | C2 | 3.28 | O | PHE | 71 |
| C7' | 3.74 | CZ | PHE | 200 | O2 | 3.45 | C | 11A | 501 |
| C7' | 3.83 | C8 | MYR | 500 | O2 | 3.39 | O | 11A | 501 |
| C8' | 3.88 | CD | GLN | 203 | O2 | 2.99 | O1 | 11A | 501 |
| C8' | 3.56 | CG | GLN | 203 | C3 | 3.75 | CD2 | HIS | 286 |
| C8' | 3.62 | NE2 | GLN | 203 | C3 | 3.69 | NE2 | HIS | 286 |
| C8' | 3.57 | O | PRO | 204 | C3 | 3.32 | O | HOH | 171 |
| O4 | 3.42 | O | HOH | 105 | OG | | | | |
| C2 | 3.67 | O | PRO | 205 | O1 | 2.99 | NH2 | ARG | 80 |
| O2 | 2.96 | O | PRO | 205 | C5' | 3.82 | CD1 | LEU | 206 |
| O2 | 3.23 | CD | PRO | 207 | C8' | 3.87 | CE2 | PHE | 200 |
| O2 | 3.51 | C | PRO | 205 | O6 | 2.52 | OE2 | GLU | 34 |
| O2 | 3.70 | CG | PRO | 207 | O6 | 3.42 | CD | GLU | 34 |
| C6 | 3.67 | O | HOH | 49 | C3' | 3.78 | C3 | IPA | 503 |
| O6 | 3.42 | CG1 | VAL | 33 | C2' | 3.73 | CD2 | LEU | 210 |
|  |  |  |  |  | C4' | 3.62 | NH1 | ARG | 80 |
|  |  |  |  |  | O1 | 3.36 | O | HOH | 147 |
|  |  |  |  |  | C5' | 3.57 | C5 | 11A | 501 |
|  |  |  |  |  | C6' | 3.81 | CB | PRO | 204 |
|  | van-der-waals | |  |  | C1 | 3.70 | NH2 | ARG | 80 |
|  | electrostatic | |  |  | C8' | 3.51 | C4 | 11A | 501 |
|  | hydrogen | |  |  | C8' | 3.74 | C5 | 11A | 501 |
|  | ionic | |  |  | O5 | 3.14 | OE2 | GLU | 34 |
|  |  |  |  |  | C5 | 3.90 | CG1 | VAL | 30 |
|  |  |  |  |  | C6 | 3.43 | OE2 | GLU | 34 |
|  |  |  |  |  | O6 | 3.68 | CG | GLU | 34 |

**Table S6:** List of interactions involving the dimer interface.

| Source atoms | Target atoms | Distance (Å) |
| --- | --- | --- |
| Leu 5A CG  Leu 5A CD1  Leu 8A CG  Leu 8A CD2  Val 9A CA  Val 9A C  Val 9A O  Val 9A CG1  Val 9A CG2  Leu 12A C  Leu 12A CB  Leu 12A CD1  Ala 13A N  Ala 13A CA  Ala 13A C  Ala 13A O  Ala 13A CB  Val 14A N  Val 14A CA  Val 14A C  Val 14A O  Gly 17A N  Gly 17A CA  Gly 17A C  Gly 17A O  Val 18A N  Val 18A CG2  Phe 21A CE1  Phe 21A CZ  Phe 21A CE2  Phe 21A CD2  Val 22A CG2  Thr 25A CB  Thr 25A OG1  Thr 25A CG2  Ser 29A O  Ser 29A CA  Ser 29A CB  Ser 29A OG  Ser 29A CA  Ser 29A CB  Ser 29A OG  Thr 32A CB  Thr 32A CG2  Val 33A CB  Val 33A CG1  Val 33A CG2  Val 33A O  Leu 37A CG  Leu 37A CD1  Leu 37A CD2 | Val 9B CG2  Val 9B CG2  Val 9B CG1  Val 9B CG1  Val 9B CG2  Ala 13B CB  Ala 13B CB  Ala 13B CA  Ala 13B CB  Leu 12B C  Leu 12B O  Ala 13B N  Ala 13B CA  Ala 13B CB  Leu 12B CB  Ala 16B CB  Val 9B O  Leu 12B CB  Val 9B CA  Val 9B CG1  Ala 13B CB  Ala 13B CB  Ala 13B CB  Leu 10B CD2  Val 9B CG1  Ala 13B CB  Ala 13B O  Gly 17B CA  Ala 13B C  Ala 13B O  Phe 21B CE2  Phe 21B CD2  Phe 21B CE2  Phe 21B CZ  Gly 17B CA  Ala 16B C  Gly 17B N  Gly 17B CA  Ala 13B C  Ala 13B O  Ala 16B CB  Phe 21B CE2  Phe 21B CE2  Phe 21B CZ  Phe 21B CE2  Phe 21B CZ  Phe 21B CE2  Phe 21B CZ  Phe 21B CD2  Phe 21B CE2  Phe 21B CG  Phe 21B CD2  Phe 21B CD1  Phe 21B CE2  Phe 21B CZ  Phe 21B CE1  Phe 21B CG  Phe 21B CD2  Phe 21B CD1  Phe 21B CE2  Phe 21B CZ  Phe 21B CE1  Phe 21B CD1  Phe 21B CE1  Phe 21B CE2  Phe 21B CZ  Phe 21B CE1  Phe 21B CZ  Phe 21B CE1  Val 22B CG2  Val 22B CA  Val 22B CB  Thr 25B OG1  Val 22B CG2  Val 22B CA  Val 22B CB  Phe 21B C  Phe 21B O  Val 22B N  Thr 25B CB  Thr 25B OG1  Thr 25B CG2  Val 22B CG2  Phe 21B C  Phe 21B O  Val 22B N  Thr 25B CB  Thr 25B OG1  Thr 25B CG2  Val 22B CG2  Phe 21B CB  Thr 25B CG2  Phe 21B CD1  Phe 21B CE1  Thr 25B CB  Thr 25B OG1  Thr 25B CB  Thr 25B OG1  Thr 25B CG2  Thr 25B CB  Thr 25B OG1  Val 33B CG2  Val 33B CG2  Val 33B CG2  Thr 32B CG2  Ser 29B CB  Thr 32B CB  Ser 29B CA  Ser 29B CB  Ser 29B OG  Ser 29B C  Ser 29B O  Thr 32B CB  Val 33B CG2  Val 33B CG2  Thr 32B CG2  Ser 29B CB  Thr 32B CB  Thr 32B CG2  Thr 32B CB  Val 33B CG2  Val 33B CG1  Val 33B CG2  Gly 36B CA  Gly 36B O  Leu 37B N  Gly 36B CA  Gly 36B C  Val 33B C  Val 33B O  Leu 37B N  Gly 36B CA  Gly 36B C  Thr 32B C  Val 33B N  Val 33B CA  Val 33B CB  Val 33B CG1  Val 33B CG2  Thr 32B O  Leu 37B CD2  Leu 37B CD2  Gly 36B O  Arg 83B NH2  Leu 37B CD2  Leu 37B O  Leu 37B CA  Leu 37B C | 3.95  4.49  3.87  3.89  4.43  4.42  4.45  3.97  3.66  4.09  4.10  3.94  3.80  4.38  4.41  4.17  3.88  4.33  4.45  4.13  4.41  3.76  4.48  3.62  3.84  4.17  4.44 *  4.43  4.46  3.72  4.14  3.84  3.42  4.45  4.24  4.38  3.86  4.02  4.38  3.51  4.17  4.27  3.66  4.06  4.13  4.29  3.78  3.63  4.27  4.07  3.77  3.54  3.97  3.52  3.75  3.96  4.43  4.45  4.13  4.19  3.89  3.86  4.33  4.25  4.29  3.78  4.00  4.15  4.47  4.38  4.23  4.42  3.95  3.69  3.68  4.28  4.02  3.93  3.93  4.15  3.03  4.24  3.75  4.23  4.28  4.36  4.44  3.69  4.08  4.48  3.90  3.87  3.68  3.74  4.46  4.31  3.88  4.16 *  4.43  4.33  3.78  3.97  4.26  4.27  3.75  4.43  4.13  3.64  3.15  3.55 *  4.29  4.09 *  4.45  4.26  4.25  3.77  4.40  4.13  3.52  4.13  4.16  4.05  3.93  4.45  3.58  3.90  3.66  3.46  4.18  3.81  4.35  3.95  4.32  4.47  4.37  3.58  4.29  4.38  4.27  3.90  4.16  4.16  3.97  4.06  4.13  3.79  4.26  4.44 |

^#^The cut-off value is 4.5 Å (includes van der Waals interaction).

All contacts are part of the N-terminal TM-JM helix.

**Table S7:** List of interactions^#^ involving the catalytic triad residues S137, D258 and H286.

| **Source atoms** | **Target atoms** | **Distance (Å)** |
| --- | --- | --- |
| *Monomer A*  Ser 137A N  Ser 137A O  conformation A  Ser 137 A OG  conformation B  Ser 137 A OG  Asp 258A N  Asp 258A OD1  Asp 258A OD2  His 286A N  His 286A ND1  His 286A NE2  His 286A O  *Monomer B*  Ser 137B N  Ser 137B O  conformation A  Ser 137B OG  conformation B  Ser 137B OG  Asp 258B N  Asp 258B OD1  Asp 258B OD2  His 286B N  His 286B ND1  His 286B NE2  His 286B O | Asn 136A ND2  Ile 160A O  Asp 161A O  Ala 163A N  Gly 139A N  Gly 140A N  His 141A N  His 286A NE2  HOH 229S O  MYR 500A O1  HOH 229S O  Gly 255A O  Leu 261A O  Leu 261A N  His 286A ND1  Arg 259A N  Val 260A N  Trp 254A NE1  Leu 261A O  Asp 161A OD2  His 286A ND1  Val 199A O  Asp 258A OD1  Asp 258A OD2  Ser 137A OG  Asn 136A ND2  Asn 136B ND2  Ile 160B O  Asp 161B O  Ala 163B N  Gly 140B N  His 141B N  Gly 139B N  His 286B NE2  HOH 171S O  IPA 504B C1<<<  HOH 171S O  Gly 255B O  Arg 259B N  Leu 261B N  Leu 261B O  Val 260B N  His 286B ND1  Trp 254B NE1  Asp 161B OD2  Leu 261B O  His 286B ND1  Val 199B O  Asp 258B OD2  Asp 258B OD1  Ser 137B OG  Asn 136B ND2 | 3.39  3.20  3.04  3.01  3.17  2.89  2.97  3.04  2.23  3.43  2.52  3.13  3.25  2.93  3.12  3.00  2.75  3.49  3.08  2.63  2.72  3.08  3.12  2.72  3.04  3.25  3.35  3.16  3.05  3.02  2.89  3.02  3.19  2.83  2.62  3.25  2.80  3.08  2.97  2.94  3.29  2.77  3.12  3.43  2.67  3.22  2.72  2.97  2.72  3.12  2.83  3.16 |

^#^The cut-off value for hydrogen bond selection used is ≤ 3.5 Å.

**Table S8:** Residues lining the active site cavity and their interactions with ligands.

| Residue | Interacting ligand |
| --- | --- |
| L27 | - |
| A28 | - |
| V30 | OG(A,B) |
| R31 | 11A |
| E34 | OG(A,B) |
| G70 | - |
| F71 | MYR, IPA503 |
| G72 | - |
| A73 | MYR |
| D74 | - |
| D76 | - |
| N77 | MYR, 11A |
| W78 | - |
| L79 | MYR, 11A |
| R80 | OG(A,B) |
| F81 | - |
| N136 | MYR, IPA503 |
| S137* | MYR, IPA503 |
| M138 | - |
| H141 | - |
| A163 | - |
| F174 | - |
| L173 | - |
| L184 | - |
| V185 | - |
| V186 | - |
| F192 | - |
| L195 | - |
| L196 | - |
| V199 | - |
| F200 | OG(A,B) |
| N203 | OG(A) |
| P204 | OG(A,B) |
| L206 | OG(A,B) |
| L210 | OG(B) |
| L214 | MYR |
| R217 | - |
| A218 | - |
| S222 | - |
| N225 | - |
| F229 | - |
| L232 | - |
| V260 | - |
| L261 | - |
| H286* | MYR, IPA503 |
| M289 | MYR |
| V287 | - |
| V290 | - |

*active site residues

**Table S9:** Michaelis-Menten constants for inhibition of PlaF with decanoic acid (FA C10).

| c(FA C10) [mM] | *K*_m_ [mM]* | *v*_max_ [*U*/mg]* |
| --- | --- | --- |
| 0.0 | 0.17±0.02 | 899.5±37.2 |
| 0.5 | 0.23±0.03 | 916.0±39.4 |
| 1.5 | 0.25±0.03 | 830.9±40.0 |
| 2.5 | 0.34±0.04 | 717.8±35.1 |
| 5.0 | 0.45±0.05 | 512.2±26.6 |
| 7.5 | 0.66±0.04 | 390.8±13.7 |

* Results are mean ± S.D. of three experiments each measured with three samples.

**Table S10:** Average 2D-RMSD_all atom_ of residues 25 to 315 of the structures sampled along MD trajectories.^[a]^

|  | di-PlaF_A_^[b]^ | di-PlaF_B_^[b]^ | PlaF_A_^[c]^ | t-PlaF_A_^[d]^ | PlaF_B_^[c]^ | t-PlaF_B_^[d]^ |
| --- | --- | --- | --- | --- | --- | --- |
| di-PlaF_A_^[b]^ | 3.42 ± 0.59 | 3.96 ± 0.69 | 3.63 ± 0.58 | 3.60 ± 0.61 | 3.94 ± 0.57 | 3.91 ± 0.60 |
| di-PlaF_B_^[b]^ |  | 4.01 ± 0.81 | 4.05 ± 0.68 | 4.05 ± 0.72 | 4.29 ± 0.70 | 4.23 ± 0.71 |
| PlaF_A_^[c]^ |  |  | 3.59 ± 0.60 | 3.71 ± 0.63 | 4.08 ± 0.58 | 4.02 ± 0.64 |
| t-PlaF_A_^[d]^ |  |  |  | 3.58 ± 0.72 | 4.05 ± 0.61 | 3.93 ± 0.65 |
| PlaF_B_ |  |  |  |  | 4.17 ± 0.76 | 4.21 ± 0.62 |
| t-PlaF_B_ |  |  |  |  |  | 3.99 ± 0.80 |

^[a]^ RSMD values in Å, mean ± S.D., were computed in a pair-wise manner for respective structures sampled every ns along the MD trajectories.

^[b]^ PlaF molecules in dimeric form starting from the crystal structure.

^[c]^ PlaF_A_ and PlaF_B_ obtained from the dimeric form by removal of the opposite chain.

^[d]^ PlaF_A_ and PlaF_B_ in the tilted monomeric form.

**SI References**

1.	Martinez-Garcia, E. and V. de Lorenzo, *Engineering multiple genomic deletions in Gram-negative bacteria: analysis of the multi-resistant antibiotic profile of Pseudomonas putida KT2440.* Environ Microbiol, 2011. **13**(10): p. 2702-2716.

2.	Choi, K.-H., et al., *A Tn7-based broad-range bacterial cloning and expression system.* Nature methods, 2005. **2**(6): p. 443-448.

3.	Hall, T., Ibis Biosciences, <http://www.mbio.ncsu.edu/bioedit/bioedit.html>, 2007.

4.	Diekman, J., J.B. Thomson, and C. Djerassi, *Mass spectrometry in structural and stereochemical problems. CLXXIII. The electron impact induced fragmentations and rearrangements of trimethylsilyl esters of w-phenoxyalkanoic acids.* The Journal of Organic Chemistry, 1969. **34**(10): p. 3147-3161.

5.	Lack, N.A., et al., *Characterization of a carbon-carbon hydrolase from Mycobacterium tuberculosis involved in cholesterol metabolism.* J. Biol. Chem., 2010. **285**(1): p. 434-443.

6.	Nardini, M., et al., *Crystal structure of Pseudomonas aeruginosa lipase in the open conformation. The prototype for family I.1 of bacterial lipases.* J. Biol. Chem., 2000. **275**(40): p. 31219-31225.

7.	Habe, H., et al., *Crystal structure of a histidine-tagged serine hydrolase involved in the carbazole degradation (CarC enzyme).* Biochem Biophys Res Commun, 2003. **303**(2): p. 631-9.

8.	Winn, M.D., et al., *Overview of the CCP4 suite and current developments.* Acta Crystallogr D Biol Crystallogr, 2011. **67**(Pt 4): p. 235-42.

9.	Dickson, C.J., et al., *Lipid14: the Amber lipid force field.* J. Chem. Theory Comput., 2014. **10**: p. 865-879.

10.	Tristram-Nagle, S., H.I. Petrache, and J.F. Nagle, *Structure and interactions of fully hydrated dioleoylphosphatidylcholine bilayers.* Biophysical journal, 1998. **75**(2): p. 917-925.

11.	Liu, Y. and J.F. Nagle, *Diffuse scattering provides material parameters and electron density profiles of biomembranes.* Physical Review E, 2004. **69**(4): p. 040901.

12.	Vemparala, S., S. Mehrotra, and H. Balaram, *Role of loop dynamics in thermal stability of mesophilic and thermophilic adenylosuccinate synthetase: a molecular dynamics and normal mode analysis study.* Biochim Biophys Acta, 2011. **1814**(5): p. 630-7.

13.	Deforet, M., D. van Ditmarsch, and J.B. Xavier, *Cell-Size Homeostasis and the Incremental Rule in a Bacterial Pathogen.* Biophysical journal, 2015. **109**(3): p. 521-528.
